# Atypical epigenetic and small RNA control of degenerated transposons and their fragments in clonally reproducing *Spirodela polyrhiza*

**DOI:** 10.1101/2024.04.03.587901

**Authors:** Rodolphe Dombey, Daniel Buendía-Ávila, Verónica Barragán-Borrero, Laura Diezma-Navas, Arturo Ponce-Mañe, José Mario Vargas-Guerrero, Rana Elias, Arturo Marí-Ordóñez

**Affiliations:** Gregor Mendel Institute of Molecular Plant Biology (GMI) of the Austrian Academy of Sciences, Vienna, 1030, Austria; Vienna BioCenter PhD Program, Doctoral School of the University of Vienna and Medical University of Vienna, Austria; Biology School, Costa Rica University, San José, 11501-2060, Costa Rica

## Abstract

A handful of model plants have provided insight into silencing of transposable elements (TEs) through RNA-directed DNA methylation (RdDM). Guided by 24-nt long small-interfering RNAs (siRNAs), this epigenetic regulation installs DNA methylation and histone modifications like H3K9me2, which can be subsequently maintained independently of siRNAs. However, the genome of the clonally propagating duckweed *Spirodela polyrhiza* (*Lemnaceae*) has low levels of DNA methylation, very low expression of RdDM components, and near absence of 24-nt siRNAs. Moreover, some genes encoding RdDM factors, DNA methylation maintenance, and RNA silencing mechanisms are missing from the genome. Here, we investigated the distribution of TEs and their epigenetic marks in the *Spirodela* genome. While abundant degenerated TEs have largely lost DNA methylation and H3K9me2 is low, they remain marked by the heterochromatin associated H3K9me1 and H3K27me1 modifications. By contrast, we found high levels of DNA methylation and H3K9me2 in the relatively few intact TEs which are source of 24-nt siRNAs like RdDM-controlled TEs in other angiosperms. The data suggest that, potentially as adaptation to vegetative propagation, RdDM extent, silencing components, and targets are different from other angiosperms, preferentially focused on potentially intact TEs. It also provides evidence for heterochromatin maintenance independently of DNA methylation in flowering plants. These discoveries highlight the diversity of silencing mechanisms that exist in plants and the importance of using disparate model species to discover these mechanisms.

## Introduction

Transposable elements (TEs), or transposons, are mobile genetic elements which can change location and generate new copies in their host genomes. As a result, TEs populate eukaryotic genomes and can contribute to a significant fraction of their nuclear DNA (Wells and Feschotte 2020). Owing to their mobile nature, TEs have an important role in evolution and genetic innovation, but also pose a threat to the genome integrity (Biémont and Vieira 2006; Feschotte 2008). To control TEs, various mechanisms have evolved to prevent or reduce their mobility.

In eukaryotes, transcriptional gene silencing (TGS) of TEs relies on the deposition of chromatin modifications leading to the formation of compact chromatin states, collectively known as heterochromatin, incompatible with canonical Pol II transcription (Allshire and Madhani 2018). Heterochromatin is generally characterized by repressive epigenetic marks, prominently DNA methylation (5’methylcytosine; 5mC), histone modifications such as histone H3 lysine-9 mono- and dimethylation (H3K9me1/H3K9me2, constitutive heterochromatin) or histone H3 lysine-27 triple methylation H3K27me3 (facultative or developmentally regulated heterochromatin), and the presence of specific histone variants (Henikoff and Smith 2015; Jamge et al. 2023). Upon silencing, TEs accumulate mutations and degenerate into non-autonomous TEs and fragmented remnants or relics (Blumenstiel 2019). Despite their inability to mobilize, degenerated TEs provide alternative DNA and RNA regulatory sequences (e.g., transcription factor binding sites, splicing information, transcriptional start or termination sites) or induce recombination due to their repetitiveness (Ito et al. 2011; Kim and Zilberman 2014; Zervudacki et al. 2018; Sammarco et al. 2022; Ilık et al. 2024). Therefore, maintaining epigenetic control is important even for degenerated TEs, and in consequence, TE abundance and location largely shapes the epigenetic landscape of genomes (Houben et al. 2003; Seymour et al. 2014; Sigman and Slotkin 2016; Wyler et al. 2020; Klein and Anderson 2022).

Another conserved mechanism to suppress TEs and maintain their epigenetic silencing depends on ∼18-30-nt long small RNAs (sRNAs) and members of the ARGONAUTE (AGO) protein family. Loaded into AGOs, TGS-associated sRNAs guide chromatin modifying complexes to TEs through sequence complementarity (Moazed 2009; Malone and Hannon 2009). In flowering plants (angiosperms) TE silencing is mostly associated with DNA methylation and H3K9me2, and the mechanism by which sRNAs guide chromatin modifications is known as RNA-directed DNA methylation (RdDM). Best studied in *Arabidopsis thaliana,* during RdDM, H3K9me2 and DNA methylation recruit the plant-specific RNA Polymerases IV and V (Pol IV, V). Pol IV transcripts are processed into 24-nt siRNAs which, loaded into members of the AGO4/6 clade, target Pol V transcripts to guide the sequence-specific deposition of DNA methylation in all cytosine contexts (CG, CHG and CHH, where H is any nucleotide but G). Once stablished, DNA methylation can reinforce the deposition of H3K9me2 through the action of SUPRESSOR OF VARIEGATION 3-9 HOMOLOGUE (SUVH) family H3K9 methyltransferases, which can read DNA methylation. In turn, CHROMO-METHYL TRANSFERASE 3 and 2 (CMT3, CMT2) can bind to H3K9me2 to mediate CHG and CHH methylation respectively. This results in a self-reinforcing loop between non-CG methylation and H3K9me2 independent of siRNAs. Symmetrical CG methylation is maintained across cell divisions through the action of METHYL-TRANSFERASE 1 (MET1), recruited to hemi-methylated CG sites after DNA replication by VARIANT IN METHYLATION (VIM) proteins (reviewed in (Erdmann and Picard 2020; Tirot et al. 2021)). Loss of RdDM, 5mC or H3K9me2 results in the transcriptional reactivation of TEs (Mirouze et al. 2009; Tsukahara et al. 2010; Stroud et al. 2014; He et al. 2021; Osakabe et al. 2021). Therefore, DNA methylation, together with H3K9me2 and in cooperation with sRNAs, keep TEs silenced.

Loss of TE control in somatic tissue is likely less deleterious as long as it does not affect the germline or, in plants, stem cells in the meristems or gamete precursor cells giving rise to the next generation, in which stability of the genome is highly relevant. In accordance, small RNA-guided TGS pathways are prominently active in reproductive tissues and silence active TEs encountered during fertilization (Bourc’his and Voinnet 2010; Marí-Ordóñez et al. 2013; Parhad and Theurkauf 2019). However, unlike in metazoans, where such pathways are predominantly active in the gonads, in most flowering plants RdDM is not constrained to reproductive organs. 24-nt siRNAs are also present in vegetative, somatic organs such as leaves or roots across all developmental stages (Havecker et al. 2010; Patel et al. 2021; Zhou et al. 2022). With interesting exceptions: the aquatic dicot carnivorous plant *Utricularia gibba* (L.), and the monocot *Spirodela polyrhiza* (L.) Scheid. Both contain much less 24-nt siRNAs and display reduced genome-wide levels of DNA methylation in vegetative tissues compared to other angiosperms (Michael et al. 2017; Fourounjian et al. 2019; Cervantes-Pérez et al. 2021). While in the *U. gibba* such features can be a consequence of low TE content as repeats represent less than 3% of genome (Ibarra-Laclette et al. 2013), the TE content of the *Spirodela* genome is similar to that of *Arabidopsis* (∼20%) (Wang et al. 2014). Therefore, reduced genomic TE load is unlikely to be the cause behind the low levels of DNA methylation and small RNAs in *Spirodela*. Furthermore, in *Spirodela*, several sRNA silencing components including some involved in RdDM have been shown to be missing, low or not expressed. In addition, CMT2, which maintains CHH methylation, is lost in duckweeds (An et al. 2019; Ernst et al. 2023; Harkess et al. 2024).

The *Spirodela* genus is the most basal of the *Lemnaceae* family (order Alismatales), a monophyletic group of free-floating freshwater monocot plants, commonly known as duckweeds, composed of 5 genera (*Spirodela*, *Landoltia*, *Lemna*, *Wolfiella* and *Wolffia*) (Bog et al. 2019; Tippery et al. 2021). Comprised of a leaf-like structure known as a frond, ranging from ∼2 cm to 1 mm, duckweeds display an evolutionary reduction and simplification of their body plan compared to most angiosperms, including vestigiality and loss of organs such as roots and vasculature (Landolt 1986; Lemon and Posluszny 2000; Lam and Michael 2022; Ware et al. 2023). Although flowering and seed production has been reported, it is rare in most species (Landolt 1986; Pieterse 2013; Fourounjian et al. 2021). Duckweeds mostly propagate through rapid clonal reproduction (doubling of individuals ∼1-5 days), known as vegetative budding: new fronds originate from budding pockets where meristems containing stem cells are located (Landolt 1986; Ziegler et al. 2015). Mother fronds (MF) continuously give rise to daughter fronds (DF), which carry grand-daughter fronds (GDF) in their budding pockets even before detaching from the mother frond (Landolt 1986; Lemon and Posluszny 2000). This extremely efficient amplification of somatic tissue, omitting regular sexual propagation, and the free-floating way of living, make *Spirodela* a unique model to explore if the differences in epigenetic parameters are connected with these differences in lifestyle compared to other angiosperms. Thus, the aim of this study is to investigate the TE silencing landscape in *Spirodela polyrhiza* by profiling its genetic composition of silencing pathways, small RNAs and epigenetic modifications commonly associated with TE control in angiosperms.

## Results

### Genome sequencing of *S. polyrhiza* #9509 from the Landolt collection

The genomes of several *Spirodela polyrhiza* accessions (clones) have been sequenced (Wang et al. 2014; Michael et al. 2017; Xu et al. 2019), including a high-resolution reference genome assembly of its 20 chromosomes from the *S. polyrhiza* 9509 and 7498 clones (Sp9509, Sp7498) (Michael et al. 2017; Hoang et al. 2018; Harkess et al. 2021). For our study, we retrieved Sp9509 from the Landolt duckweed collection (Zurich, Switzerland), now managed at the Institute of Agriculture Biology and Biotechnology (IABB, CNR, Milan, Italy) (Morello et al. 2024). We performed 88× coverage short-read Illumina genome sequencing to account for potential genetic differences due to independent clonal propagation over thousands generations between this accession and that of the Rutgers Duckweed Stock Cooperative (RDSC, Rutgers University, NJ, USA) used to generate the reference genome (Hoang et al. 2018). The Landolt Sp9509 genome showed ∼163k indels and ∼512k SNPs (∼0.4% of the *Spirodela* genome) compared to the Sp9509 reference (**Supplemental Fig. S1**). Given our special interest on TEs, we performed a *de novo* TE annotation using a combination of EDTA and RepeatModeler2 to identify repeats that were then further classified using DeepTE (Ou et al. 2019; Flynn et al. 2020; Yan et al. 2020).

To explore the TE silencing landscape during *Spirodela* asexual clonal reproduction, we used *Arabidopsis* as a comparative system to investigate *Spirodela* TE-derived sRNAs, TE epigenetic marks and the genetic composition and expression of silencing pathways. Although TE regulatory mechanisms might differ between the two species given their evolutionary distance (**Supplemental Fig. S2**), we choose *Arabidopsis* as its silencing pathways and epigenetic mechanisms are well characterized by molecular and genetic means. Furthermore, the two species display similar genome size and TE content (Fig. 1A).

**Figure 1:**
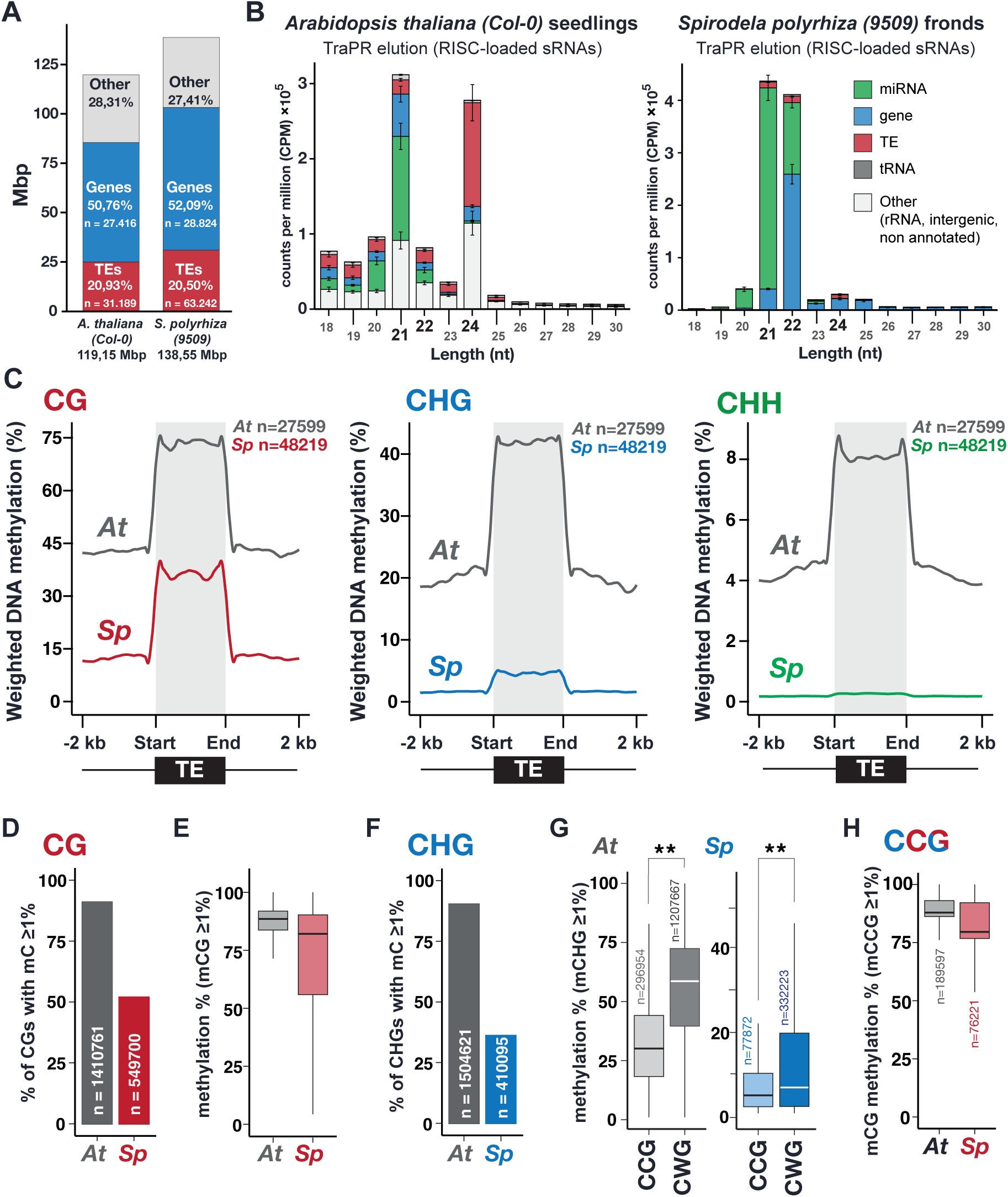
*Spirodela polyrhiza* general developmental, small RNA and DNA methylation patterns. **(A)** Genome size, number and genomic occupancy of genes and TEs in *Spirodela* and *Arabidopsis*. **(B)** Size distribution and genomic feature annotation of TraPR-purified small RNA from *Arabidopsis* and *Spirodela*. Error bars represent standard deviation of the mean of three biological replicates. **(C)** Metaplots of averaged weighted DNA methylation in all three contexts (CG, CHG and CHH, where H is any but G) over transposons (TEs) (≥100 bp) and 2 kb flanking regions in *Arabidopsis* and *Spirodela*. **(D)** number and percentage of transposon CG sites displaying methylation levels equal or above 1% in *Arabidopsis* and *Spirodela*. **(E)** DNA methylation levels at CG sites from (D). **(F)** number and percentage of transposon CHG sites displaying methylation levels equal or above 1% in *Arabidopsis* and *Spirodela*. **(G)** DNA methylation levels at CCG and CWG in *Arabidopsis* and *Spirodela*. **(H)** Inner 5mC (CmCG) DNA methylation levels at CCG sites with outer 5mC (mCCG) equal or higher than 1% in *Arabidopsis* and *Spirodela*. (**) indicates p-value <2.22 × 10^-16^ (Wilcoxon rank sum test). In all boxplots: median is indicated by solid bar; the boxes extend from the first to the third quartile and whiskers stretch to the furthest value within 1.5 times the interquartile range. At: *Arabidopsis*, Sp: *Spirodela*.

### AGO-loaded small RNA and DNA methylation patterns in *Spirodela* imply low RdDM activity in *Spirodela*

Given that 24-nt sRNAs are considered one of the hallmarks of RdDM, we first compared the sRNA profiles of *Spirodela* confluent cultures (including fronds in all developmental stages) to those of 10 day-old *Arabidopsis* seedlings as the closest equivalent developmental stage. As sRNAs on the 21-to-24-nt range were very low abundant in profiles generated from total RNA (**Supplemental Fig. S3**), we enriched our small RNA libraries by investigating AGO-loaded small RNAs using TraPR (Grentzinger et al. 2020). In contrast to *Arabidopsis*, where AGO-loaded TE-derived 24-nt siRNAs are highly abundant, *Spirodela* TraPR profile displays very low levels of 24-nt siRNAs and it is dominated by 21-22-nt sRNAs (Fig. 1B**, Supplemental Fig. S3**), predominantly contributed by regulatory microRNAs (miRNA). The lack of 24-nt siRNAs in *Spirodela* TraPR profile is more pronounced than their loss in *Arabidopsis* RdDM siRNA biogenesis defective Pol IV mutant (*nrpd1*) (**Supplemental Fig. S3**) (Teixeira et al. 2009; Nuthikattu et al. 2013; Creasey et al. 2014) Hence, low levels of AGO-loaded 24-nt siRNAs in *Spirodela* are compatible with the reduced or lack of RdDM activity at TEs during its asexual clonal growth.

Given the connection between sRNAs and DNA methylation, we next investigated *Spirodela* DNA methylation patterns. Methylome analysis by Enzymatic Methyl-seq (EM-seq) (Feng et al. 2020) showed similar results to those previously published using bisulfite sequencing (Michael et al. 2017). TE 5mC in all contexts is lower in *Spirodela* compared to *Arabidopsis*, with very low mCHG levels and hardly any mCHH (Fig. 1C). This is in agreement with the partial dependency of mCHH on 24-nt siRNAs. In addition, *Spirodela* lacks gene body CG methylation (gbM) (Michael et al. 2017; Harkess et al. 2024) (**Supplemental Fig. S4A**). Although the role of gbM is unclear, it generally associates with highly expressed genes. In contrast, mCG near the transcription start site (TSS) is associated with gene silencing (Niederhuth et al. 2016; Bewick and Schmitz 2017). Although expressed genes do not display gbM, like in *Arabidopsis*, mCG around the TSS is higher in non-expressed genes of *Spirodela* (**Supplemental Fig. S4B,C**), indicating that mCG in gene regulatory sequences influences transcription also in *Spirodela*.

### CG and CHG methylation signatures are compatible with functional maintenance of DNA methylation independently of small RNAs

Once established, DNA methylation can be further maintained independently of small RNAs through several pathways depending on the sequence context (Mathieu et al. 2007; Cokus et al. 2008; Reinders et al. 2009) In *Spirodela*, the ratio of mCG sites at TEs (51,9%) is nearly half of that in *Arabidopsis* (90,9%) (Fig. 1D). However, at CG sites with 5mC levels equal or higher than 1%, mCG is similarly stable as in *Arabidopsis*, displaying median methylation levels above 80% (Fig. 1E), although a higher variation was observed for *Spirodela* than *Arabidopsis*. Hence, the reduced global mCG level at TEs in *Spirodela* is the result of less individual CG sites being methylated.

As for mCG, the number of mCHG sites at TEs is much lower in *Spirodela* (36,1%) than in *Arabidopsis* (90,3%) (Fig. 1F). CHG sites can be further divided into CCG and CWG (CAG/CTG) sub-contexts. In *Arabidopsis*, mCWG is higher than mCCG (Fig. 1G) due to the combined differential affinities of SUVH4/5/6 and CMT3 to read and methylate individual cytosine sequence contexts (Cokus et al. 2008; Gouil and Baulcombe 2016; Li et al. 2018; Fang et al. 2022). This applies also to *Spirodela*, although methylation in both sub-contexts is lower than in *Arabidopsis* (Fig. 1G). Furthermore, maintenance in the mCCG sub-context relies additionally on MET1. Being both CG and CHG contexts, methylated CCG sites are generally observed as mCmCG due to the outer cytosine methylation (mCCG) being largely dependent on the internal CG methylation (CmCG) mediated by MET1 (Stroud et al. 2013; Yaari et al. 2015; Gouil and Baulcombe 2016; Li et al. 2018). Consequently, mCCG sites in *Arabidopsis* display a high internal CG methylation level, a pattern also observed in *Spirodela* (Fig. 1H). The similar mCHG patterns between the two species indicate that the responsible enzymes, SUVHs and ZMET, the CMT3 homolog in monocots (Bewick et al. 2017) probably share the same sequence preference albeit their activity seems to be lower than in *Arabidopsis*. Nonetheless, given that CMT3/ZMET has been shown to be involved in gbM (Niederhuth et al. 2016; Bewick et al. 2016; Wendte et al. 2019), lower mCHG and lack of gbM in *Spirodela* might be related.

Altogether, our analysis indicates that mCG and mCHG maintenance mechanisms in *Spirodela* might operate in a similar fashion as in *Arabidopsis*. However, while low mCG is a consequence of reduced number of CG sites being methylated, low mCHG appears to be the combined result of lower number of targeted CHG sites and decreased maintenance activity.

### Genetic composition and expression of silencing pathways partially account for sRNAs and DNA methylation features in *Spirodela*

In addition to low signatures of RdDM in *Spirodela*, coding potential for several of its components have been shown to be missing (An et al. 2019; Ernst et al. 2023; Harkess et al. 2024). To gain further insights on why levels of 24-nt siRNA and DNA methylation are reduced, we set to identify and examine the expression of known core components of angiosperm silencing pathways in *S. polyrhiza* (Fig. 2**, Supplemental Fig. S5**). The identification of orthologues and paralogues in the genome of *S. polyrhiza* 9509 was based on the presence of conserved functional domains. Annotations were further curated based on phylogeny trees built with proteins from several angiosperms and the prediction of known key domains and their organization (**Supplemental Fig. S6-S18**). For gene expression, we performed short-read transcriptome analysis in *Spirodela* and compared it to that of recently published *Arabidopsis* seedlings (Li et al. 2023) (Fig. 2). We further performed long-read cDNA-seq with PacBio full-length transcript Isoform-Sequencing (Iso-Seq) technology in *Spirodela* to validate and eventually improve gene model annotations.

**Figure 2:**
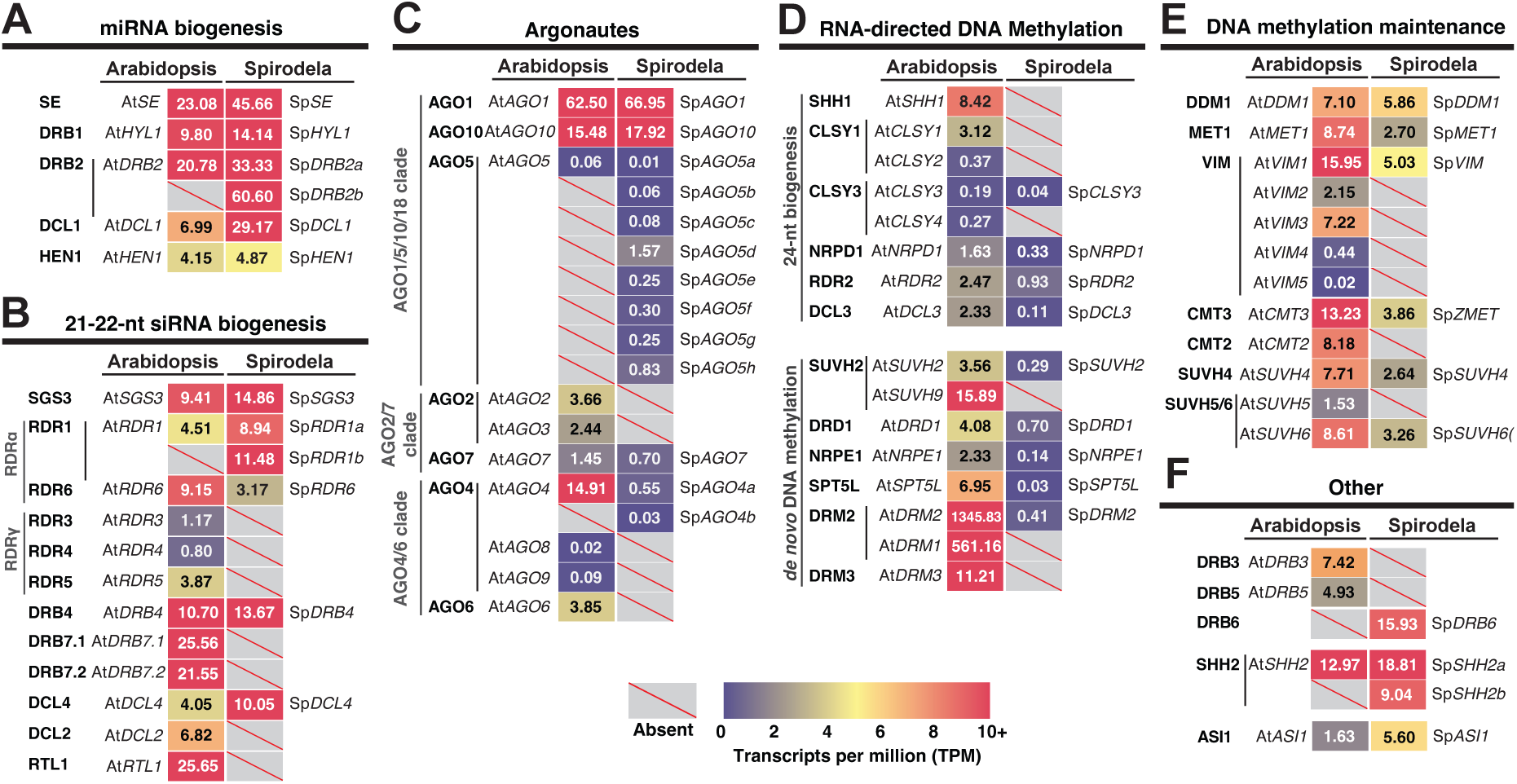
Genetic composition and expression of silencing pathways in *Spirodela*. **(A-F)** Expression in transcripts per million (TPM) of silencing components in *Arabidopsis* and their identified orthologues and paralogues in the *Spirodela polyrhiza* #9509 (Sp9509) genome. Gene IDs for both *Arabidopsis* and *Spirodela* can be found in Supplemental Figure S5.

As expected from the prevalence of miRNAs in *Spirodela* TraPR analysis, conserved plant miRNA biogenesis factors (Zhan and Meyers 2023) are expressed at similar levels as those in *Arabidopsis* (Fig. 2A). Likewise, genes required for the production of regulatory 21-nt siRNA trans-acting (ta)siRNA (Yoshikawa et al. 2005; Liu et al. 2020): SUPRESSOR OF GENE SILENCING 3 (SGS3), RNA-DEPENDENT RNA POLYMERASE 6 (RDR6) and DICER-LIKE 4 (DCL4), are present and expressed (Fig. 2B). *Spirodela* lacks many components, effectors and regulators of 21-22-nt siRNA pathways, such as RNASE THREE-LIKE 1 (RTL1), DOUBLE-STRANDED RNA BIDING PROTEIN 7 (DRB7) (Shamandi et al. 2015; Tschopp et al. 2017; Montavon et al. 2017), and orthologues of the RDRγ clade highly conserved in plants and fungi (Zong et al. 2009; Willmann et al. 2011) (Fig. 2B**; Supplemental Fig. S6, S7**). Gene absences also include the antiviral factors DICER-LIKE 2 (DCL2) and ARGONAUTE 2 (AGO2) (Deleris et al. 2006; Garcia-Ruiz et al. 2010; Harvey et al. 2011; Carbonell et al. 2012) (Fig. 2B**,C****; Supplemental Fig. S8, S9**). In contrast, two RDR1 and eight tandemly repeated copies of AGO5 are present (Fig. 2B**,C****; Supplemental** Fig. S7, S9, S10). Both have been implicated in virus exclusion and TE silencing in meristems in *Arabidopsis* (Yu et al. 2003; Brosseau and Moffett 2015; Incarbone et al. 2023; Bradamante et al. 2023) and, in monocots, AGO5 participates in anther development and TE silencing during male gametogenesis (Zhai et al. 2015; Fei et al. 2016; Lee et al. 2021).

Regarding the RdDM pathway, *Spirodela* contains the minimal set of its components, either with no or with a reduced number of paralogues compared to *Arabidopsis* (Fig. 2C, D). In *Spirodela*, SAWADEE HOMEODOMAIN HOMOLGUE 1 (SHH1), a dual reader for repressive H3K9me2 and unmethylated H3K4(me0), and the chromatin remodeler CLASSY 1/2 (CLSY1/2) are absent (Fig. 2D**; Supplemental Fig. S11, S12**). However, a single orthologue of CLSY3 was found in *Spirodela* (Fig. 2D**; Supplemental Fig. S12**). In *Arabidopsis*, the SHH1-CLSY1/2 module mainly recruits Pol IV to short TEs within gene-rich regions, while CLSY3 and its paralogue CLSY4 recruit Pol IV to TE-rich pericentromeric regions in a SHH1-idependent but DNA methylation-dependent manner (Law et al. 2011; 2013; Zhou et al. 2018; 2022). However, *Spirodela* has two SHH2 orthologues (Fig. 2**; Supplemental Fig. S11**). Part of a nucleosome positioning complex during transcription initiation in *Arabidopsis* (Diego-Martin et al. 2022), in maize, SHH2 binds preferentially to H3K9me1 *in vitro* and interacts *in vivo* with both Pol IV and V in maize, possibly connecting RdDM to H3K9me1 installation (Haag et al. 2014; Wang et al. 2021).

Similarly, no AGO6 orthologue was found, yet two AGO4, both required non-redundantly to maintain DNA methylation at most RdDM target loci in *Arabidopsis* (Duan et al. 2014) (Fig. 2D**; Supplemental Fig. S9**). A single SUVH2 orthologue is present in *Spirodela* (Fig. 2D**, Supplemental Fig. S13**), which together with its paralogue SUVH9 redundantly recruit Pol V to RdDM targets by binding to methylated DNA in *Arabidopsis* (Johnson et al. 2008; 2014; Liu et al. 2014). Regarding the RdDM-associated DNA methyltransferases, DOMAIN REARRANGED METHYLTRASFERASE 2 (DRM2) but no DRM3, a conserved catalytically inactive paralogue that contributes to DRM2 activity (Chan et al. 2004; Wierzbicki et al. 2009; Zhong et al. 2015; Liu et al. 2018; Fang et al. 2021), orthologues were found (Fig. 2D**, Supplemental Fig. S14**). Additionally, although NRPE1 (Pol V) and its partner SUPPRESSOR OF TY INSERTION-LIKE (SPT5L) (a paralogue of the general transcription elongation factor SPT5) are conserved in *Spirodela*; we noticed that both contain fewer tryptophan/glycine (GW/WG) repeats than those of other angiosperms (Fig. 2D**; Supplemental Fig. S15, S16**) (El-Shami et al. 2007; Bies-Etheve et al. 2009; Rowley et al. 2011; Huang et al. 2015; Trujillo et al. 2016). GW/WG repeats, or AGO-hooks, allow AGO4/6 binding to Pol V to scan its nascent transcripts for positive siRNA matches to recruit DRM2. These, together with NRPD1, RDR2, DCL3, and the rest of conserved RdDM components, might suffice for a functional pathway in *Spirodela* (Fig. 2**; Supplemental** Fig. S7, S8, S12, S15). Nonetheless, their low expression levels (Fig. 2C**,D**) rationalize the low abundance of AGO-bound 24-nt siRNAs and associated CHH methylation during clonal propagation in *Spirodela* (Fig. 1B**,C**).

Lastly, we inspected DNA methylation maintenance pathways. As predicted from the patterns of mCG and mCHG (Fig. 1D**-H**), MET1, VIM (ORTH), CMT3/ZMET, SUVH4, SUVH5/6 and DECREASE IN DNA METHYLATION 1 (DDM1) orthologues were found present and expressed in *Spirodela*, albeit at a lower degree than in *Arabidopsis* (Fig. 2E**; Supplemental** Fig. S12, S13, S14, S17). A single orthologue of the SUVH5/6 orthogroup was found in *Spirodela* (Fig. 2E**; Supplemental Fig. S13**). In *Arabidopsis*, SUVH5 and SUVH6 display different 5mC sub-context binding affinities as well as genomic targets (Johnson et al. 2007; Rajakumara et al. 2011; Li et al. 2018; Zhang et al. 2023). Absence of either could have implications for the DNA methylation landscape if their functions are conserved outside of *Arabidopsis*. *Spirodela* SUVH5/6 othologue contains a conserved motif present in *Arabidopsis* SUVH6, but absent in SUVH5, required for interaction with ANTI-SILENCING 1 (ASI1) to regulate silencing at intronic TEs and expression of their host genes (Saze et al. 2013; Wang et al. 2013; Duan et al. 2017; Zhang et al. 2023). As an ASI1 orthologue was also found in *Spirodela* (Fig. 2F**; Supplemental Fig. S18**), we named the *Spirodela* SUVH5/6 orthologue as spSUVH6(5) until its sequence binding preference is determined. In addition, with regards to mCHH maintenance, CMT2, required to maintain mCHH together with SUVH4 (Stroud et al. 2014), is missing like in maize (Bewick et al. 2017), as previously reported (Harkess et al. 2024) (Fig. 2E**, Supplemental Fig. S13, S14**).

Thus, reduced 24-nt siRNA and mCHH levels in *Spirodela* can be explained by the combined reduction in RdDM composition and expression in addition to the absence of CMT2. However, it does not account for the global decrease in mCG and mCHG as their maintenance pathways seem intact and expressed in *Spirodela*.

### Few transposable elements from different superfamilies display high levels of DNA methylation in *Spirodela*

To further gain insight into the patterns of TE DNA methylation, we examined its distribution along the *Spirodela* genome. To exclude any confounding effects of gbM, we calculated mean DNA methylation levels exclusively on TEs and their density in 100-kb windows. At gene-rich regions, DNA methylation is very scarce (Fig. 3A**; Supplemental Fig. S19A**). As in *Arabidopsis*, *Spirodela* DNA methylation is concentrated at TE-dense regions (presumably centromeric and pericentromeric domains). However, not all TE-dense regions are highly methylated (**Supplemental Fig. S19B**). Hence, we investigated if local DNA methylation levels correlate with TE density as in other angiosperms (Zhang et al. 2006; Regulski et al. 2013; Seymour et al. 2014; Wyler et al. 2020). At equal TE-content, DNA methylation is lower in all contexts in *Spirodela* compared to *Arabidopsis*, especially at CHG and CHH (Fig. 3B**; Supplemental Fig. S20A**). Moreover, at the CG context, *Spirodela* displays a broad variation range for a given TE content: even at TE densities above 50% or higher, average mCG still spans between 0 and 75% (Fig. 3B).

**Figure 3:**
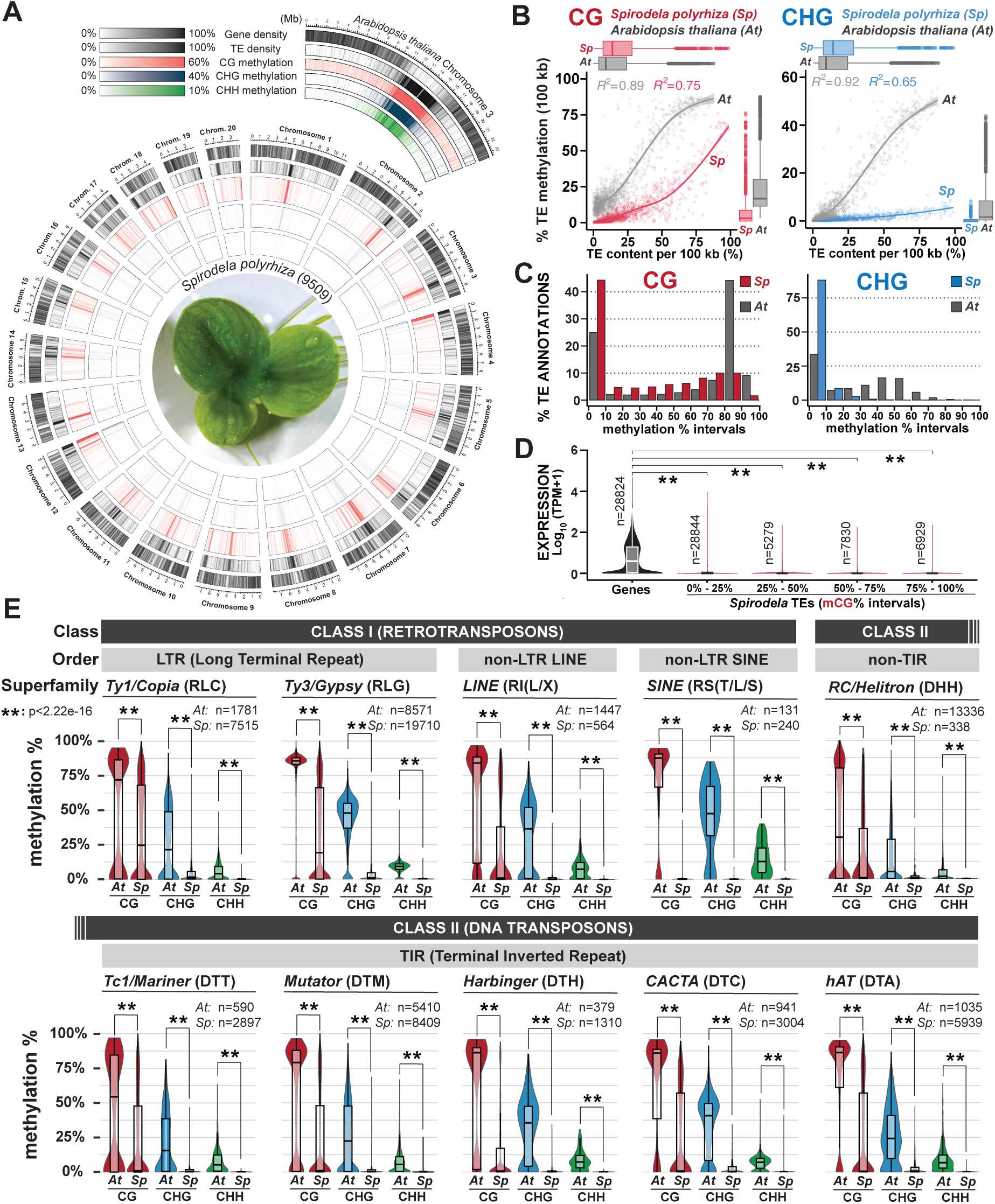
Distribution of DNA methylation in *Spirodela* chromosomes and transposable elements. **(A)** Circos plot of genome-wide distribution of DNA methylation densities by chromosome in *Spirodela*. From outside to inside: gene density, TE density, CG, CHG and CHH methylation. Mean DNA methylation, gene and TE densities values within the indicated ranges are displayed for 100 kbp intervals in 50 kbp sliding windows. *Arabidopsis* Chromosome 3 is shown for comparison. **(B)** Scatter plot and best fit line of average TE CG and CHG DNA methylation against TE content per 100 kb genomic bins in *Arabidopsis* and *Spirodela*. Colored shadows indicate the standard error of the mean. **(C)** Bar plot of the distribution (%) of TE annotations (≥100 bp) within 10% incremental intervals of CG and CHG methylation in *Arabidopsis* and *Spirodela*. **(D)** Analysis of *Spirodela* TE expression within 25% incremental intervals of CG methylation. Global gene expression values are shown for comparison. **(E)** Distribution of CG, CHG and CHH methylation across the main TE superfamilies in *Arabidopsis* and *Spirodela*. The number of TE annotations belonging to each superfamily by species is indicated (n). (**) indicates p-value <2.22 × 10^-16^ (Wilcoxon rank sum test). In boxplots, median is indicated by solid bar, the boxes extend from the first to the third quartile and whiskers stretch to the furthest value within 1.5 times the interquartile range. At: *Arabidopsis*, Sp: *Spirodela*.

To address the origin of such variation, we investigated the DNA methylation levels of individual TEs. This revealed that the majority of TEs in *Spirodela* display low or very low levels of DNA methylation in all contexts, especially CHG and CHH (Fig. 3C**; Supplemental Fig. S20B,C**). Nonetheless, despite this substantially reduced DNA methylation, increased expression at TEs was not observed (Fig. 3D**; Supplemental Fig. S20D**). TEs with high or low DNA methylation levels did not belong to any specific TE class or superfamily. For all the main plant TE superfamilies (Wicker et al. 2007) (TE classification and codes used here are summarized in **Supplemental Fig. S21**), DNA methylation was nearly absent except for a few elements (Fig. 3E). Thus, low genome-wide methylation levels in *Spirodela* result from only a few methylated TEs, although these can display high mCG levels similar to those found in *Arabidopsis* (Fig. 3C**,E**).

### Potentially intact transposons are the conspicuous targets of DNA methylation in *Spirodela*

Although TE location and density influences their epigenetic regulation (Sigman and Slotkin 2016), in *Arabidopsis* and tomato DNA methylation of longer, evolutionary younger and therefore potentially autonomous TE insertions are differentially regulated than often fragmented TE relics resulting from the accumulation of mutations and truncations over evolutionary times following their silencing (Sigman and Slotkin 2016; Blumenstiel 2019; Zattera and Bruschi 2022). On one hand, DNA methylation in long TEs is maintained by a combination of RdDM acting on their boundaries (mainly in young active TEs) and CMT activity along their body. On the other hand, short, degenerated TE remnants do not need CMT to maintain their methylation (Zhong et al. 2012; Zemach et al. 2013; Wang and Baulcombe 2020). To explore if TE length influences DNA methylation in *Spirodela*, we sorted TEs into short TE fragments (100 to 500 bp), intermediate TEs (0.5 to 4 kb) and long TEs (above 4 kb). This revealed that only long TEs are heavily methylated at CG and, to a lesser extent, in CHG. Intermediate TEs display a wide range of mCG levels, whereas TE fragments are minimally or not methylated (Fig. 4A).

**Figure 4:**
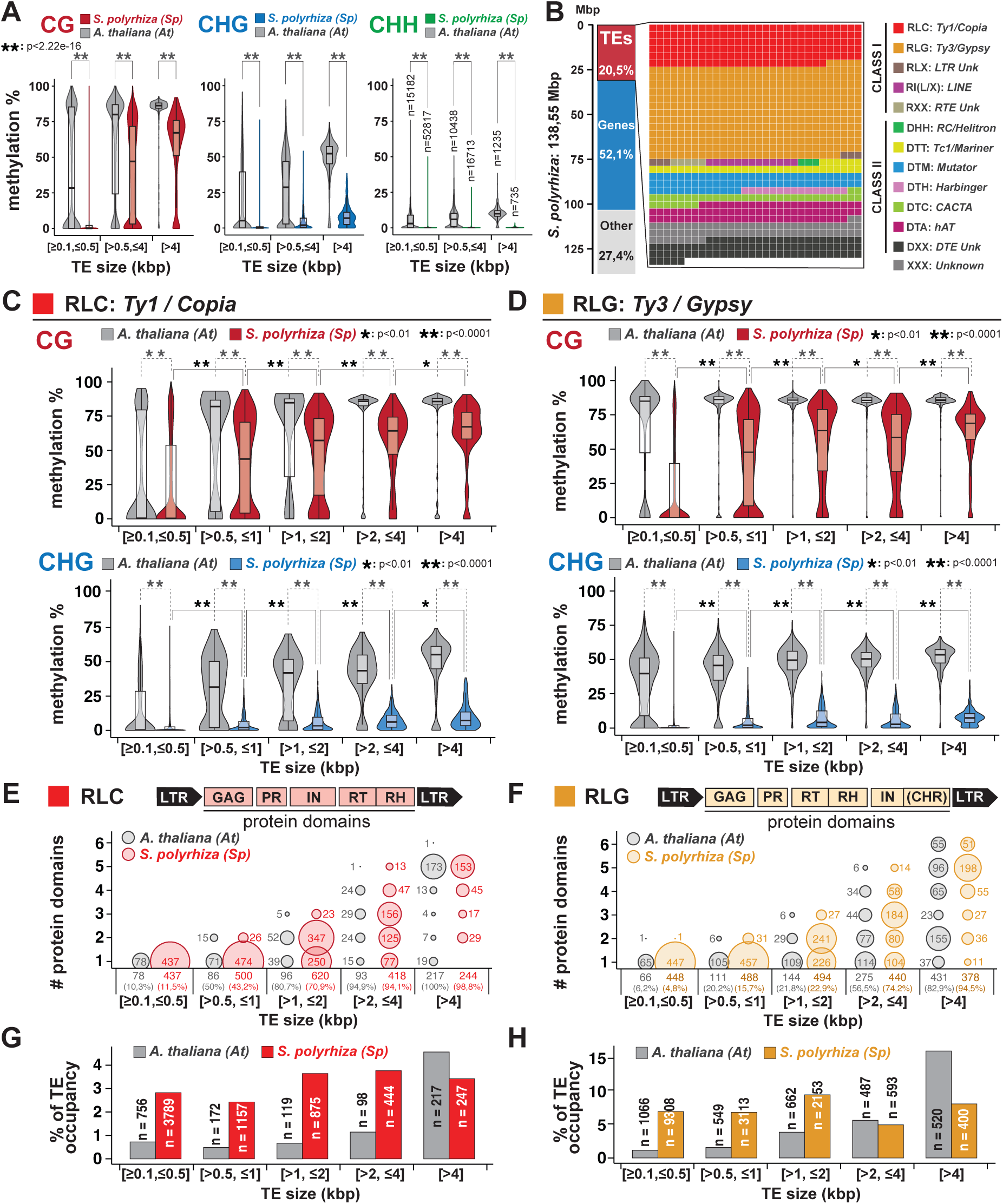
Length-dependent TE DNA methylation in *Spirodela polyrhiza*. (A), CHG and CHH methylation levels (%) across all TEs (≥100 bp) in *Arabidopsis* and *Spirodela* split by size into short (≥100 bp, <500 bp), intermediate (>500 bp, ≤ 4 kbp) and long (> 4kbp). n values for each size category in each genome is indicated within the plot for mCHH (B) Waffle chart of the relative contribution of different TE superfamilies over the total genomic space occupied by TEs in *Spirodela*. (C-D) CG and CHG methylation levels (%) between different length range groups for RLC: *Ty1/Copia* (C) and RLG: *Ty3/Gypsy* (D) in *Arabidopsis* and *Spirodela*. n values are indicated in panels G-H (E-F) Presence and number of RCL (E) and RLG (F) TE protein domains, determined by TEsorter, across TE annotations of varying sizes in *Arabidopsis* and *Spirodela*. Bubble plots depict domain count distributions for TEs with at least one domain. Numbers above TE size category denotes TE annotations with at least one domain, with percentages shown in brackets relative to total TE annotations per size category (G-H). Bubble size is relative to the number of TEs (indicated within or next to the bubble). Schematic representation of RLC and RLG protein and domain structure is included on top of each plot. GAG: Nucleocapsid; PR: Protease; IN: Integrase; RT: Reverse transcriptase; RH: RNase H; CHR: Chromodomain, present only in the Chromoviridae genus of RLG. (G-H) Number of TE annotations and relative contribution to total genomic TE load (TE occupancy %) at each size category for RLC (G) and RLG (H). (*) indicates p-value <0.01 (**) indicates p-value <0.0001 (Wilcoxon rank sum test). In all boxplots: median is indicated by solid bar; the boxes extend from the first to the third quartile and whiskers stretch to the furthest value within 1.5 times the interquartile range. At: *Arabidopsis*, Sp: *Spirodela*.

To further explore the connection between TE length and DNA methylation we focused on the two superfamilies of long terminal repeat (LTR) retroelements (RTEs): *Ty1/Copia* (RLC) and *Ty3/Gypsy* (RLG). LTR-RTEs are the most abundant class of TEs in the *Spirodela* genome, collectively representing more than 50% of the TE fraction of the genome and largely contributing as well to that of *Arabidopsis* (Fig. 4B**; Supplemental Fig. S22**). In addition, the genetic structure and domain organization of complete functional elements are well known (Kumar and Bennetzen 1999; Gorinšek et al. 2004; Neumann et al. 2019). Thus, LTR-RTEs offer a good case study to investigate any link between DNA methylation and TE length, using the latter as a proxy for their completeness and potential autonomy for transposition under the assumption that degenerated and fragmented shorter TEs have lost some or all the required domains to mobilize autonomously (non-autonomous elements) (Blumenstiel 2019). For better resolution, we split TE sizes into further size categories in the intermediary range between 0.5 and 4 kb. Within each size bin, *Spirodela* DNA methylation is lower than in *Arabidopsis*. However, in *Spirodela*, average mCG and mCHG gradually increase with TE size, although mCHG does not achieve levels comparable to those of *Arabidopsis* (Fig. 4C**,D****; Supplemental Fig. S23A,B**). The proportion of TEs displaying high CG methylation rises across TE-size groups. While very few TE-fragments below 500 bp are highly methylated, the majority of long TEs (>4 kb) have mCG above 60% (Fig. 4C**,D**). At the same time, the fraction of TEs with at least one intact cognate protein domain and the number of domains identified per TE annotation increase with each size category, with the majority of long LTR-RTEs containing all domains associated with each superfamily (Fig. 4E**,F**).

Therefore, in *Spirodela*, most potentially intact TEs display high levels of DNA methylation, which decreased as they degenerated and decayed. Given that long intact LTR-RTEs represent a minority of *Spirodela* TEs (**Supplemental Fig. S23C,D**), their low abundance helps explain the few TE numbers with high DNA methylation (Fig. 3C**,E**). In addition, the proportion of hypomethylated shorter LTR-RTE fragments relative to methylated intact TEs is higher in *Spirodela* than in *Arabidopsis* (Fig. 4G**,H****; Supplemental Fig. S23G,H**), resulting in the global reduction of TE methylation (Fig. 1E, Fig. 3A**,B**).

### Levels of H3K9me2, but not H3K9me1, are reduced in *Spirodela*

Considering the association between DNA and H3K9 methylation in plants, we explored how far DNA methylation in *Spirodela* would impact the histone methylation landscape. H3K9me2 and H3K9me1 are commonly linked with heterochromatin formation at silenced TEs (Jackson et al. 2004; Fuchs et al. 2006; Stroud et al. 2014; Jamge et al. 2023). Western blot analysis revealed that the H3K9me2 level in *Spirodela* is lower than in *Arabidopsis*, whereas H3K9me1 displayed similar levels (Fig. 5A). This was unexpected as we anticipated in *Spirodela* a decrease of both marks, given their connection with DNA methylation and reduction upon loss of non-CG methylation in *Arabidopsis* (Stroud et al. 2014; Choi et al. 2021). To corroborate the observation, we investigated histone post-translational modifications (PTMs) by mass-spectrometry (Johnson et al. 2004; Scheid et al. 2022), which confirmed an approximately 8-10-times lower level of H3K9me2 but similar H3K9me1 levels in *Spirodela* relative to *Arabidopsis* (**Supplemental Fig. S24**).

**Figure 5:**
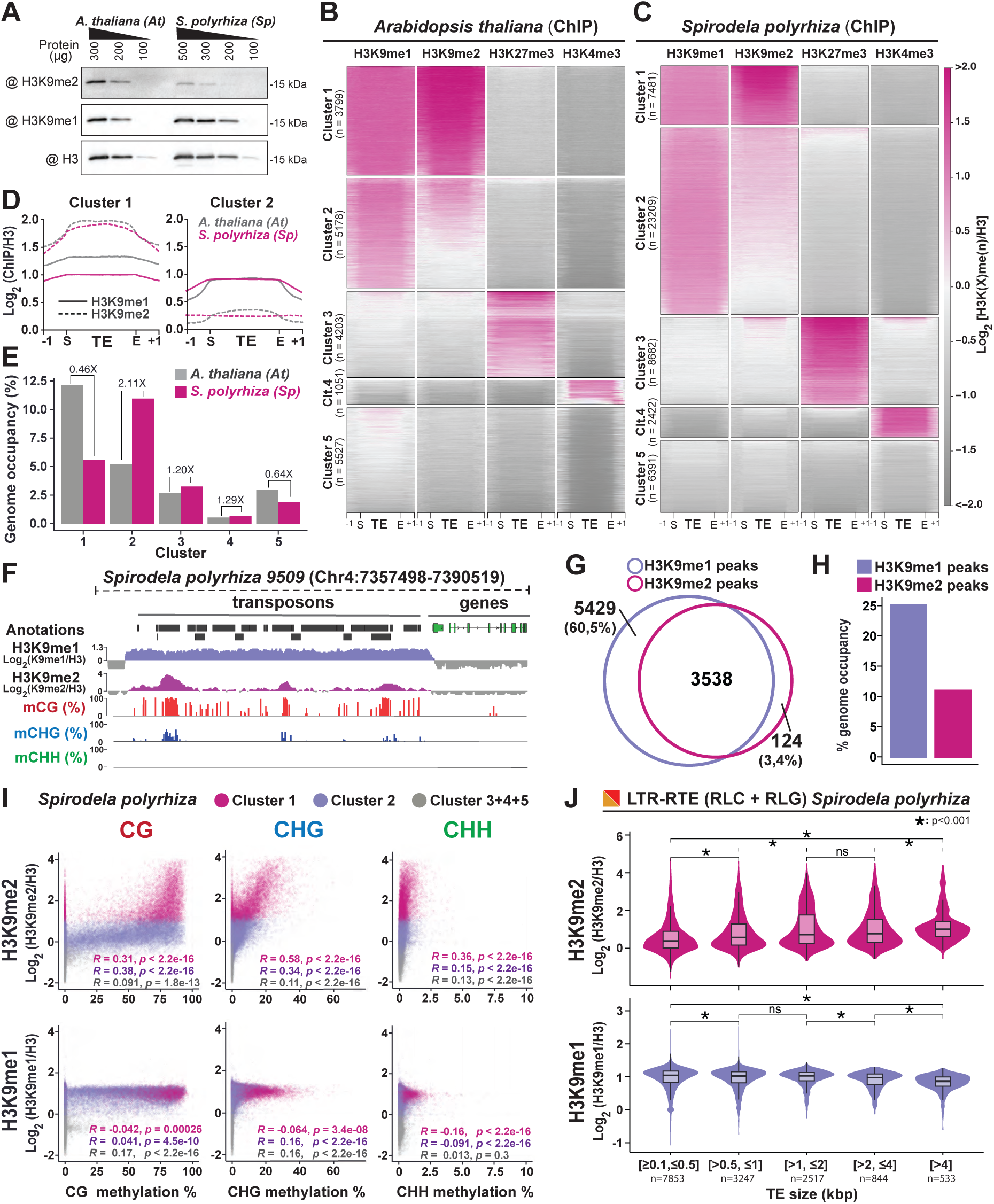
Patterns of H3K9me1/2 distribution in *Spirodela polyrhiza* transposons. (A) Protein blot analysis of H3K9me1 and H3K9me2 abundance in *Spirodela* compared to *Arabidopsis* at different protein amounts (μg). Total H3 protein levels are shown as loading control. (BC) Heatmap of *k*-means clustering of transposons in *Arabidopsis* (B) and *Spirodela* (C) using H3K9me1, H3K9me2, H3K27me3 and H3K4me3 enrichment over H3, presented as log2[ChIP(X)/ChIP(H3)]. Enrichment is represented by color, while depletion is shown in gray. Each row corresponds to a length-normalized TE annotation ± 1 kb. Transposons are grouped into clusters 1 to 5 based on the abundance of examined marks. (D) Metaplots of average weighted H3K9me1 (solid line) and H3K9me2 (dashed line) enrichments over H3 at Cluster 1 and 2 transposons ± 1 kb in *Arabidopsis* (colored) and *Spirodela* (gray). (E) Relative genome (%) occupancy of each of the TE clusters defined in B-C in *Arabidopsis* (grey) and *Spirodela* (colored). (F) Genome browser capture of the distribution of H3K9me1 and H3K9me2 enrichments, together with DNA methylation, along genes and transposons in *Spirodela*. (G) Venn diagram of the overlap between H3K9me1 and H3K9me2 peaks along *Spirodela* genome. (H) Genome occupancy (%) of H3K9me1 and H3K9me2 in *Spirodela* genome. (I) Scatter plots of DNA methylation in all three contexts against H3K9me1 and H3K9me2 enrichments of TEs in Cluster 1, 2 and 3/4/5 combined in *Spirodela*. *R* and *p* indicate Pearson correlation coefficients and P-values. (J) Analysis (violin and box plots) of H3K9me1 and H3K9me2 enrichments in different TE length groups for RLC and RLG combined in *Spirodela*. In all boxplots: mean is indicated by solid bar; the boxes extend from the first to the third quartile and whiskers stretch to the furthest value within 1.5 times the interquartile range. (*) indicates p-value <0.001. n.s.: not significant.

### A low proportion of TEs show high H3K9me2 enrichment in *Spirodela*

To further delve into the distribution of H3K9me1/2 in *Spirodela* and how the general organization of heterochromatic marks is affected by its reduced DNA methylation levels, we performed chromatin immunoprecipitation (ChIP) in *Spirodela* fronds and *Arabidopsis* 10 days-old seedlings (**Supplemental Fig. S25**) and focused on the association of these modifications with the TEs. In addition to H3, H3K9me1 and H3K9me2, we generated genomic profiles through ChIP-seq for H3K27me3 and H3K4me3. Although Polycomb-mediated H3K27me3 generally mediates developmentally regulated gene silencing, its association with TEs was investigated given that it also regulates TEs in the absence or loss of DNA methylation (Mathieu et al. 2005; Walter et al. 2016; Montgomery et al. 2020; Rougée et al. 2021), and may contribute directly to TE silencing (Hisanaga et al. 2023). H3K4me3, was taken as an euchromatic mark control (Houben et al. 2003; Zhang et al. 2009; Roudier et al. 2011; Ding et al. 2012; Pratx et al. 2023).

As observed for DNA methylation (Fig. 3A), H3K9me1/2 is concentrated at TE-dense regions in *Spirodela* (**Supplemental Fig. S26**), likely centromeric and pericentromeric heterochromatic regions (Houben et al. 2003; Jackson et al. 2004; Harkess et al. 2024). The reduced size of such regions (spanning a few kbps per chromosome, **Supplemental Fig. S26**) in *Spirodela*, compared to *Arabidopsis* larger pericentromeric regions (Mbps long), and their increased number (20 chromosomes in *Spirodela* in contrast to 5 in *Arabidopsis*), fits the speckled pattern of small dense DNA foci in DAPI staining of interphase nuclei of *Spirodela* (**Supplemental Fig. S27**). This is also compatible with previous observations of dispersed H3K9me2 foci in *Spirodela* nuclei immunostainings (Cao et al. 2015).

A more detailed *k*-means clustering analysis of histone mark profiles relative to H3 over TEs, identified 5 clusters with similar features in both *Arabidopsis* and *Spirodela* (Fig. 5B**,C**). Clusters 1 and 2 contain TEs enriched for H3K9me1/2 while clusters 3 and 4 consist of those marked by H3K27me3 and H3K4me3 respectively. Lastly, cluster 5 encompasses TEs depleted of any of the profiled marks (Fig. 5B**,C**).

Although cluster 1 and 2 TEs are marked by H3K9me1/2, they display homogeneous H3K9me1 but contrasting H3K9me2 levels (Fig. 5D**, Supplemental Fig. S28**). TEs in cluster 1 are more enriched in H3K9me2 than in H3K9me1 while in cluster 2 TEs H3K9me2 is below H3K9me1 (Fig. 5D). The two clusters comprise very different fractions of the two genomes and TE space. In *Spirodela*, cluster 1 only comprises about 20% of the TE space, representing less than 6% of the genome, while in *Arabidopsis* is the most abundant one, encompassing ∼35% of the TE space and ∼12% of the genome. Cluster 2, on the contrary, occupies 45% of the *Spirodela* TE space (more than 10% of its genome), but only about 20% in *Arabidopsis* (5% of the genome) (Fig. 5E**, Supplemental Fig. S29A**).

Given the similar genome size and TE content in both species, the opposed contribution of cluster 1 and cluster 2 to their genomes is in line with the levels of H3K9me2 observed by western blot and mass-spectrometry. Furthermore, similar H3K9me1 levels in both clusters concur with its even abundance in *Spirodela* and *Arabidopsis*. Hence, reduced DNA methylation levels in *Spirodela* correlate with a smaller fraction of the genome displaying a high H3K9me2 enrichment over TEs but it does not impact H3K9me1 distribution, nor it leads to a widespread compensatory regulation of TEs by H3K27me3. We noticed that, despite cluster 3 occupies a comparable fraction of TE sequences in both species (Fig. 5E), *Spirodela*, in contrast to *Arabidopsis*, displays TE-rich regions devoid of DNA methylation and H3K9me1/2 but rich in H3K27me3 (e.g. Chromosomes 1, 6, 10, 14, 20) (**Supplemental Fig. S26**). No obvious differences in TE-superfamily composition were observed between cluster 3 and clusters 1 and 2 (**Supplemental Fig. S29B**), suggesting no particular TE superfamily is preferentially regulated by Polycomb.

### H3K9me1 and H3K27me1 mark TEs independently of their length and methylation status in *Spirodela*

A closer inspection of the distribution of H3K9me1/2 along *Spirodela* TEs revealed that H3K9me1 uniformly covers TEs, while H3K9me2 peaks are found only at individual TEs (Fig. 5F). Furthermore, while more than 95% of H3K9me2 peaks overlapped with H3K9me1 domains, a significant fraction of H3K9me1 peaks (60,5%) did not with H3K9me2-associated regions (Fig. 5G). Moreover, most H3K9me2 peaks were embedded within H3K9me1 (**Supplemental Fig. S29C**). Adding to that, H3K9me1-enriched regions occupy 25% of the *Spirodela* genome whereas H3K9me2 peaks represent less than 12% (Fig. 5H). This indicates that, similarly to DNA methylation, high levels of H3K9me2 are only found within few TEs. Such observations prompted us to investigate the association between H3K9me1/2 and DNA methylation in *Spirodela* and indeed, H3K9me2 levels in *Spirodela* positively correlate with DNA methylation, including mCHG, like in *Arabidopsis* (Fig. 5I**, Supplemental Fig. S30**). Parallel to DNA methylation (Fig. 3), H3K9me2 increased with TE size (Fig. 5J). This further supports that, as previously inferred, the positive feedback loop between mCHG and H3K9me2 operates in *Spirodela*. However, this did not apply to H3K9me1. While in *Arabidopsis* TEs enriched in H3K9me1 invariably displayed high levels of DNA methylation (**Supplemental Fig. S30**), they did not so in *Spirodela* with its broad range of DNA methylation levels at TEs (Fig. 5I). Accordingly, H3K9me1 levels vary little across TE sizes (Fig. 5J).

These results indicate that, in *Spirodela*, H3K9me1 serves as broad TE mark but does not rely on DNA methylation as H3K9me2, suggesting that they might be differentially regulated. Although *Arabidopsis* H3K9 histone methyl-transferases (HMTs) can catalyze both H3K9me1 and me2 in vitro (Jackson et al. 2004; Ebbs and Bender 2006), it is unclear how their mono- and di-methyl-transferase activity is regulated in vivo or if both marks are deposited by the same HMTs. Loss of SUVH4, the main HMT in *Arabidopsis*, results in a strong reduction of H3K9me2 and loss of epigenetic silencing, only marginally affecting H3K9me1 and global heterochromatin organization (Jackson et al. 2002; Jasencakova et al. 2003; Jackson et al. 2004). This originally led to the proposition that H3K9me2 and H3K9me1 play different roles in silencing and heterochromatin maintenance respectively (Jackson et al. 2004).

Given that heterochromatin maintenance is not only relevant for TE silencing by as well for genome stability, we sought to investigate the distribution of other epigenetic marks associated with heterochromatin and genome stability. Specifically, H3K27me1, which in plants is largely associated with maintenance of constitutive heterochromatin, especially during DNA replication, and its distribution is not affected by loss of mCG (Mathieu et al. 2005; Shi and Dawe 2006; Jacob et al. 2009; Sequeira-Mendes et al. 2014; Montgomery et al. 2020; Jamge et al. 2023). *Arabidopsis* mutants lacking H3K27me1 do not only show loss of TE silencing but as well defects in chromatin organization and excess of repetitive DNA due to heterochromatin amplification during DNA replication (Jacob et al. 2009; Stroud et al. 2012; Dong et al. 2021). To study its distribution in *Spirodela*, we performed H3K27me1, along with H3K9me1/2, ChIP. The resulting profiles and *k*-means clustering analysis showed that H3K27me1 displays an overlapping pattern with H3K9me1, present at TEs independently of DNA methylation and occupying as well a large fraction of the genome (**Supplemental Fig. S31)**. Hence, in *Spirodela*, despite the reduced levels of DNA methylation and H3K9me2, H3K9me1 and H3K27me1 remain associated with TEs, likely to maintain heterochromatin and genome integrity.

### *5S rDNA* also display low levels of DNA methylation and H3K9me1/2 in *Spirodela*

Besides TE silencing, DNA methylation, H3K9me2 and RdDM are also involved in the epigenetic regulation of other repeats in plants. To investigate if the unique patterns seen on TEs also affect other repeats in *Spirodela*, we looked into the epigenetic landscape of the *5S rDNA* repeats. Present in hundreds or thousands of copies clustered in several large tandem arrays, most *5S rDNA* repeats are generally silenced in *Arabidopsis* and are well known targets of RdDM (Llave et al. 2002; Xie et al. 2004; Blevins et al. 2009; Simon et al. 2018). Although *Spirodela* and other duckweeds have been shown to have abnormally low numbers of *5s rDNA* repeats (<80-100 copies), two *5S rDNA* clusters in *S. polyrhiza* have been located on Chromosomes 6 and 13 (Hoang et al. 2018; 2020; Chen et al. 2021; 2024). A close inspection to the two *5S rDNA* clusters in *Spirodela* revealed clear differences with the epigenetic patterns found in *Arabidopsis* (**Supplemental Fig. S32A**). In *Spirodela*, *5S rRNA* repeats did not display DNA methylation in any context, nor H3K9me1/2 and H3K27me1/3 enrichments (**Supplemental Fig. S32B,C**). Nonetheless, in *Spirodela*, 24-nt siRNAs also mapped to *5S rDNA* repeats (**Supplemental Fig. S32A**). Although, in contrast to *Arabidopsis*, they were exclusively in sense to the *5S rDNA* loci and showed a guanosine 5’-nucleotide (5’G) bias (>75%) (**Supplemental Fig. S32D**) (not known to be preferentially loaded by any of the *Arabidopsis* AGOs), instead of the RdDM effector AGO4 characteristic 5’A signature seen in *Arabidopsis* (Mi et al. 2008; Havecker et al. 2010; Jullien et al. 2020).

Hence, lack of DNA methylation and H3K9me2 in *Spirodela* does not only impact degenerated and short TEs, but also other RdDM targets such as *5S rDNA* repeats, and further strengthens the hypothesis that such silencing mechanisms only apply to long, complete, TEs. Indeed, increased H3K9me2 and DNA methylation at long TEs hints that their self-reinforcement is focused at potentially functional or recently active ones to ensure their silencing.

### Long, complete TEs show signatures of RdDM activity

To gain further insights into the epigenetic patterns on long TEs, we focused on the complete LTR-RTEs of the *Copia* (RLC) and *Gypsy* (RLG) superfamilies, in which H3K9me2 levels were higher than those of H3K9me1. However, the patterns differed depending on the TE superfamily. While in RLCs had more H3K9me2 towards the center of the TE, the same modification peaked on the edges of RLGs (Fig. 6A). Such patterns resembled those found in *Arabidopsis* intact LTR-RTEs, (**Supplemental Fig. S33A**). DNA methylation at CG and CHG followed a similar trend as H3K9me2 in both cases (Fig. 6B), although mCHG was comparatively lower (Fig. 6B**, Supplemental Fig. S33B**). Furthermore, albeit very low, peaks of mCHH were also observable at RLGs edges (Fig. 6B). Hence, the layout of epigenetic silencing marks along complete TEs in *Spirodela* are very similar to those in *Arabidopsis*.

**Figure 6:**
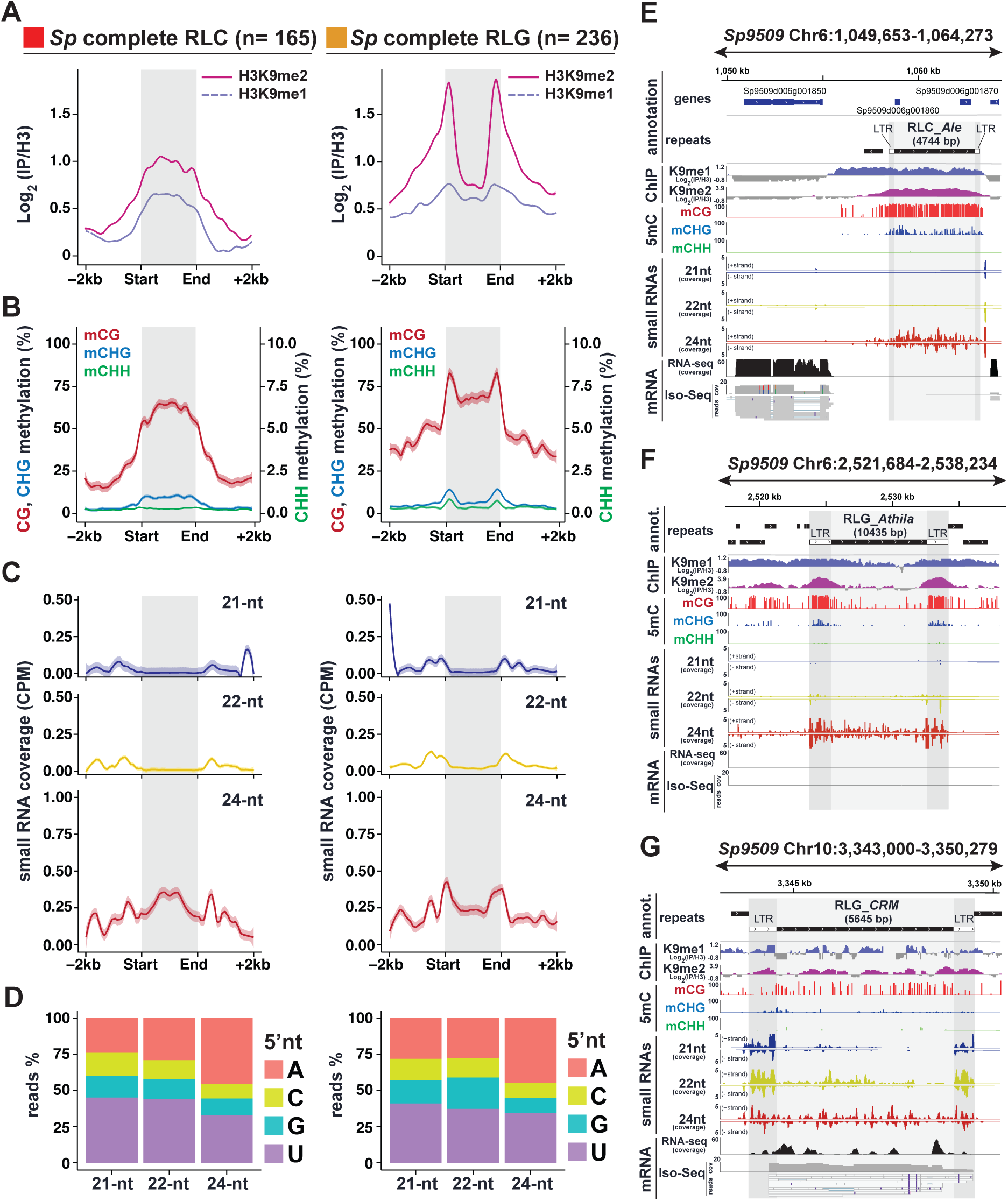
RdDM signatures in *Spirodela polyrhiza* complete TEs. (A) Metaplots of average weighted H3K9me1 (solid line) and H3K9me2 (dashed line) enrichments over H3 at complete RLC (left) and RLG (right) transposon annotations ± 2 kb in *Spirodela*. Complete TEs were defined as those for which all RLC and RLG protein domains where identified. n indicates number of complete elements in each superfamily. (B) Metaplots of averaged weighted DNA methylation in all three contexts over RLC and RLG transposons ± 2 kb flanking regions in *Spirodela*. mCHH values are scaled to 10% max values instead of 100%. (C) Metaplots of averaged weighted 21-, 22- and 24-nt small RNA abundance over intact RLC and RLG ± 2 kb flanking regions in *Spirodela*. In all metaplots (a-d), solid lines represent the mean and colored shadows the standard error. (D) 5’-nucleotide (5’nt) composition distribution (as % of reads) of 21-, 22- and 24-nt siRNAs mapping to intact RLC and RLG ± 2 kb flanking regions in *Spirodela*. (E-G) Genome browser captures of selected complete TEs in *Spirodela*. From top to bottom tracks display: gene and TE annotations, H3K9me1 and H3K9me2 enrichment profiles; mCG, mCHG and mCHH methylation; 21-, 22- and 24-nt siRNA mapping to Watson (+) and Crick (-) DNA strands; RNA expression by short (illumina) and long reads (Iso-Seq). Superfamily, clade and length (in bp) of TEs is indicated above their annotation. Light and dark gray shadow delimit TEs and LTRs respectively across all tracks.

Given that in *Arabidopsis* these patterns are partially determined by RdDM (Stroud et al. 2013; Zemach et al. 2013; Stroud et al. 2014), we inspected if small RNA mapped to complete TEs in *Spirodela*. Indeed, siRNAs derived from regions with DNA methylation and H3K9me2 were present in *Spirodela*, specially 24-nt siRNAs, despite at lower levels than in *Arabidopsis* (Fig. 6C**, Supplemental Fig. S33C**). Moreover, in contrast to *5S rDNA* siRNAs, 24-nt siRNAs mapping to complete TEs displayed an increased 5’A bias, characteristic of small RNAs loaded into AGO4 (Fig. 6D), although not as preferential as in *Arabidopsis* (**Supplemental Fig. S33D**). Complementarity of siRNAs with DNA methylation and H3K9me2 within the TE body and/or their LTRs were validated for individual complete TEs (Fig. 6E**-G****, Supplemental Fig. S34**).

To assess if TE-derived siRNAs might participate in RdDM in *Spirodela*, also given the unconventional 5’G bias of *5S rDNA* 24-nt siRNAs, we tested if its AGO4 (SpAGO4) has the same small RNA loading preference as *Arabidopsis* AGO4 (AtAGO4). To do so, we cloned genomic *AGO4* loci from both species with a Flag-HA tag for heterologous transient expression in *Nicotiana benthamiana* leaves. No Sp*AGO4* protein was detectable due to intron retention events when expressed in *N. benthamiana* (**Supplemental Fig. S35A-C**). This was likely due to the inability of dicots (*N. benthamiana*) to properly process introns of the monocots (*Spirodela*) (Keith and Chua 1986; Goodall and Filipowicz 1991; Lou et al. 1993). To circumvent this limitation, we amplified and cloned full length cDNA from one of the two *Spirodela* AGO4, *AGO4a*, and expressed it in *N. benthamiana* leaves (**Supplemental Fig. S35C-D**). To examine their small RNA loading preferences, AtAGO4 and SpAGO4a were immunoprecipitated (IP) from *N. benthamiana* leaf lysates (Jullien et al. 2020) (Fig. 7A) and small RNAs extracted and sequenced. Like AtAGO4, SpAGO4a displayed a strong preference for 5’A 24-nt long siRNA and reduced affinity for 21-nt siRNA, compared to the population of siRNAs present in *N. benthamiana* leaves (Fig. 7B**,C**).

**Figure 7:**
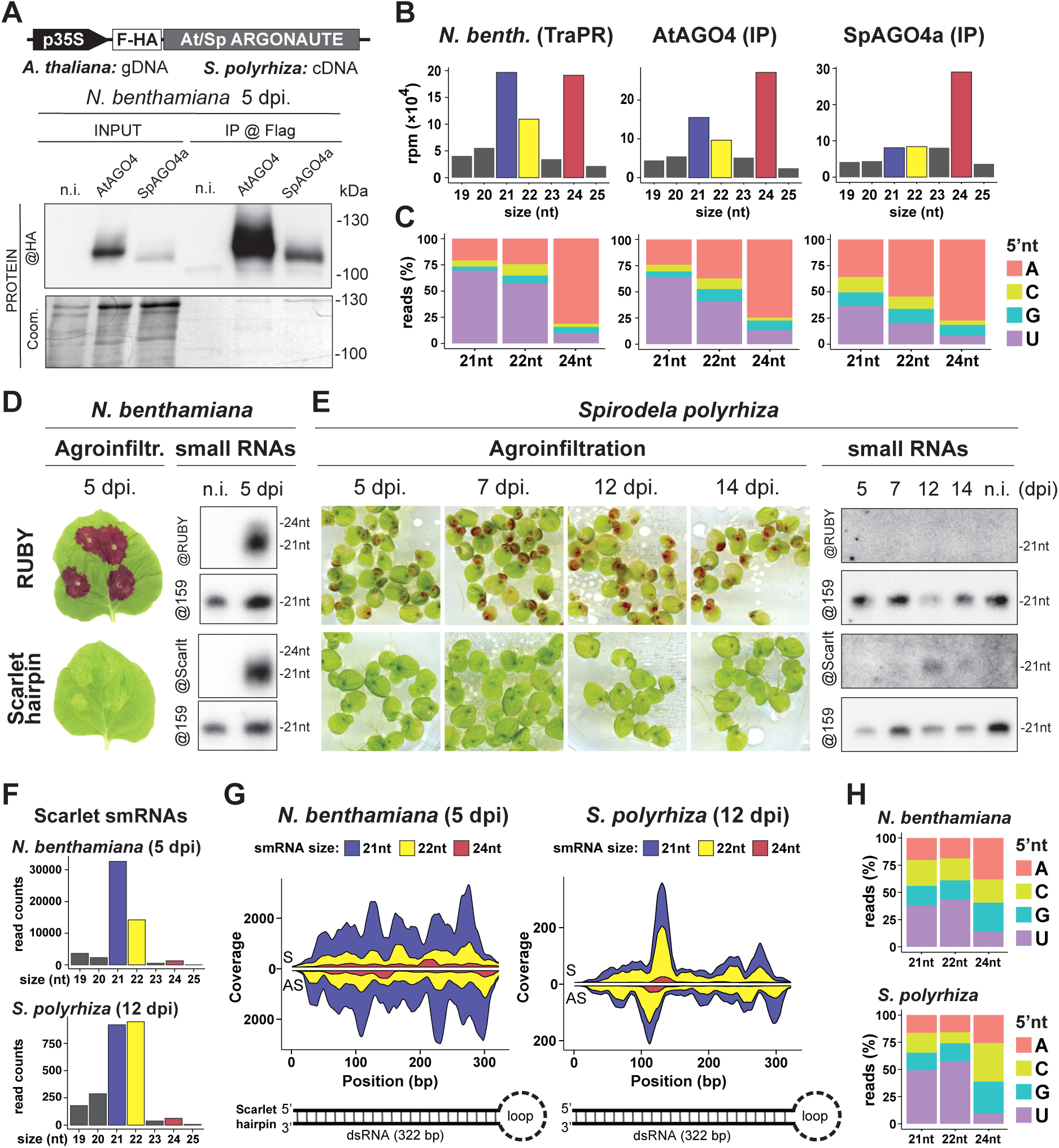
*Spirodela polyrhiza* AGO4a siRNA loading preferences and IR-derived siRNAs in transient expression. **(A)** Protein blot analysis of *Arabidopsis* and *Spirodela* AGO4 (AtAGO4 and SpAGO4a) input and IP fractions from transient expression in *N. benthamiana* 5 days post infiltration (dpi). n.i.: non-infiltrated. Coomassie (Coom.) staining of blot is shown as control for input and IP fractions. Scheme of constructs infiltrated in *N. benthamiana* leaves for transient expression is shown above. p35S: CaMV 35S promoter; F-HA: in-frame N-term Flag and HA peptide tags. Genomic (gDNA) or cDNA origin of the AGO4 sequences is indicated. **(B)** Size distribution and abundance in reads per million (rpm) of TraPR purified *N. bethamiana* siRNAs (as control) and siRNAs extracted from AtAGO4 and spAGO4a IPs. **(C)** 5’-nucleotide (5’nt) composition distribution (as % of reads) of 21-, 22- and 24-nt siRNAs from b. **(D)** Representative pictures of *N. benthamiana* leaves agroinfitrated for transient expression of RUBY or the scarlet hairpin (*hpScarlet*) IR next to small RNA blot analysis in non-infiltrated and 5 dpi leaves. miRNA159 is shown as loading control. **(E)** Representative pictures of *Spirodela* fronds agroinfitrated for transient expression of *RUBY* or *hpScarlet* and small RNA blot analysis in non-infiltrated and infiltrated samples at different timepoints. miRNA159 as loading control. **(F)** Size distribution and abundance of small RNAs mapping to *hpScarlet* from 5 dpi *N. bethamiana* and 12 dpi *Spirodela*. **(G)** Stacked distribution of sense (S) and antisense (AS) small RNA coverage by size along *hpScarlet* hairpin in *N. bethamiana* and *Spirodela*. Schematic representation of the IR is depicted below. **(H)** 5’-nucleotide (5’nt) composition distribution of 21-, 22- and 24-nt siRNAs mapping to *hpScarlet* in *N. bethamiana* and *Spirodela*.

Therefore, small RNAs might be indeed involved in maintaining DNA methylation and H3K9me2 in *Spirodela* at long complete TEs through canonical RdDM. Our analysis of silencing pathways and epigenetic patterns on TEs shows that core components required for RdDM are present in the *Spirodela* genome and, although poorly expressed, probably active during vegetative clonal propagation to silence long TEs.

### Transcriptionally active TEs are source of 21/22-nt-siRNAs in *Spirodela*

We also detected the presence of very low amounts of 21-22-nt siRNAs at the edges of complete TEs (Fig. 6C). In some instances, 24-nt siRNAs overlapped with equally abundant 21-22-nt siRNAs, especially at the flanking LTR sequences (Fig. 6G**, Supplemental Fig. S36**). 21-22-nt siRNAs can arise from TE edges in *Arabidopsis* (**Supplemental Fig. S33C**), as result of DCL2/4 processing of Pol IV transcripts, and direct DNA methylation (Nuthikattu et al. 2013; McCue et al. 2015; Panda et al. 2020; Sigman et al. 2021). It is unlikely that the observed TE-derived 21-22-nt siRNAs participate in RdDM, due to the low affinity of SpAGO4 to bind such siRNA sizes (Fig. 7B), and those mapping to TEs have less 5’A bias but more 5’U (Fig. 6D), a signature of AGO1-loaded small RNAs (Mi et al. 2008; Havecker et al. 2010). This hints to a potential role rather in post-transcriptional silencing (PTGS). During PTGS, 21-22-nt siRNAs loaded into ARGONAUTE proteins of the AGO1/5/10/18 and AGO2/7 clades target Pol II transcripts to guide mRNA cleavage or translational repression (reviewed in (Fang and Qi 2016)). Coincidentally, we observed expression of TEs producing 21-22-nt siRNAs in our transcriptome datasets, including, in some intances, full-length transcripts in the long-read cDNA sequencing Iso-Seq data (Fig. 6G**, Supplemental Fig. S35**).

Although PTGS is a well-documented control mechanism against transcriptionally active TEs in *Arabidopsis* (Teixeira et al. 2009; Slotkin et al. 2009; Marí-Ordóñez et al. 2013; Creasey et al. 2014; Lee et al. 2020; Oberlin et al. 2022), the potential role of 21-22-nt siRNAs in TE silencing in *Spirodela* remains to be experimentally tested.

### 22-nt siRNAs are produced in *Spirodela* despite the lack of a DCL2 orthologue

We were intrigued by the presence of TE-derived 22-nt siRNAs, given that any identifiable ortholog of DCL2, their cognate Dicer in plants (Xie et al. 2005; Henderson et al. 2006; Wang et al. 2018; Jia et al. 2020), is missing in *Spirodela* and duckweed genomes in general (Fig. 2B**, Supplemental Fig. S8**) (Ernst et al. 2023; Harkess et al. 2024).

Their co-occurrence with 21-nt siRNAs, (Fig. 6G**, Supplemental Fig. S35**) suggested a common RNA substrate. Given the low amounts of 21-22-siRNAs mapping to individual TEs, to rule out artifacts and inspect if other endogenous siRNA sources are as well processed into 22-nt siRNAs, we investigated *Spirodela* trans-acting siRNAs (tasiRNAs). In *Arabidopsis*, trans-tasiRNAs are processed from transcripts of specific loci (TAS), mainly into 21-nt-siRNA by DCL4 to, through their association with AGO1, regulate several target mRNAs by PTGS (Yoshikawa et al. 2005). As the TAS3 locus had been previously identified in *Spirodela* (Michael et al. 2017), we scored for TAS3 matching siRNAs in our TraPR datasets. In contrast to *Arabidopsis*, where TAS3 tasiRNAs are almost exclusively 21-nt long, in *Spirodela*, TAS3 tasiRNAs were found as 21- and 22-nt, the later accounting for about one third of TAS3 tasiRNAs (**Supplemental Fig. S37A,B**). Their 5’U bias also suggest their preferential loading into AGO1 (**Supplemental Fig. S37C**).

Next, to independently test if *Spirodela* can produce 22-nt siRNAs in the absence of DCL2, we attempted to stimulate small RNA biogenesis through expression of suitable exogenous substrate RNAs. As stable transformation in *Spirodela* is difficult, we chose *Agrobacterium*-mediated transient expression. This technique has been widely used to trigger and investigate silencing mechanisms, owing to the quick onset of siRNA production in response to expression of double-stranded RNA (dsRNA) following infiltration of plant tissue with *Agrobacterium* carrying the corresponding transgenes (Hamilton et al. 2002; Bertazzon et al. 2012; Andrieu et al. 2012; Sun et al. 2014; Jay et al. 2023). We developed a modified protocol for frond transformation in *Lemna minor* (Yang et al. 2018) and applied it successfully to induce transient transgene expression in *Spirodela* fronds (**Supplemental Fig. S38**). We used the visual reporter *RUBY* (He et al. 2020) and an inverted repeat (IR) derived from the coding sequence of the fluorescent reporter *Scarlet* (*Scarlet* hairpin, *hpScarlet*) (Incarbone et al. 2023). Both constructs triggered siRNAs when infiltrated into *N. benthamiana* as expected (Fig. 7D). In *Spirodela*, however, we could not detect *RUBY*-derived siRNAs by RNA blotting even after two weeks post-infiltration. Given transgene sense PTGS (S-PTGS) is generally associated with high levels of transgene expression, the apparent lack of S-PTGS in *Spirodela* might be due to reduced expression of the reporter compared to *N. benthamiana*. On the other hand, *hpScarlet*-derived siRNAs were observed 12- and 14-days post-infiltration and were lost shortly after (Fig. 7E), likely due to loss of transgene expression inherent to the *Agrobacterium*-mediated transient expression method. Sequencing sRNAs from both *N. benthamiana* and *Spirodela* infiltrated with *hpScarlet* showed that the IR is processed into 21-22-nt siRNAs in both plants. However, *Spirodela* produced relatively equal amounts of 21- and 22-nt siRNAs, as observed for TEs, whereas the profile in *N. benthamiana* is dominated by 21-nt siRNAs (Fig. 7F). Both siRNA sizes were distributed along the length of the IR dsRNA (Fig. 7G) and displayed the 5’U bias signature of AGO1 loading (Fig. 7H).

Hence, *Spirodela* is capable of producing 22-nt siRNAs in the absence of DCL2, supporting that 21-22-nt siRNAs mapped to TEs are *bona-fide* siRNAs. Given the similar 21-22-nt siRNA patterns observed between TEs, miRNA-triggered secondary tasiRNA and IR-derived siRNAs, 21-22-nt siRNAs mapped to transcriptionally active TEs are likely the result of: (i) a shared miRNA targeting event to initiate secondary siRNAs, shown to regulate some TEs in other plants (Creasey et al. 2014; Šurbanovski et al. 2016), or; (ii) direct processing of RNA hairpins commonly present in LTRs (Benachenhou et al. 2013).

## Discussion

Based on high resolution genomic information of a duckweed species during its clonal reproduction, we have provided a detailed analysis of TE content, of the epigenetic configuration at TEs, and several components expected to contribute to the silencing pathways that control TE activity in other angiosperms. Our results reinforce the idea that, with few exceptions, the *Spirodela* genome encodes only a reduced set of both PTGS and TGS components compared to *Arabidopsis*, some of which are absent or display a reduced set of paralogues, partially accounting for its reduced levels of DNA methylation and other epigenetic marks generally associated with TEs. The unique epigenome of *Spirodela* is the result of the loss of DNA methylation, and reduced levels of H3K9me2 associated to it, in highly abundant degenerated TE remnants scattered across the genome which remain nonetheless associated to the heterochromatic marks H3K9me1 and. H3K27me1. In contrast, potentially intact or recently active TEs display all the epigenetic signatures associated with silencing in angiosperms. Another interesting observation is the loss of any of the silencing marks inspected here in *5S rDNA* repeats. Although this might reflect the preferential selectivity of RdDM for long TEs, lack of silencing might as well be due to the reduced number of *5S rDNA* repeats in *Spirodela*, compared to other angiosperms, to ensure enough 5S rRNA expression for ribosome biogenesis. These changes in epigenetic regulation and silencing might be a consequence of the simplified morphology or reduced tissue and cell types in duckweeds compared to other angiosperms (Wang et al. 2014; Michael et al. 2020; Abramson et al. 2021; Ware et al. 2023; Ernst et al. 2023). However, it could also be a consequence of their different lifestyle or propagation mode.

*Spirodela* displays a broad range of adaptations to their free-floating aquatic lifestyle, such as the loss of complete stomatal closure response to prevent water loss (Fang et al. 2023). Besides TE control, RdDM plays a role in the regulation of gene expression and physiological adaptation to several abiotic stresses, such as heat, drought, salinity, or low environmental humidity (Erdmann and Picard 2020; Liu and He 2020; Urquiaga et al. 2021). Given that water supply is not limiting in aquatic environments, and their higher stability to fluctuations in these parameters compared to terrestrial ones, RdDM activity in adult somatic tissues to respond to stress might have become less relevant in *Spirodela*.

Alternatively, but not mutually exclusive, the epigenetic landscape in the *Spirodela* genome might be a consequence of their reproduction strategy. Asexual reproduction has been linked to lower TE activity and content as TEs require sexual reproduction to spread horizontally within populations. In absence of sexual reproduction, TEs may instead become locked in vertical clonal lineages. Additionally, they might go extinct if their proliferation becomes detrimental for the host without increased purifying selection of new TE insertions facilitated by sex (Arkhipova and Meselson 2000; Wright and Finnegan 2001; Bast et al. 2019; Dechaud et al. 2019). Furthermore, RdDM activity over short TE remnants, leading to the production of 24-nt siRNAs, is seen as an immunity memory against mobilization or invasion by related intact TEs (Nguyen and Gutzat 2022), akin to the piRNA pathway in animals (Czech et al. 2018). Thus, TEs potentially being less active during *Spirodela* clonal propagation might have led to the loss of DNA methylation, RdDM expression, and lack of components required for its activity during vegetative development such as SHH1 and CLSY1/2 (Zhou et al. 2022) over TE fragments. Maintenance of heterochromatin at such repetitive TEs remnants through H3K9me1 and H3K27me1 independently of DNA methylation might ensure proper chromatin organization and stability, preventing TE interference with transcriptional and RNA processes (Kim and Zilberman 2014; Huff et al. 2016; Sammarco et al. 2022; Ilık et al. 2024), illegitimate recombination or re-replication of repeats during cell division (Jacob et al. 2009; Stroud et al. 2012; Chénais et al. 2012; Bourque et al. 2018; Dong et al. 2021). The presence of dense TE clusters marked by H3K27me3 might also prevent genome instability as their alternative regulation by DNA and H3K9 methylation could mimic centromeric and pericentromeric regions, leading to the formation of polycentromeric chromosomes highly unstable during mitosis (Fu et al. 2012). As inactivation of secondary centromeres has been linked to increased H3K27me2/3 in wheat (Zhang et al. 2010), such TE-rich clusters might correspond to inactive centromeres in *Spirodela*. Deeper characterization of the chromatin structure (e.g., chromatin accessibility) and landscape (additional histone modifications, histone variants …), together with functional studies, might shed light onto the regulation of H3K9me1, H3K27me1, H3K27me3 and their role in shaping heterochromatin, TE silencing and genome stability regulation in *Spirodela* and other plants.

Furthermore, unlike vegetative propagation in other plants where clones originate from differentiated organs such as stems, roots or leaves, with tissue-specific and developmentally regulated TEs, new individuals in duckweeds originate from a dedicated group of stem cells (Landolt 1986). This might have further favored a relaxation of epigenetic and RdDM-mediated TE control in somatic tissues which do not participate in vegetative reproduction in *Spirodela*. Nevertheless, potentially intact TEs are under several layers of epigenetic control, including RdDM, suggesting the pathway might be tissue-specific or developmentally regulated.

In contrast to stem cells in other angiosperms, contained in a well-organized shoot apical meristem (SAM) that generates new organs (Gaillochet and Lohmann 2015), meristems in *Spirodela* and all duckweeds are formed by unstructured groups of few undifferentiated cells, which generate new meristems that develop into daughter fronds. (Landolt 1986; Lemon and Posluszny 2000; Sree et al. 2015; Yang et al. 2021; Li et al. 2023). In order to maintain genome integrity along the clonal lineage, RdDM might be expressed in the few stem cells within the budding pockets or during the developmental window between the formation of a new meristem and its development into a new frond with its own stem cells.

Furthermore, lack of, or weakened, RdDM activity due to losses such as DRM3, CLSY1, SHH1 and absence of CMT2, in combination with weak CMT3/ZMET activity in somatic tissues during development, will suffice to explain the global loss of mCHH and low levels of mCHG. Additionally, the higher variation in mCG levels in *Spirodela* could be a consequence of mCG maintenance being less efficient, not active in all cells, or not all CG sites being as faithfully maintained. Altogether, this might, in some instances, permit transcription of some TEs that then trigger 21-22-nt siRNAs. Alternatively, transient relaxation of TE control and TE expression has been shown to take place in *Arabidopsis* meristematic stem cells during developmental transitions (Gutzat et al. 2020). A similar phenomenon taking place during the development of new meristems and fronds might as well explain the paradox of observed expression from TEs covered in silencing marks and the apparent simultaneous presence of small RNAs associated with TGS and PTGS. Further tissue- and cell-specific investigation of epigenetic parameters will be able to address these hypotheses.

Nonetheless, although infrequent, *Spirodela* does flower occasionally (Fourounjian et al. 2021) and its population genetic structure shows signs of sexual reproduction (Ho et al. 2019; Wang et al. 2024). Hence, RdDM might still be relevant and active during sexual reproduction as suggested by the presence of CLSY3, which recruits RdDM to TEs during flowering in *Arabidopsis* (Long et al. 2021; Zhou et al. 2022). Nonetheless, we note that the function of silencing components present and absent in *Spirodela* might play different roles in duckweeds (and other monocots) compared to those described in *Arabidopsis* given their evolutionary distance. Hence, a full understanding of TE silencing in duckweeds will require further molecular genetics and biochemical characterization of their silencing pathways.

Adding to the unique epigenetic regulation of degenerated TEs and their relics in *Spirodela* here described, one of the intriguing results of our study is the constatation that, despite lack of DCL2, *Spirodela* can produce 22-nt siRNAs, nearly as abundantly loaded in AGOs as 21-nt siRNAs, not only from TEs but also from endogenous and exogenous sources of dsRNA. As these sources of siRNAs are generally products of DCL4 in plants (Xie et al. 2005; Fusaro et al. 2006; Parent et al. 2015), given the overlap between 22- and 21-nt but not 24-nt (DCL3), we consider processing dsRNA into 22-nt siRNAs by *Spirodela* DCL4 as a plausible explanation. Alternatively, DCL3 in *Arabidopsis* has been shown to be able to process certain dsRNA precursors into 22-nt siRNAs *in vitro* (Loffer et al. 2022) and is responsible for 22-to-24-nt siRNA biogenesis in the moss *Physcomitrium patens* (Cho et al. 2008). Therefore, *Spirodela* DCL3 might as well be responsible for the biogenesis of 22-nt siRNAs in *Spirodela*. Future genetic and biochemical characterization of *Spirodela* DCLs will be required to understand the factors responsible for the biogenesis of 22-nt siRNAs and their role in *Spirodela* and other duckweeds. In *Arabidopsis*, DCL2 and its capacity to process dsRNA into 22-nt siRNAs have been proposed to play a dual role in antiviral immunity. On one hand its processing of viral dsRNA acts as an innate immunity sensor (Nielsen et al. 2023; 2024). On the other hand, 22-nt siRNAs in plants guide the recruitment of RDRs to boost siRNAs production from their targets and amplify silencing (Chen et al. 2010; Zhang et al. 2012; Taochy et al. 2017; Chen et al. 2018). Their presence in *Spirodela* independently of DCL2 and the lack of several other RNAi factors associated with antiviral defense in angiosperms, as well as the apparent lack of sense transgene silencing (S-PTGS) triggered by transient expression of the *RUBY* reporter, raises several intriguing questions about RNA-based defense mechanisms in duckweeds that deserve further investigation in the future.

Hence, *Spirodela* might represent a unique model to investigate not only transposon dynamics and the epigenetic and genetic consequences of long periods of asexual reproduction but also noncanonical PTGS pathways.

## Methods

### *Spirodela* culture and growth conditions

#### Spirodela polyrhiza

*Spirodela* was grown on half-strength Schenk and Hildebrandt (**½** SH, Duchefa Biochemie, #S0225) liquid medium in Magenta^TM^ GA-7 containers (Sigma Aldrich, #V8505), culture dishes (Greiner Bio-One #664160), 6-wells culture plates (Greiner Bio-One #657185), or in 1L glass beakers (covered with a sterile plastic lid), all sealed with Leucopore tape (Duchefa Biochemie #L3301), under long-day conditions (16h light /8h dark) at 21°C with a light intensity of 85 µM m^-2^ s^-1^. The *Spirodela polyrhiza* #9509 clone was obtained from the Landolt collection (Zürich, Switzerland), recently transferred to the CNR-IBBA_MIDW collection (Milano, Italy: https://biomemory.cnr.it/collections/CNR-IBBA-MIDW) (Morello et al. 2024).

Further information regarding *Spirodela* sterilization for axenic cultures and long-term storage, *Arabidopsis* and *N. benthamiana* material information and culture and plant imaging can be found within Supplemental Methods in Supplemental Material.

#### Genome sequencing and gene annotation

Genomic DNA from *Spirodela* was purified using Dneasy® Plant mini kit (Qiagen #69104) following the manufacturer’s instructions. Library preparation was carried using NEBNext® Ultra™ II DNA Library Prep Kit for Illumina® (NEB #E7103). Sequencing was performed on an Illumina HiSeq4A to produce paired-end reads of 125 bp. The Landolt Sp9505 genome was generated by pseudo-assembly with MaSuRCA (Zimin et al. 2013) (version 3.4.1, default parameters), using as scaffold the Sp9509_Oxforrd_v3 reference assembly(Hoang et al. 2018). The pseudo-assembly was polished using POLCA (Zimin and Salzberg 2020). Variation, between the two assemblies, was further evaluated using GAKT(McKenna et al. 2010) (version 4.0.1.2), in particular with HaplotypeCaller (Poplin et al. 2018) (--standard-min-confidence-threshold-for-calling 30). Variants identified were summarized into a table using GAKT VariantsToTable (-F CHROM -F POS -F TYPE -GF AD). Gene annotations were transferred from the latest annotated version of Sp9509 (Ernst et al. 2023) obtained from (https://www.lemna.org). Annotation transfer was performed using Liftoff (v1.6.3) (Shumate and Salzberg 2021) using default parameters. Information regarding the phylogenetic analysis and protein domain annotations of *Spirodela* silencing components can be found within Supplemental Methods in Supplemental Material.

#### Transposon annotation

The annotation of transposable elements (TEs) in *Spirodela* polyrhiza was performed by combining EDTA (Ou et al. 2019) and RepeatModeler2 (Flynn et al. 2020) tools. To ensure accuracy, both tools were run in triplicate and the resulting annotation libraries were aligned and merged. The classification of TEs was further refined using DeepTE (Yan et al. 2020) (-sp P). The presence of the protein domain for LTR-RT was evaluated using TEsorter (Zhang et al. 2022) (version 1.3, -db rexdb-plant -st nucl).

#### Transcriptome analysis

To extract RNA from *Spirodela*, 100 mg of plant material was flash-frozen upon harvesting and ground. The samples were homogenized in 1 mL of TRIzol^TM^ (ThermoFisher, #15596018) using Silamat S6 (Ivoclar Vivadent). RNA was extracted according to manufacturer’s instructions. RNA library preparation was performed using the NEBNext® Ultra™ II RNA Library Prep Kit for Illumina® (NEB #E7775) or the SMRTbell Prep Kit 3.0 for Illumina and Iso-Seq respectively. Illumina sequencing was performed on HiSeq5A to generate 125 bp pairedend reads. Long read sequencing was performed on PacBio® Sequel® II. For Illumina data, base calls were performed using bcl2FASTQ (v2.17). RNA-seq reads were trimmed using Trim Galore! (v0.6, --illumina -phred33 -stringency 1 --FASTQc --length 20). Trimmed reads were mapped using HISAT2 (v2.1.0). Gene and transposable element expression levels were quantified separately using kallisto (Bray et al. 2016) (v0.46) with default parameters. The resulting expression levels (measured in TPM) were then averaged across the three replicates. For *Arabidopsis* seedlings, data was obtained from (GSM6892967, GSM6892968, GSM6892969) (Li et al. 2023). Iso-Seq long reads were processed using nf-core/isoseq (https://github.com/nf-core/isoseq/tree/1.0.0). High quality transcripts were then mapped to *Spirodela* genome using minimap2 (Li 2021) (v2.17, default parameters).

#### DNA methylation analysis

For EMseq, DNA extraction was performed as above. Libraries were prepared using the NEBNext® Enzymatic Methyl-seq Kit (NEB, #M7634) following the manufacturer’s instructions. Sequencing was performed on an Illumina NovaSeq SP to produce paired-end reads of 150 bp. Briefly, basecalls were performed using bcl2FASTQ (v2.17) and sequenced reads were quality filtered and adaptor trimmed using Trim Galore! version 0.6.2 (https://github.com/FelixKrueger/TrimGalore). Enzymatic-converted reads were aligned either to the TAIR10 genome or *Spirodela* polyrhiza genome using Bismark version 0.22.2 with *Bismark* (--non_directional *-q* --score-min *L,0,–0.4*) (Krueger and Andrews 2011) to generate coverage files per cytosine. Conversion rates were assessed by mapping (SAMtools v1.9) the reads to the unmethylated lambda phage DNA, with the same parameters used for the genomes BAM files containing clonal deduplicated and uniquely mapped reads were then used to extract weighted methylation rates at each cytosine as described in (Schultz et al. 2015). Replicates were merged into a single bedFile using the BEDTools (Quinlan and Hall 2010) merge function and only cytosines with coverage with ≥4 were considered for further analyses. 5mC was calculated using ViewBS (Huang et al. 2018) (v0.1.11) (https://github.com/xie186/ViewBS) using default parameters.

#### Small RNA library preparation and analysis

The libraries were prepared according to (Hayashi et al. 2016). In summary, small RNAs purified from plant tissues or immunoprecipitated AGO (Supplemental Methods in Supplemental Material), using TraPR (Grentzinger et al. 2020) or from total small RNA as described in Supplemental Methods (Supplemental Material), were ligated to 3′ barcoded DNA adapters using truncated T4 RNA ligase 2 (New England Biolabs, #M0373). These fragments were separated in an 12% denaturing polyacrylamide-urea gel, excised, and purified using ZR small-RNA PAGE Recovery Kit (Zymo Research, # R1070). The elution was then ligated to 5′ barcoded RNA adapters using T4 RNA ligase 1 (New Englland Biolabs, #M0204). To reduce ligation biases, the sequence of both adaptors included 4 random nucleotides at the ends. The resulting fragments were reverse transcribed and amplified by PCR using primers compatible with the Illumina Platform. The size distribution of the libraries was assessed using the 5200 Fragment Analyser System (Agilent, #M5310AA). Sequencing was performed on an Illumina NextSeq550 to produce single-end reads of 75 bp. Basecalls were performed using bcl2FASTQ (v2.17). Reads were trimmed and quality checked using Trimmomatic (Bolger et al. 2014) (v0.39). Trimmed reads were mapped to *Arabidopsis* TAIR10 and *Spirodela* genomes, and *RUBY* or *hpScarlet* sequences using Bowtie 2 (Langmead and Salzberg 2012) (v1.2.2; -e 50 -a -v 0 --best --strata --nomaqround -y --phred33-quals --no-unal --sam). Mapped reads were then normalized using bamCoverage (Ramírez et al. 2016) (DeepTools v,--normalizeUsing CPM --binSize 1 --smoothLength 10 --maxFragmentLength 30). To visualise strand, mapped reads were split using SAMtools view (Li et al. 2009) (SAMtools v1.9, using -h -b -F 16 or -h -b -f 16 options).

#### Chromatin immunoprecipitation (ChIP)

Chromatin immunoprecipitation (ChIP) was conducted on 2 g of *Arabidopsis* 10-day-old seedlings (WT Col-0) and *Spirodela polyrhiza* #9509 fronds. Following standard procedures (see detailed method in Supplemental Methods in Supplemental Material) using the following antibodies: anti-H3 (rabbit polyclonal, Abcam, #ab1791), anti-H3K27me3 (rabbit polyclonal, Merck-Millipore, #07–449), anti-H3K4me3 (rabbit polyclonal, Abcam, #ab8580), anti-H3K9me1 (rabbit polyclonal, Abcam, 3ab8896), anti-H3K9me2 (mouse monoclonal, Abcam, #ab1220) or anti-H3K27me1 (rabbit polyclonal, Merck-Millipore, #17–643). Further detailed information on ChIP sample preparation and histone PTM procedures and analysis by mass spectrometry, can be found within Supplemental Methods in Supplemental Material.

#### ChIP-seq library prep and data processing

For ChIP-seq, libraries were prepared using the NEBNext® Ultra™ II DNA Library Prep Kit for Illumina® (NEB, #E7103) according to the manufacturer’s instructions. Sequencing was performed on an Illumina NovaSeq S4 to produce paired-end reads of 150 bp. The raw BAM files were converted to FASTQ using BEDTools (Quinlan and Hall 2010) (v2.27.1, default parameters) for quality assessment of the sequenced samples. FASTQc (https://www.bioinformatics.babraham.ac.uk/projects/FASTQc/) (v0.11.8, default parameters) was used to generate quality reports for all sequencing data. Basecalls were performed using bcl2FASTQ (v2.17). Trimming of reads was performed using trim_galore (https://zenodo.org/doi/10.5281/zenodo.5127898) (v0.6.5, --dont_gzip --stringency 1 --fastqc--length 5’). The aligned reads were then mapped to either the TAIR10 *Arabidopsis* genome or *Spirodela* polyrhiza genome using Bowtie 2 (Langmead and Salzberg 2012) (v2.4.1, default parameters). Duplicate reads were removed using Picard (http://broadinstitute.github.io/picard/) (v2.22.8, default parameters). Mapped reads were then normalized using bamCoverage (deepTools v3.3.1). Correlations between ChIP samples were evaluated using deepTools (Ramírez et al. 2016) (v3.1.2, default parameters). The bamCompare function from deepTools was used to normalize ChIP samples to Input or H3 and produce log2 ratio (ChIP/H3) bigWig files. Peak calling was performed using MACS2 (v2.2.5 –nomodel –nolambda –broad -q 0.01 -f BAMPE -g 138550563) (Zhang et al. 2008).

#### Plasmids used in this study

All plasmids generated have been submitted to Addgene (www.addgene.org) and can be retrieved under the following IDs: p*35S*:*FHA*-At*AGO4*_gDNA (#216838); p*35S*:*FHA*-Sp*AGO4a*_gDNA (#216841); p*35S*::*FHA*-Sp*AGO4a*_cDNA (#216842). The following plasmids previously originated in (He et al. 2020) and (Oberlin et al. 2017) were obtained from Addgene: p*35S*:*RUBY* (#160908); p*ZmUbq*:*RUBY* (#160909); p*35S*:*GFP-GUS* (#167122). p*3*At*UBQ*:*hpScarlet* was a gift from Marco Incarbone (Max-Planck-Institute of Plant Physiology, Golm, Germany). Cloning and transient expression procedures in both *Spirodela* and Tobacco can be found in Supplemental Methods (Supplemental Material). Primers used for cloning can be found in **Supplemental Fig. S39**.

#### Small RNA and protein blotting

RNA and protein gels, blotting and detection were performed using standard procedures detailed in Supplemental Methods (Supplemental Material) together with probes (**Supplemental Fig. S39**) and antibodies used in this study.

#### Staining of nuclei

DAPI-staining of interphase nuclei was performed as described in Supplemental Methods (Supplemental Material).

#### Visualization and Statistical analysis

Data visualization files were generated according to each specific sections. Statistical tests were performed using R-dependent ggpubr (v0.6.0) (https://rpkgs.datanovia.com/ggpubr/), statistical test performed are indicated accordingly on each figure legend figure. Plots under R (v4.2.2) (https://www.r-project.org) were generated using Circlize (Gu et al. 2014) (v0.4.15), ggplot2 (Wickham 2016) (v3.4.4), profileplyr (v1.14.1) (https://bioconductor.org/packages/release/bioc/html/profileplyr.html), ViewBS (Huang et al. 2018) (v0.1.11), or pheatmap (v 1.0.12) (https://rdocumentation.org/packages/pheatmap/versions/1.0.12).

#### Raw data and source files

An inventory of all the raw data used to generate each figure panel (as well as those in Supplementary information) along with all raw image files (including RNA, protein blots and agarose gels), DNA and protein sequences, alignments, and phylogenetic tree files can be found in Zenodo (www.zenodo.org) under the doi: 10.5281/zenodo.14037413. Additionally, ready-to-use IGV genome browser tracks for all the sequencing datasets generated here are available as well in the associated Zenodo dataset.

## Data access

All the sequencing data generated for this study have been submitted to the NCBI BioProject database (https://www.ncbi.nlm.nih.gov/bioproject/) under accession number PRJNA1164696. The mass spectrometry proteomics data have been deposited to the ProteomeXchange Consortium via the PRIDE partner repository (https://www.ebi.ac.uk/pride/) with the dataset identifier PXD050443.

## Acknowledgements

This work was supported by the Gregor Mendel Institute of Plant Molecular Biology (GMI) of the Austrian Academy of Sciences (ÖAW) core funding attributed to Arturo Marí-Ordóñez. We thank current and past members of the Marí-Ordóñez group and colleagues from the Gregor Mendel Institute and the Vienna BioCenter (VBC) campus for fruitful discussions, ideas, and feedback. We would also like to thank Klaus Appenroth from the Friedrich Schiller University of Jena for useful advice on duckweed cultivation and storage, Thomas Grentzinger for technical assistance with the TraPR procedure, and Marco Incarbone from the Max-Planck-Institute of Plant Physiology in Golm for kindly providing the pAt*UBQ*:*hpScarlet* plasmid. We are grateful to the Vienna BioCenter Core Facilities (VBCF): NGS for DNA, EM-seq, PacBio, RNA-seq and ChIP-seq library preparation and sequencing, including sequencing of small RNA libraries; the IMP/IMBA/GMI Proteomics Facility, in special Ines Steinmacher for sample preparation, mass-spectrometry using the VBCF instrument pool, and Richard Imre and Elisabeth Rothinger for assistance with data analysis of histone PTMs; the Plant Science Facility for assistance with plant work; the IMP Molecular Biology Service for providing reagents and Sanger sequencing. Lastly, we would like to thank the VBC in-house COVID-19 testing facility for enabling a safe working environment during the pandemic.

## Author contributions

AM-O conceived and designed the study. RD, VB-B and DB-A performed most experiments. AM-O, RD and D-AB performed identification and phylogenetic analysis of silencing components in *Spirodela*. VB-B cloned and carried out transient *AGO4* expression experiments and prepared small RNA libraries. DB-A made nuclei DAPI staining. JMV-G performed western blot analysis of HPTMs. AM-O, DB-A, and AP-M optimized agroinfiltration-mediated transient expression in duckweeds. AP-M carried out transient expression experiments in *Spirodela*. VB-B and RE cultured and maintained *Spirodela* stocks and axenic cultures and provided technical support. RD and DB-A performed computer and statistical analysis. AM-O, RD and DB-A analyzed the data and wrote the manuscript. AMO, DB-A and LD-N addressed referee’s comments and prepared the revised version of the manuscript.

