## Supplemental Information (Figures and Methods) for "Atypical epigenetic and small RNA control of degenerated transposons and their fragments in clonally reproducing *Spirodela polyrhiza*"

#### Table of contents:

Page
2 **Supplemental Figure 1:** Genetic differences between Sp9509 from RDSC and Landolt collection
3 **Supplemental Figure 2:** Phylogenetic position of *Spirodela* within flowering plants
4 **Supplemental Figure 3:** Small RNA profiles of Arabidopsis WT, *nrpd1* and Spirodela
5 **Supplemental Figure 4:** Gene body methylation in *Spirodela*
6 **Supplemental Figure 5:** List of gene IDs shown in Figure 2
7 **Supplemental Figure 6:** Phylogenetic analysis of *Spirodela polyrhiza* DRB proteins
8 **Supplemental Figure 7:** Phylogenetic analysis of *Spirodela polyrhiza* RDR proteins
9 **Supplemental Figure 8:** Phylogenetic analysis of *Spirodela polyrhiza* DCL proteins
10 **Supplemental Figure 9:** Phylogenetic analysis of *Spirodela polyrhiza* AGO proteins
12 **Supplemental Figure 10:** *Spirodela* #9509 AGO5 cluster
13 **Supplemental Figure 11:** Phylogenetic analysis of *Spirodela polyrhiza* SHH proteins
14 **Supplemental Figure 12:** Phylogenetic analysis of *Spirodela polyrhiza* Snf2 chromatin remodelers
16 **Supplemental Figure 13:** Phylogenetic analysis of *Spirodela polyrhiza* SUVH and SUVR proteins
18 **Supplemental Figure 14:** Phylogenetic analysis of *Spirodela polyrhiza* DNA methyltransferases
19 **Supplemental Figure 15:** Phylogenetic analysis of *Spirodela polyrhiza* RNA Pol. large subunit (NRP) proteins
21 **Supplemental Figure 16:** Phylogenetic analysis of *Spirodela polyrhiza* SPT5 and SPT5L proteins
23 **Supplemental Figure 17:** Phylogenetic analysis of *Spirodela polyrhiza* VIM proteins
24 **Supplemental Figure 18:** Phylogenetic analysis of *Spirodela polyrhiza* SUVH5/6 and ASI1 proteins
25 **Supplemental Figure 19:** Genome-wide distribution of DNA methylation in *Spirodela polyrhiza*
26 **Supplemental Figure 20:** Distribution of CHH DNA methylation in *Spirodela polyrhiza*
27 **Supplemental Figure 21:** Transposon classification, nomenclature and codes used in this study
28 **Supplemental Figure 22:** Transposon composition of *Arabidopsis (Col-0)* and *Spirodela (9509)* genomes
29 **Supplemental Figure 23:** mCHH levels across TE sizes and RLC and RLG length distribution
30 **Supplemental Figure 24:** Mass-spectrometry (MS) quantification of H3K9me1/2 abundance in *Spirodela*
31 **Supplemental Figure 25:** Chromatin immunoprecipitation (ChIP) workflow and quality control in *Spirodela*
32 **Supplemental Figure 26:** Genome-wide distribution of H3K9me1, H3K9me2 and H3K27me3 in *Spirodela*
33 **Supplemental Figure 27:** *Spirodela* chromatin organization in interphase nuclei
34 **Supplemental Figure 28:** Distribution of H3K9me1/2 in *Spirodela* TEs
35 **Supplemental Figure 29:** Relative position of H3K9me1 and H3K9me2 peaks to each other in *Spirodela*
36 **Supplemental Figure 30:** Analysis of H3K9me1/2 association to DNA methylation in *Arabidopsis* and *Spirodela*
37 **Supplemental Figure 31:** Patterns of H3K27me1 and H3K9me1/2 distribution in *Spirodela polyrhiza* transposons
38 **Supplemental Figure 32:** Analysis of the epigenetic and small RNA landscape of *5S rRNA* repeats in *Spirodela*
39 **Supplemental Figure 33:** Distribution of silencing marks and sRNAs in *Arabidopsis* RLC and RLG complete TEs
40 **Supplemental Figure 34:** LTR identification for complete TEs in Figure 6
41 **Supplemental Figure 35:** Defects in *SpAGO4a* splicing in *N. benthamiana* prevent its transient expression
42 **Supplemental Figure 36:** Additional examples of siRNA producing TEs in Spirodela
43 **Supplemental Figure 37:** *TAS3* tasiRNAs in *Spirodela*
44 **Supplemental Figure 38:** Transient expression procedure in *Spirodela*
46 **Supplemental Figure 39:** Oligos used in this study
47 **Supplemental Methods**
58 **References to Supplemental Methods**

### SUPPLEMENTAL FIGURE S1

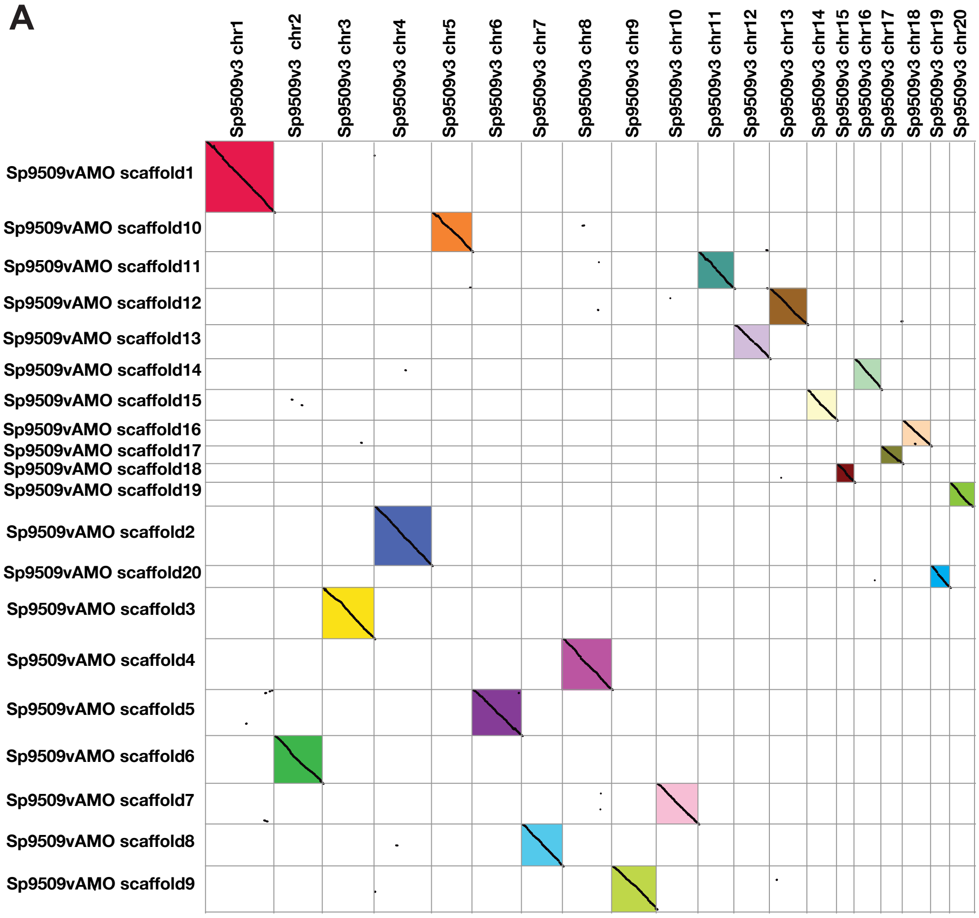

**B**

| Chr | SNPs | INDELs | Mixed |
|---|---|---|---|
| 1 | 37731 | 12583 | 95 |
| 2 | 29937 | 9288 | 68 |
| 3 | 38448 | 11399 | 68 |
| 4 | 29164 | 10081 | 85 |
| 5 | 23655 | 8101 | 66 |
| 6 | 28412 | 10142 | 91 |
| 7 | 26706 | 9233 | 85 |
| 8 | 32969 | 9520 | 70 |
| 9 | 19155 | 7873 | 53 |
| 10 | 33311 | 9768 | 78 |
| 11 | 28157 | 8431 | 65 |
| 12 | 23193 | 8181 | 82 |
| 13 | 19514 | 7285 | 51 |
| 14 | 20716 | 6278 | 28 |
| 15 | 30989 | 7471 | 71 |
| 16 | 24749 | 6791 | 44 |
| 17 | 17239 | 5592 | 56 |
| 18 | 13502 | 6255 | 23 |
| 19 | 22995 | 5061 | 41 |
| 20 | 12333 | 4364 | 44 |
| **Total** | **512875** | **163697** | **1264** |

**Supplemental Figure S1: Genetic differences between Sp9509 from RDSC and Landolt collection.**
**(A)** Syntenic dotplot between Sp9509 from RDSC and Landolt collection, the analysis was generated with SynMap from genomevolution.org using default parameters. **(B)** Summary table of identified variations between Sp9509 from RDSC and Landolt collection, using gatk (version 3.8-1).

### SUPPLEMENTAL FIGURE S2

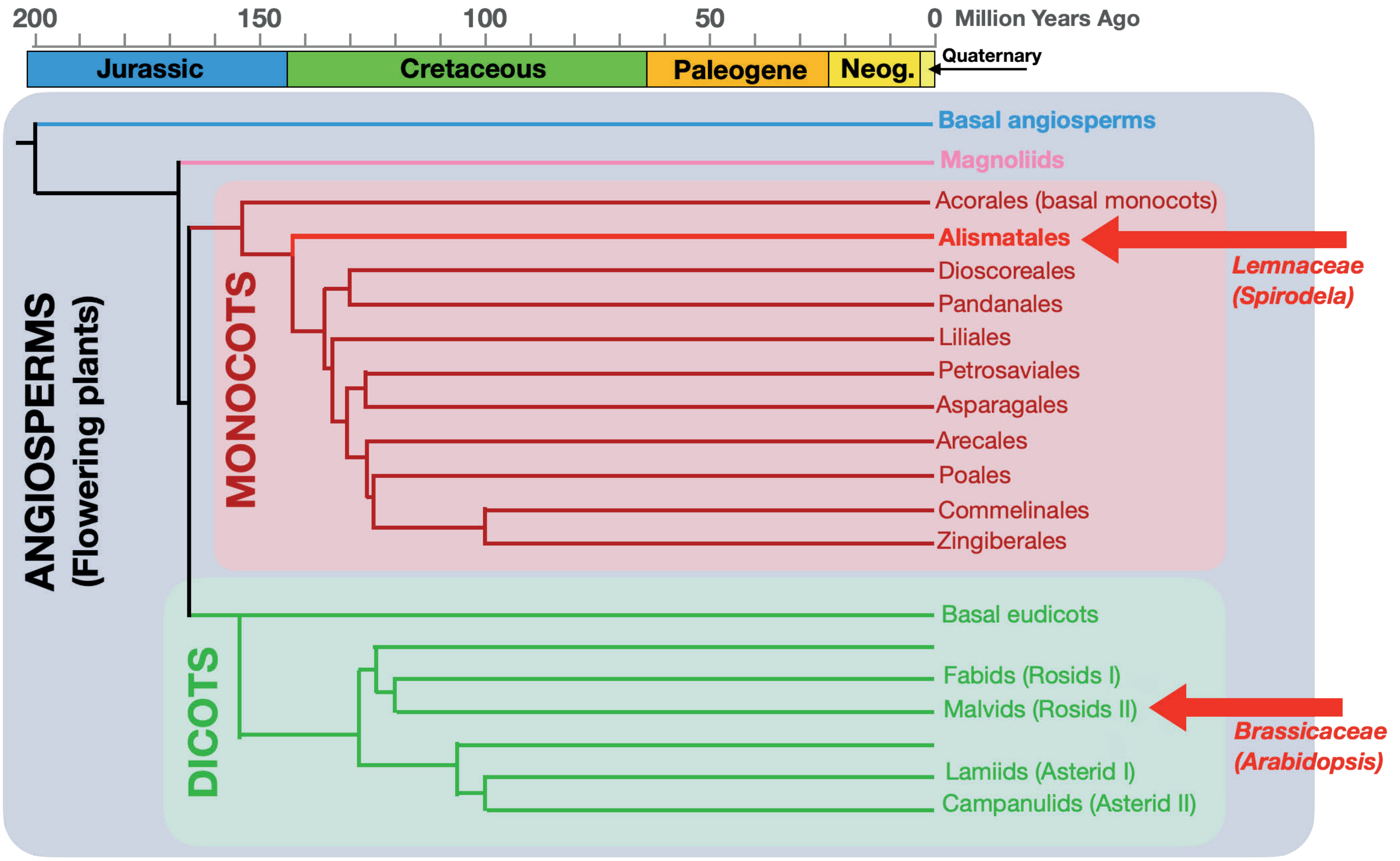

**Supplemental Figure S2: Phylogenetic position of *Spirodela* within flowering plants.**
Summary of the phylogenetic placement of *Spirodela* and *Arabidopsis* within angiosperms.

### SUPPLEMENTAL FIGURE S3

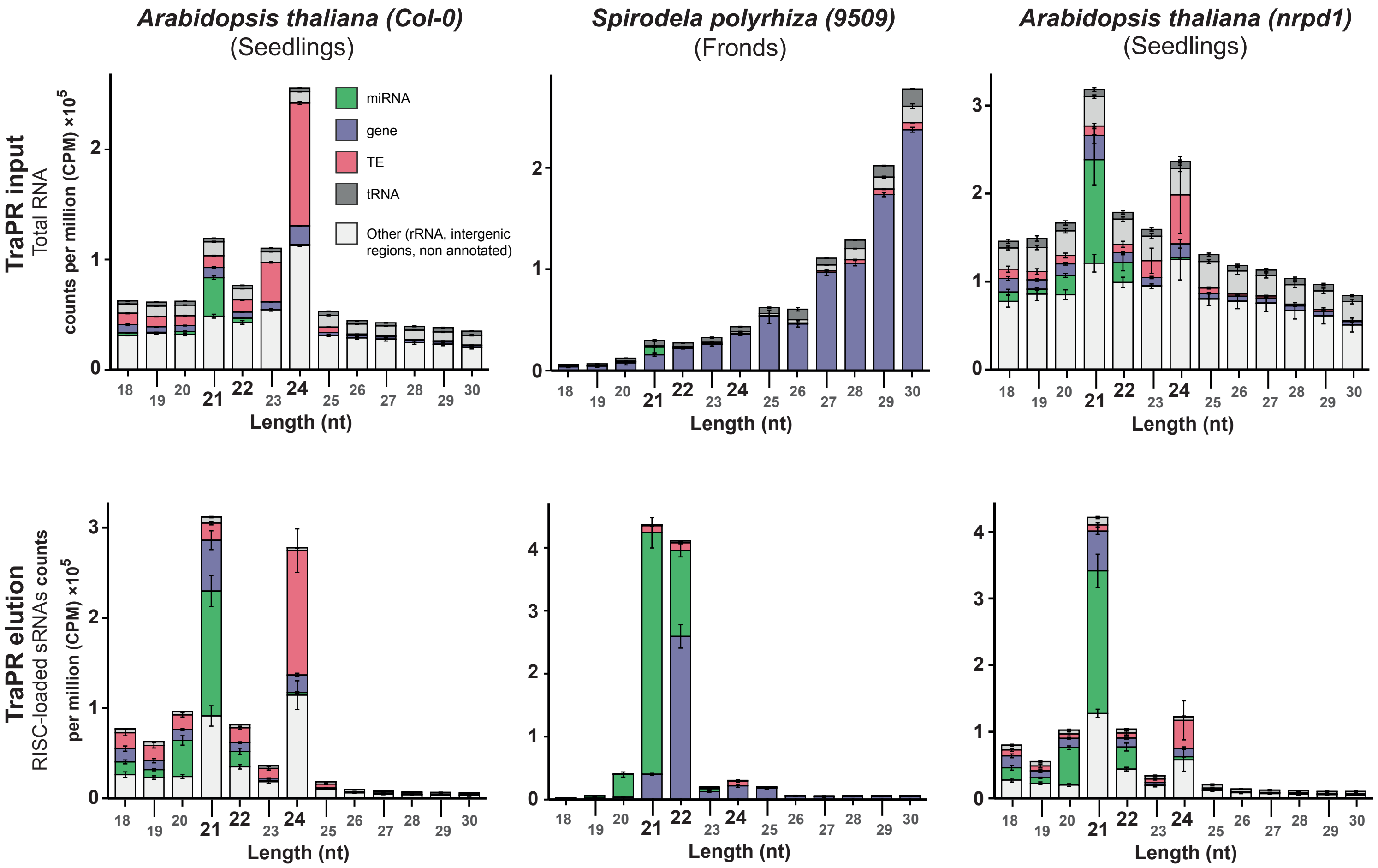

**Supplemental Figure S3: Small RNA profiles of *Arabidopsis* WT, *nrpd1* and *Spirodela*.**
Size distribution and genomic feature annotation of small RNA extracted from TraPR Input (lysate) and elution from *Arabidopsis Col-0* WT, *nrpd1* seedlings and *Spirodela* fronds. Error bars represent standard deviation of the mean from three biological replicates

### SUPPLEMENTAL FIGURE S4

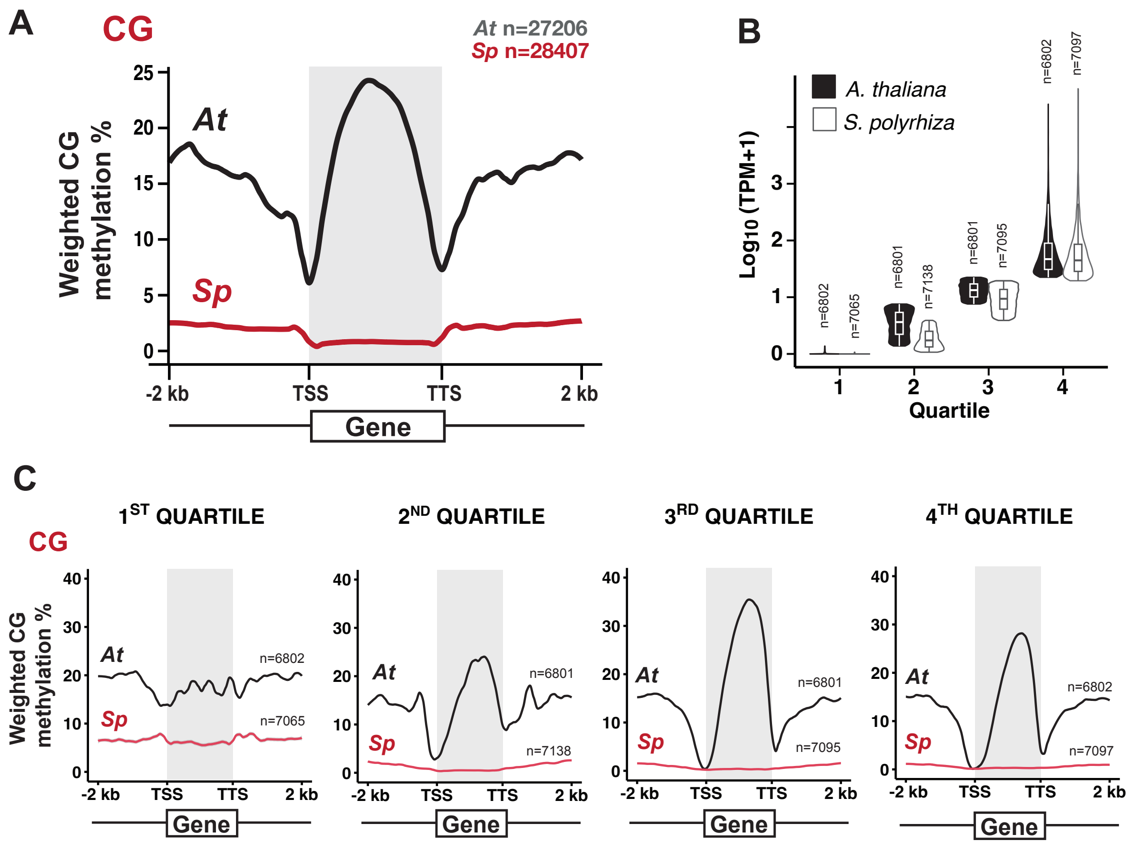

**Supplemental Figure S4: Gene-body Methylation in *Spirodela*.**
**(A)** Metaplots of average weighted CG DNA methylation over protein coding genes and ±2 kb flanking regions in *Arabidopsis* and *Spirodela*. **(B)** Violin and box plots of *Spirodela* and *Arabidopsis* gene expression sorted into quartiles based on transcript levels. In boxplots, median is indicated by solid bar, the boxes extend from the first to the third quartile and whiskers stretch to the furthest value within 1.5 times the interquartile range for each set. **(C)** Metaplots of average weighted CG DNA methylation over protein coding genes and ±2 kb flanking regions in *Arabidopsis* and *Spirodela* within each set of genes sorted by expression quartiles set in (B). *At*: *Arabidopsis*, *Sp*: *Spirodela*.

### SUPPLEMENTAL FIGURE S5

#### miRNA biogenesis

| *Arabidopsis (Col-0)* | | *Spirodela (9509)* | |
|---|---|---|---|
| **GENE** | **AGI** | **GENE** | **Sp9509GI** |
| AtSE | AT2G27100 | SpSE | Sp9509d012g007660 |
| AtHYL1 | AT1G09700 | SpHYL1 | Sp9509d001g003860 |
| AtDRB2 | AT2G28380 | SpDRB2a | Sp9509d002g000320 |
| | | SpDRB2b | Sp9509d014g000150 |
| AtHEN1 | AT4G20910 | SpHEN1 | Sp9509d017g000460 |
| AtDCL1 | AT1G01040 | SpDCL1 | Sp9509d002g008800 |

#### 21-22-nt siRNA biogenesis

| *Arabidopsis (Col-0)* | | *Spirodela (9509)* | |
|---|---|---|---|
| **GENE** | **AGI** | **GENE** | **Sp9509GI** |
| AtSGS3 | AT5G23570 | SpSGS3 | Sp9509d004g006000 |
| AtRDR1 | AT1G14790 | SpRDR1a | Sp9509d018g002090 |
| | | SpRDR1b | Sp9509d018g002070 |
| AtRDR6 | AT3G49500 | SpRDR6 | Sp9509d006g004830 |
| AtRDR3 | AT2G19910 | | |
| AtRDR4 | AT2G19920 | | |
| AtRDR5 | AT2G19930 | | |
| AtDRB4 | AT3G62800 | SpDRB4 | Sp9509d001g004790 |
| AtDRB7.1 | AT1G80650 | | |
| AtDRB7.2 | AT4G00420 | | |
| AtDCL4 | AT5G20320 | SpDCL4 | Sp9509d005g008560 |
| AtDCL2 | AT3G03300 | | |
| RTL1 | AT4G15417 | | |

#### Argonautes

| *Arabidopsis (Col-0)* | | *Spirodela (9509)* | |
|---|---|---|---|
| **GENE** | **AGI** | **GENE** | **Sp9509GI** |
| AtAGO1 | AT1G48410 | SpAGO1 | Sp9509d020g001530 |
| AtAGO10 | AT5G43810 | SpAGO10 | Sp9509d008g011640 |
| AtAGO5 | AT2G27880 | SpAGO5a | Sp9509d007g002750 |
| | | SpAGO5b | Sp9509d007g002760 |
| | | SpAGO5c | Sp9509d007g002770 |
| | | SpAGO5d | Sp9509d007g002790 |
| | | SpAGO5e | Sp9509d007g002800 |
| | | SpAGO5f | Sp9509d007g002810 |
| | | SpAGO5g | Sp9509d007g002820 |
| | | SpAGO5h | Sp9509d007g002830 |
| AtAGO2 | AT1G31280 | | |
| AtAGO3 | AT1G31290 | | |
| AtAGO7 | AT1G69440 | SpAGO7 | Sp9509d009g009840 |
| AtAGO4 | AT2G27040 | SpAGO4a | Sp9509d001g011250 |
| | | SpAGO4b | Sp9509d006g000640 |
| AtAGO8 | AT5G21030 | | |
| AtAGO9 | AT5G21150 | | |
| AtAGO6 | AT2G32290 | | |

#### 24-nt siRNA/RdDM

| *Arabidopsis (Col-0)* | | *Spirodela (9509)* | |
|---|---|---|---|
| **GENE** | **AGI** | **GENE** | **Sp9509GI** |
| AtSHH1 | AT1G15215 | | |
| AtCLSY1 | AT3G42670 | | |
| AtCLSY2 | AT5G20420 | | |
| AtCLSY3 | AT1G05490 | SpCLSY3 | Sp9509d011g003360 |
| AtCLSY4 | AT3G24340 | | |
| AtNRPD1 | AT1G63020 | SpNRPD1 | Sp9509d0013g000080 |
| AtRDR2 | AT4G11130 | SpRDR2 | Sp9509d002g008450 |
| AtDCL3 | AT3G43920 | SpDCL3 | Sp9509d005g001240 |
| AtSUVH2 | AT2G33290 | SpSUVH2 | Sp9509d020g001160 |
| AtSUVH9 | AT4G13460 | | |
| AtDRD1 | AT2G16390 | SpDRD1 | Sp9509d007g000370 |
| AtNRPE1 | AT2G40030 | SpNRPE1 | Sp9509d003g010520 |
| AtSPT5L | AT5G04290 | SpSPT5L | Sp9509d001g003550 |
| AtDRM2 | AT2G33830 | SpDRM2 | Sp9509d008g003000 |
| AtDRM1 | AT1G28330 | | |
| AtDRM3 | AT3G17310 | | |

#### DNA methylation maintenance

| *Arabidopsis (Col-0)* | | *Spirodela (9509)* | |
|---|---|---|---|
| **GENE** | **AGI** | **GENE** | **Sp9509GI** |
| AtDDM1 | AT5G66750 | SpDDM1 | Sp9509d003g002770 |
| AtMET1 | AT5G49160 | SpMET1 | Sp9509d003g004910 |
| AtVIM1 | AT1G57820 | SpVIM | Sp9509d004g007270 |
| AtVIM2 | AT1G66050 | | |
| AtVIM3 | AT5G39550 | | |
| AtVIM4 | AT1G66040 | | |
| AtVIM5 | AT1G57800 | | |
| AtCMT3 | AT1G69770 | | |
| AtCMT2 | AT4G19020 | SpZMET | Sp9509d003g011020 |
| AtSUVH4 | AT5G13960 | SpSUVH4 | Sp9509d004g011450 |
| AtSUVH5 | AT2G35160 | | |
| AtSUVH6 | AT2G22740 | SpSUVH6(5) | Sp9509d008g010000 |

#### Other

| *Arabidopsis (Col-0)* | | *Spirodela (9509)* | |
|---|---|---|---|
| **GENE** | **AGI** | **GENE** | **Sp9509GI** |
| AtSHH2 | AT3G18380 | SpSHH2a | Sp9509d020g005890 |
| | | SpSHH2b | Sp9509d009g007120 |
| | | SpDRB6 | Sp9509d019g004650 |
| AtASI1 | AT5G11470 | SpASI1 | Sp9509d005g000050 |

**Supplemental Figure S5: List of gene IDs shown in Figure 2.**

### SUPPLEMENTAL FIGURE S6

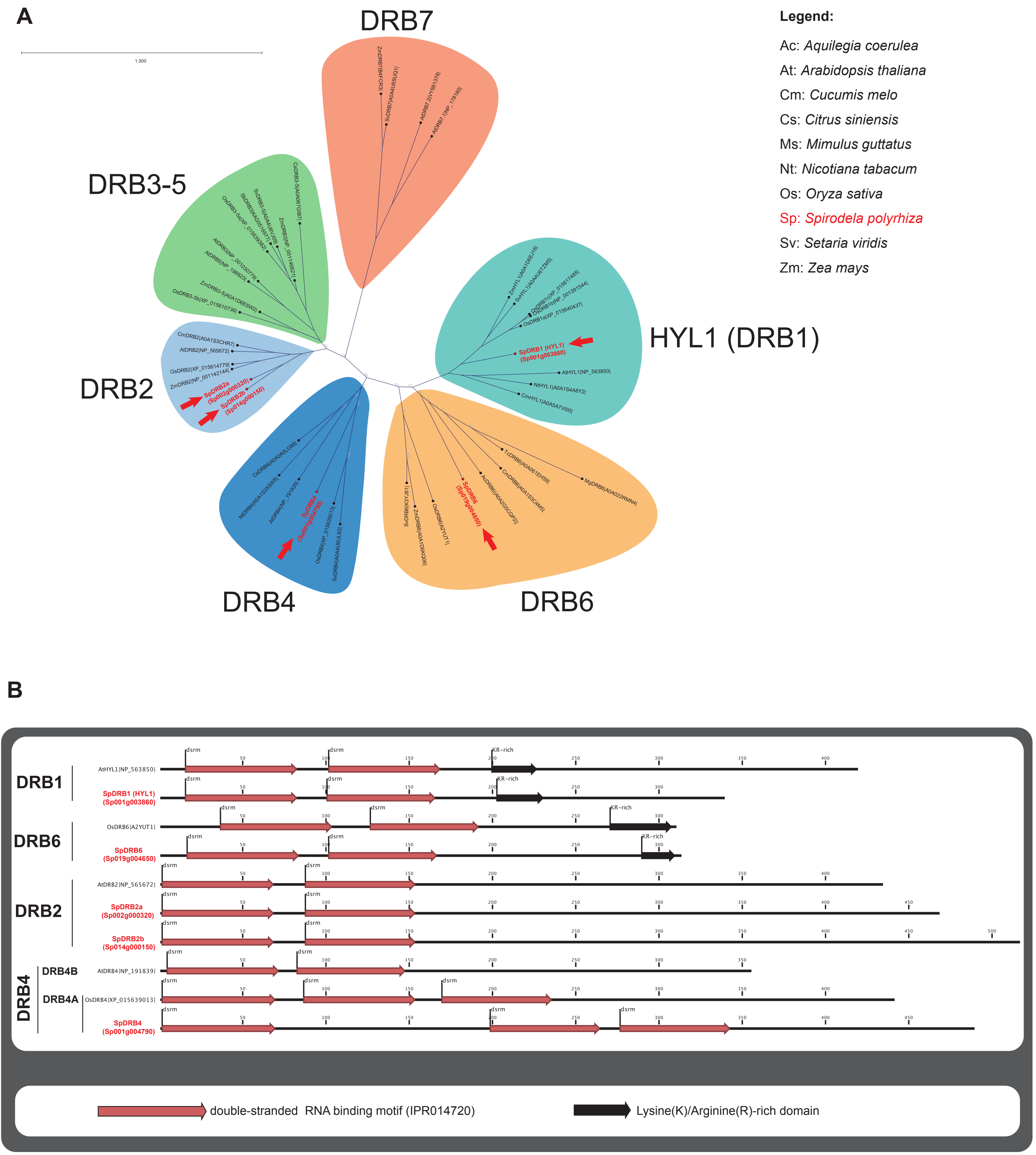

**Supplemental Figure S6: Phylogenetic analysis of *Spirodela polyrhiza* DRB proteins.**
**(A)** Unrooted phylogenetic tree of DRB proteins from several angiosperms including those identified in the *Spirodela polyrhiza* genome. The maximum likelihood tree was constructed with 1000 bootstrap replications. Protein families are highlighted in bubbles of different colors. Arrows point to *Spirodela* proteins. *Spirodela* proteins and annotations were named accordingly to their position in the tree. Uniprot entry codes appear next to the name of proteins in the tree. For *Spirodela*, the gene annotation ID is shown instead. The sequence of all proteins used to build the tree, their alignment and tree files can be found in the raw data files corresponding to this figure. **(B)** Domain annotation of *Spirodela* proteins in (A) along those of their orthologues in other plant species. Numbers along linear protein representations indicate positions in amino acids. Domain annotations were obtained using Interpro and Pfam databases, their integrated InterPro ID codes are display in the graphical legend. Filled arrows indicate significant domain predictions, empty arrows indicate non-significant predictions. For simplicity, Sp9509 gene identifiers have been shortened from Sp9509d(xxx)g(yyyyyy) to Sp(xxx)g(yyyyyy) (where (xxx) is chromosome number, and (yyyyyy) gene number).

### SUPPLEMENTAL FIGURE S7

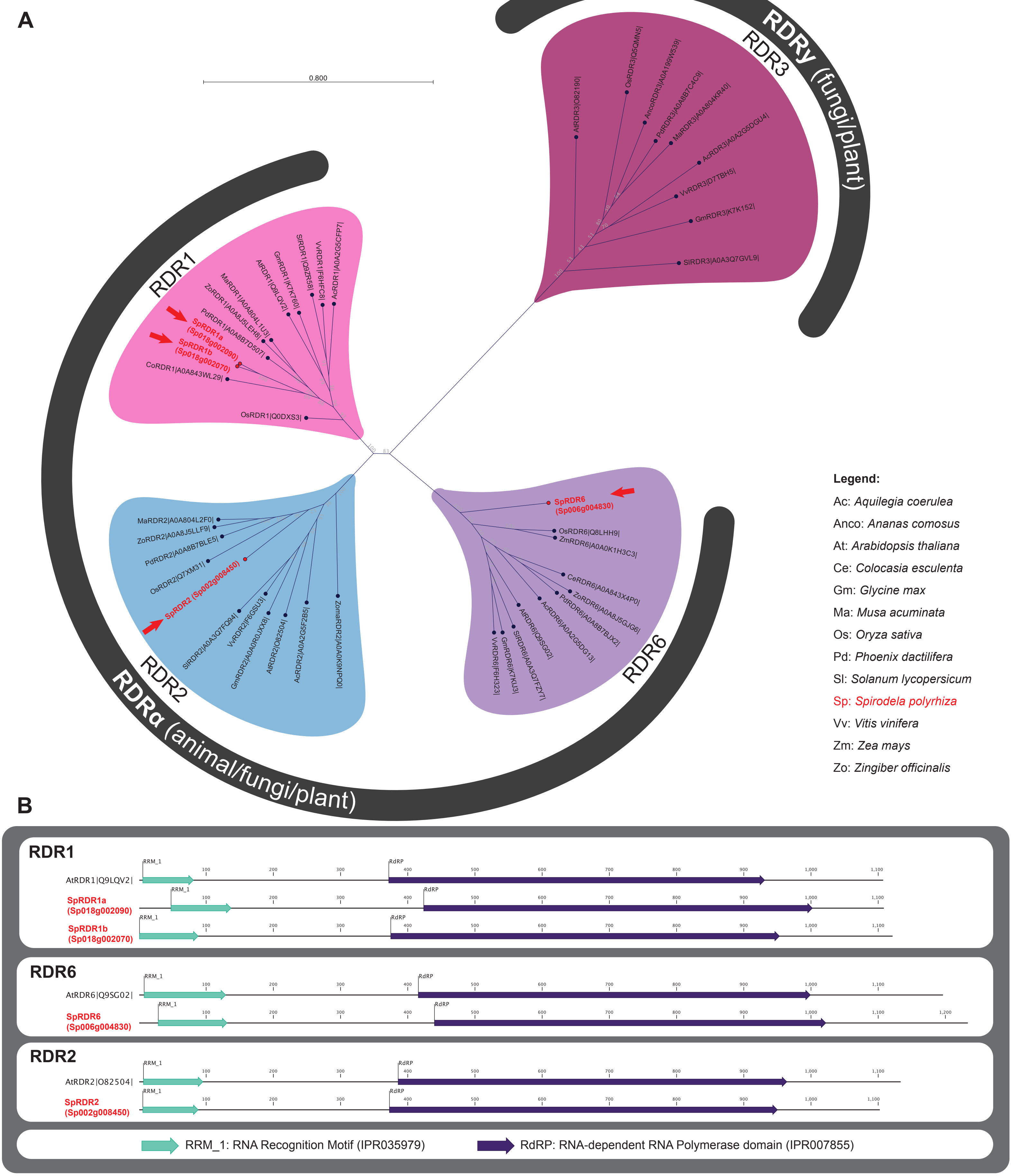

**Supplemental Figure S7: Phylogenetic analysis of *Spirodela polyrhiza* RDR proteins.**
**(A)** Unrooted phylogenetic tree of RNA-dependent RNA Polymerases, RdRPs (RDRs), proteins from several angiosperms, including those identified in the *Spirodela polyrhiza* genome. The maximum likelihood tree was constructed with 1000 bootstrap replications. Protein families are highlighted in bubbles of different colors. Arrows point to *Spirodela* proteins. *Spirodela* proteins and annotations were named accordingly to their position in the tree. Uniprot entry codes appear next to the name of proteins in the tree. For *Spirodela*, the gene annotation ID is shown instead. The sequence of all proteins used to build the tree, their alignment and tree files can be found in the raw data files corresponding to this figure. **(B)** Domain annotation of *Spirodela* proteins in (A) along those of their orthologues in other plant species. Numbers along linear protein representations indicate positions in amino acids. Domain annotations were obtained using Interpro and Pfam databases, their integrated InterPro ID codes are display in the graphical legend. Filled arrows indicate significant domain predictions, empty arrows indicate non-significant predictions. For simplicity, Sp9509 gene identifiers have been shortened from Sp9509d(xxx)g(yyyyyy) to Sp(xxx)g(yyyyyy) (where (xxx) is chromosome number, and (yyyyyy) gene number).

### SUPPLEMENTAL FIGURE S8

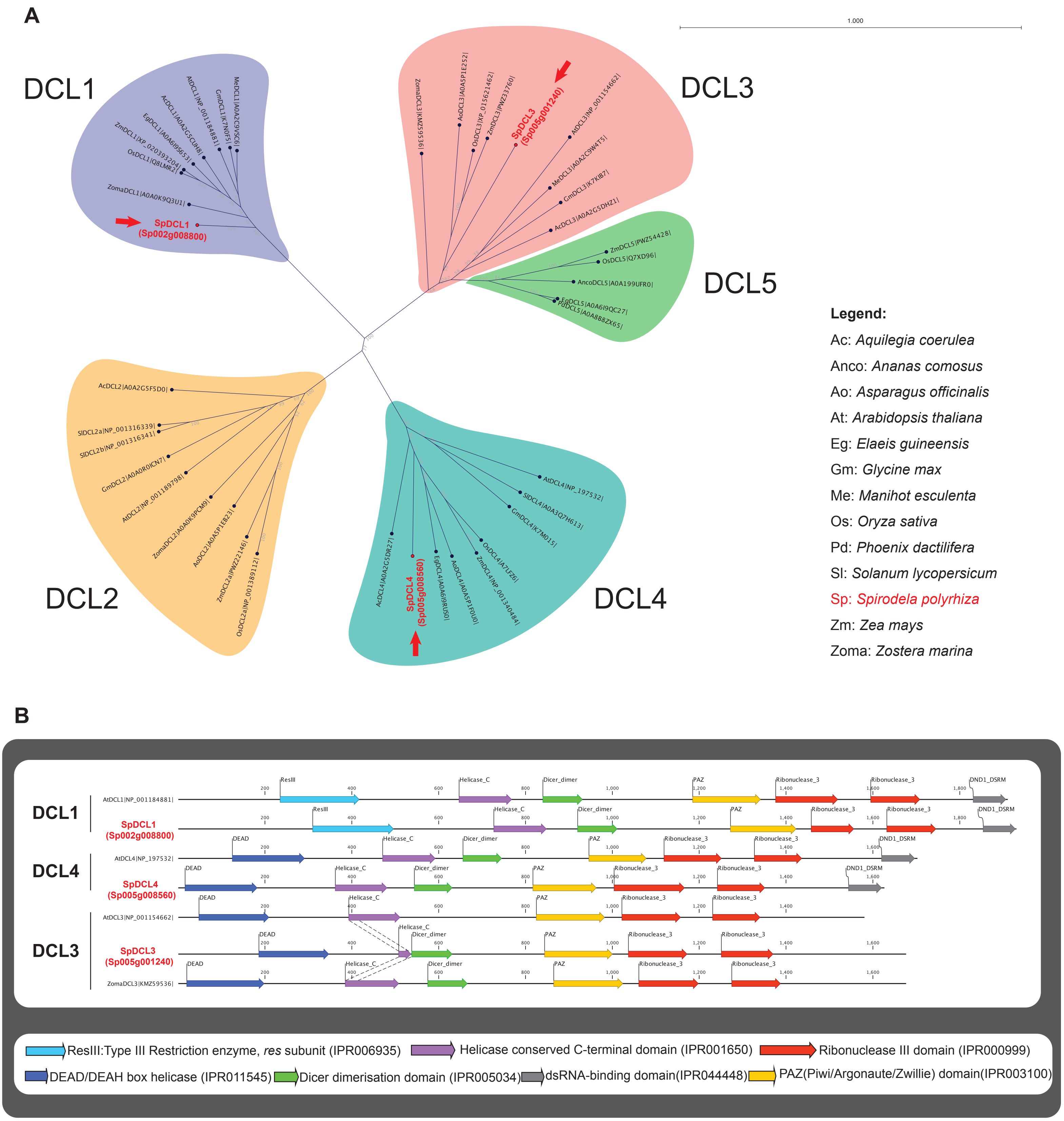

**Supplemental Figure S8: Phylogenetic analysis of *Spirodela polyrhiza* DCL proteins.**
**(A)** Unrooted phylogenetic tree of DCL proteins from several angiosperms including those identified in the *Spirodela polyrhiza* genome. The maximum likelihood tree was constructed with 1000 bootstrap replications. Protein families are highlighted in bubbles of different colors. Arrows point to *Spirodela* proteins. *Spirodela* proteins and annotations were named accordingly to their position in the tree. Uniprot entry codes appear next to the name of proteins in the tree. For *Spirodela*, the gene annotation ID is shown instead. The sequence of all proteins used to build the tree, their alignment and tree files can be found in the raw data files corresponding to this figure. **(B)** Domain annotation of *Spirodela* proteins in (A) along those of their orthologues in other plant species. Numbers along linear protein representations indicate positions in amino acids. Domain annotations were obtained using Interpro and Pfam databases, their integrated InterPro ID codes are display in the graphical legend. Filled arrows indicate significant domain predictions, empty arrows indicate non-significant predictions. For simplicity, Sp9509 gene identifiers have been shortened from Sp9509d(xxx)g(yyyyyy) to Sp(xxx)g(yyyyyy) (where (xxx) is chromosome number, and (yyyyyy) gene number).

### SUPPLEMENTAL FIGURE S9

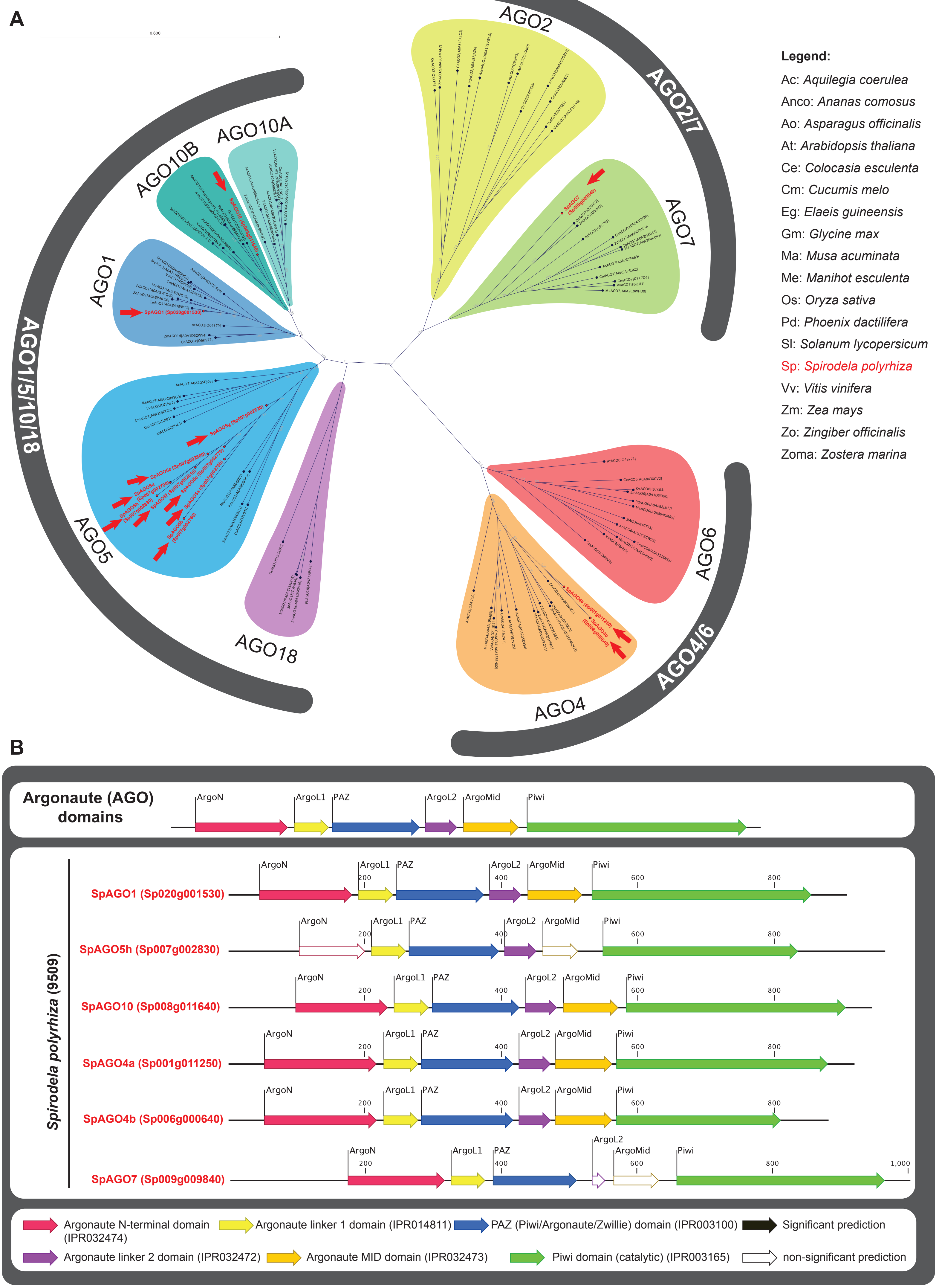

Legend on next page

**Supplemental Figure S9: Phylogenetic analysis of *Spirodela polyrhiza* AGO proteins.**

**(A)** Unrooted phylogenetic tree of AGO proteins from several angiosperms including those identified in the *Spirodela polyrhiza* genome. The maximum likelihood tree was constructed with 1000 bootstrap replications. Protein families are highlighted in bubbles of different colors. Arrows point to *Spirodela* proteins. *Spirodela* proteins and annotations were named accordingly to their position in the tree. Uniprot entry codes appear next to the name of proteins in the tree. For *Spirodela*, the gene annotation ID is shown instead. The sequence of all proteins used to build the tree, their alignment and tree files can be found in the raw data files corresponding to this figure. **(B)** Domain annotation of *Spirodela* proteins in (A) along those of their orthologues in other plant species. Numbers along linear protein representations indicate positions in amino acids. Domain annotations were obtained using Interpro and Pfam databases, their integrated InterPro ID codes are display in the graphical legend. Filled arrows indicate significant domain predictions, empty arrows indicate non-significant predictions. For simplicity, Sp9509 gene identifiers have been shortened from Sp9509d(xxx)g(yyyyyy) to Sp(xxx)g(yyyyyy) (where (xxx) is chromosome number, and (yyyyyy) gene number).

### SUPPLEMENTAL FIGURE S10

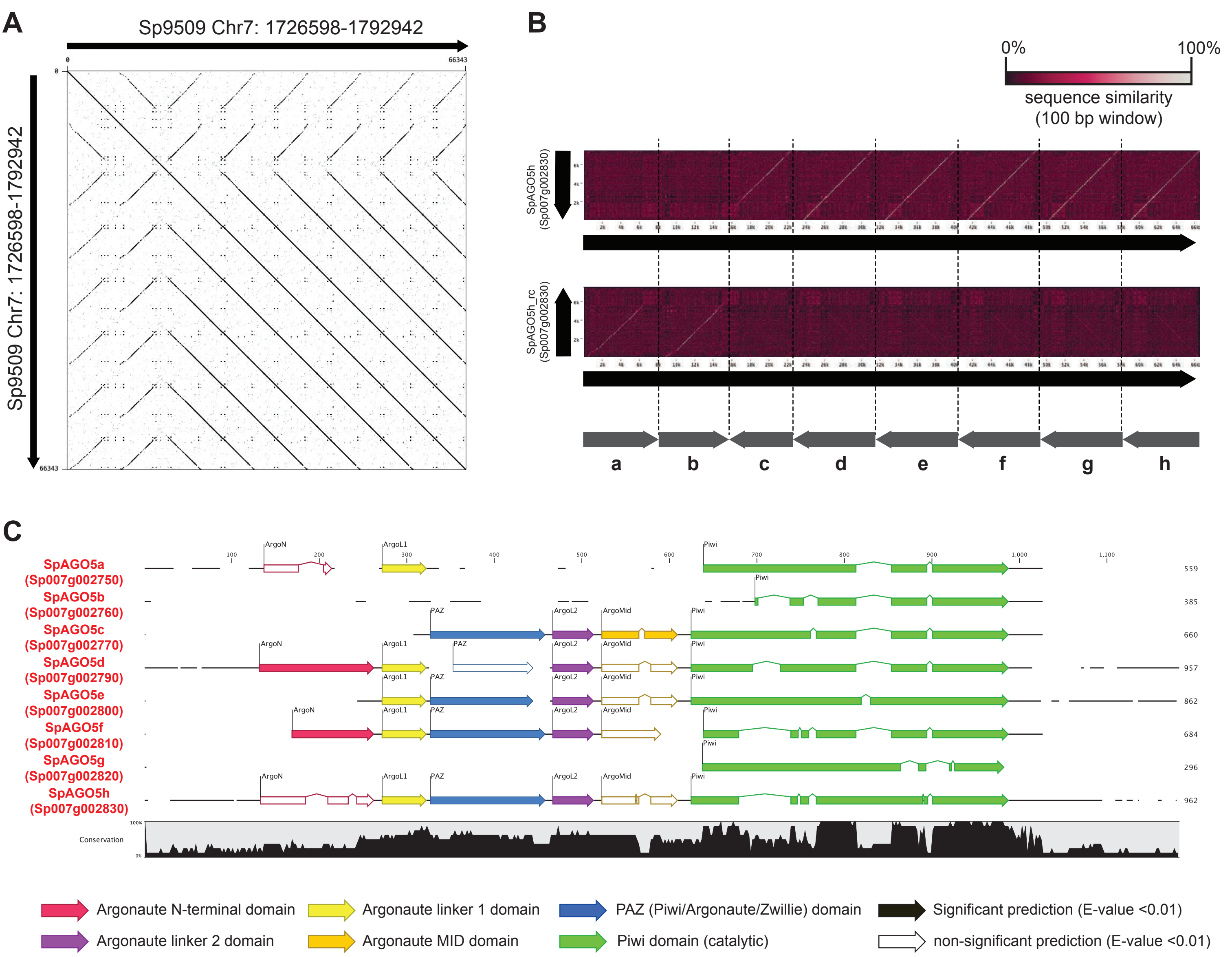

**Supplemental Figure S10: *Spirodela* #9509 AGO5 cluster.**
**(A)** Dot plot of the genomic region in Chromosome 7 containing the eight copies of *AGO5* identified in the *Spirodela polyrhiza* #9509 genome (see Supplementary Figure S9). The dot plot was generated using Gepard 2.0 with 100 bp window as word size. The plot shows the presence of eight tandem duplications, two in sense and six in antisense. **(B)** Sequence similarity dot plot of the same genomic region as in (A) against the DNA sequence of the longest *AGO5* copy (*SpAGO5h*) in both sense and antisense using a 100 bp sliding window. High sequence similarity regions were used to define an average 8 kbp region containing each of the identified *AGO5* copies that has been duplicated. **(C)** Protein alignment of all eight AGO5 copies including domain prediction annotations suggesting that many of the AGO5 copies might not be functional. Domain annotation of *Spirodela AGO5* proteins. Numbers along linear protein representations indicate positions in amino acids. Domain anno-tations were obtained using Interpro and Pfam databases. Filled arrows indicate significant domain predictions, empty arrows indicate non-significant predictions. For simplicity, Sp9509 gene identifiers have been shortened from Sp9509d(xxx)g(yyyyyy) to Sp(xxx)g(yyyyyy) (where (xxx) is chromosome number, and (yyyyyy) gene number).

### SUPPLEMENTAL FIGURE S11

**A**

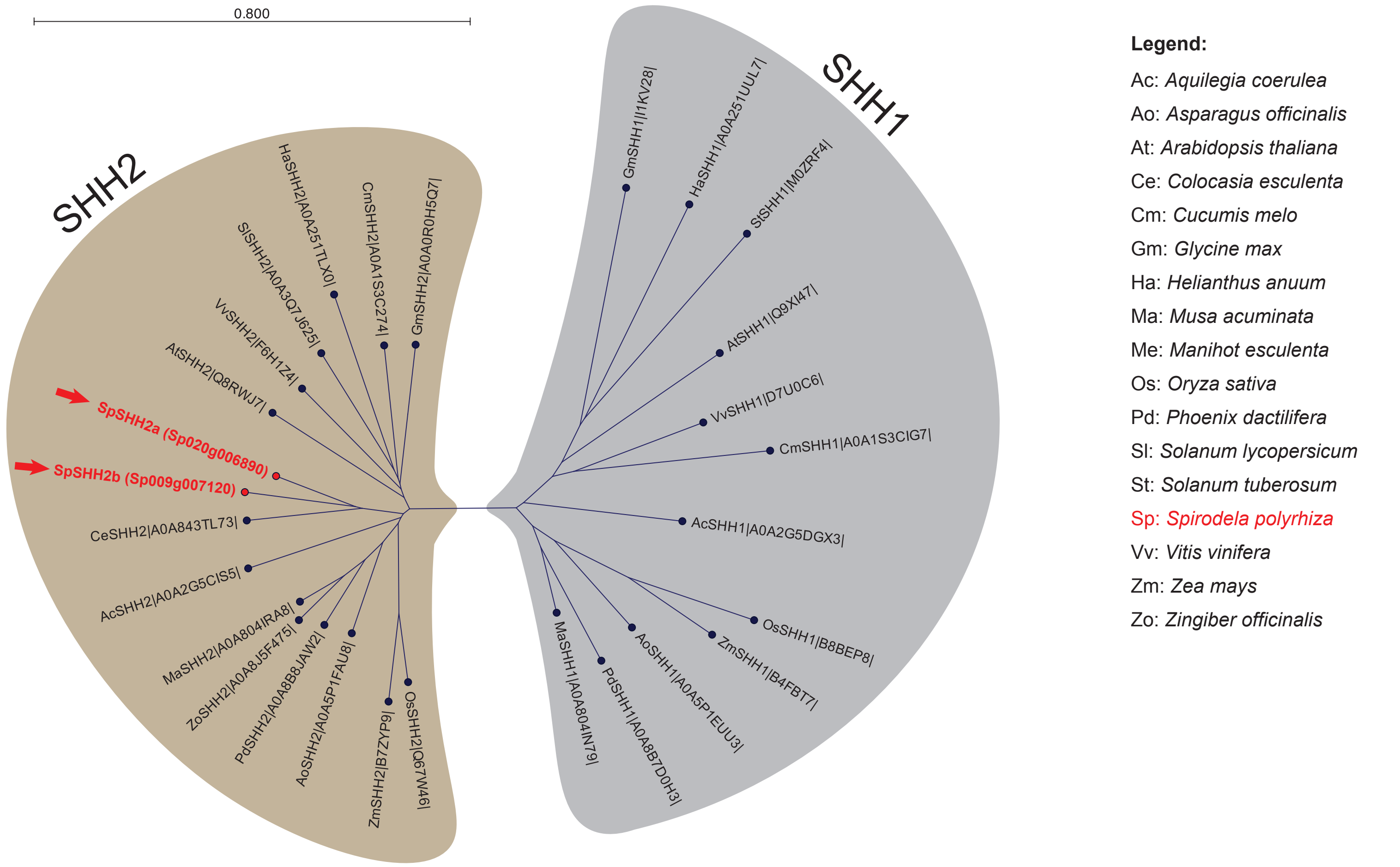

**B**

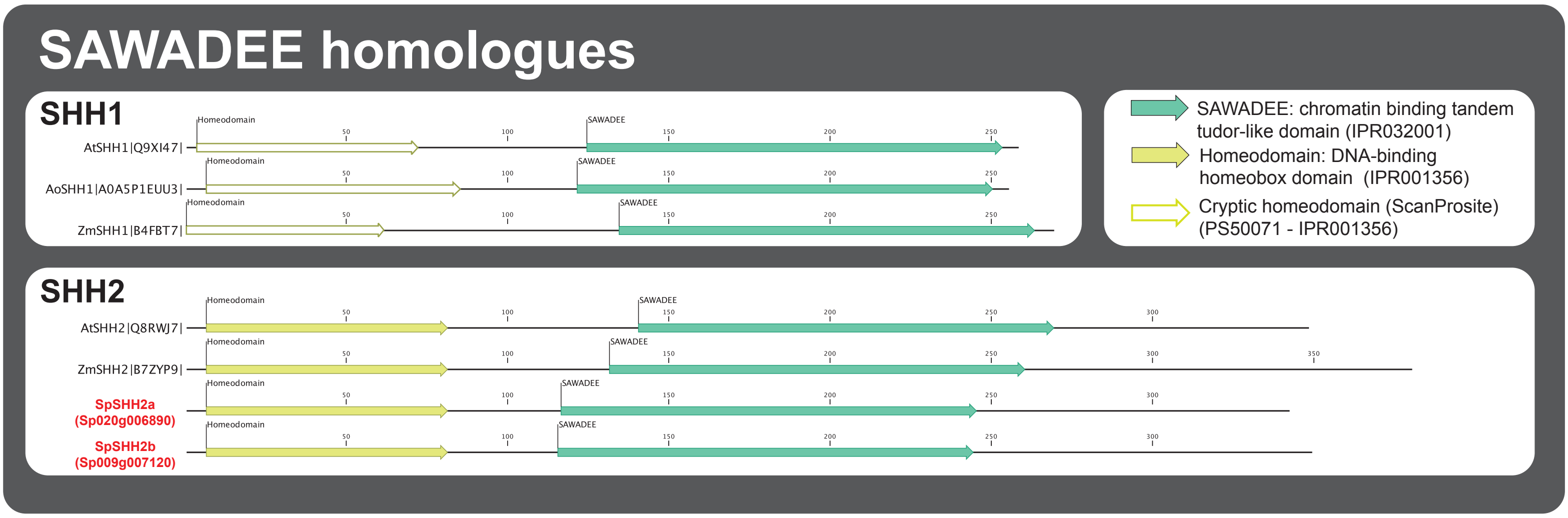

**Supplemental Figure S11: Phylogenetic analysis of *Spirodela polyrhiza* SHH proteins.**
**(A)** Unrooted phylogenetic tree of SHH proteins from several angiosperms including those identified in the *Spirodela polyrhiza* genome. The maximum likelihood tree was constructed with 1000 bootstrap replications. Protein families are highlighted in bubbles of different colors. Arrows point to *Spirodela* proteins. *Spirodela* proteins and annotations were named accordingly to their position in the tree. Uniprot entry codes appear next to the name of proteins in the tree. For *Spirodela*, the gene annotation ID is shown instead. The sequence of all proteins used to build the tree, their alignment and tree files can be found in the raw data files corresponding to this figure. **(B)** Domain annotation of *Spirodela* proteins in (A) along those of their orthologues in other plant species. Numbers along linear protein representations indicate positions in amino acids. Domain annotations were obtained using Interpro, Pfam and Prosite databases, their integrated InterPro ID codes are display in the graphical legend. The cryptic homeodomain found within SHH1 was only identified using ScanProsite. Filled arrows indicate significant domain predictions, empty arrows indicate non-significant predictions. For simplicity, Sp9509 gene identifiers have been shortened from Sp9509d(xxx)g(yyyyyy) to Sp(xxx)g(yyyyyy) (where (xxx) is chromosome number, and (yyyyyy) gene number).

### SUPPLEMENTAL FIGURE S12

**A**

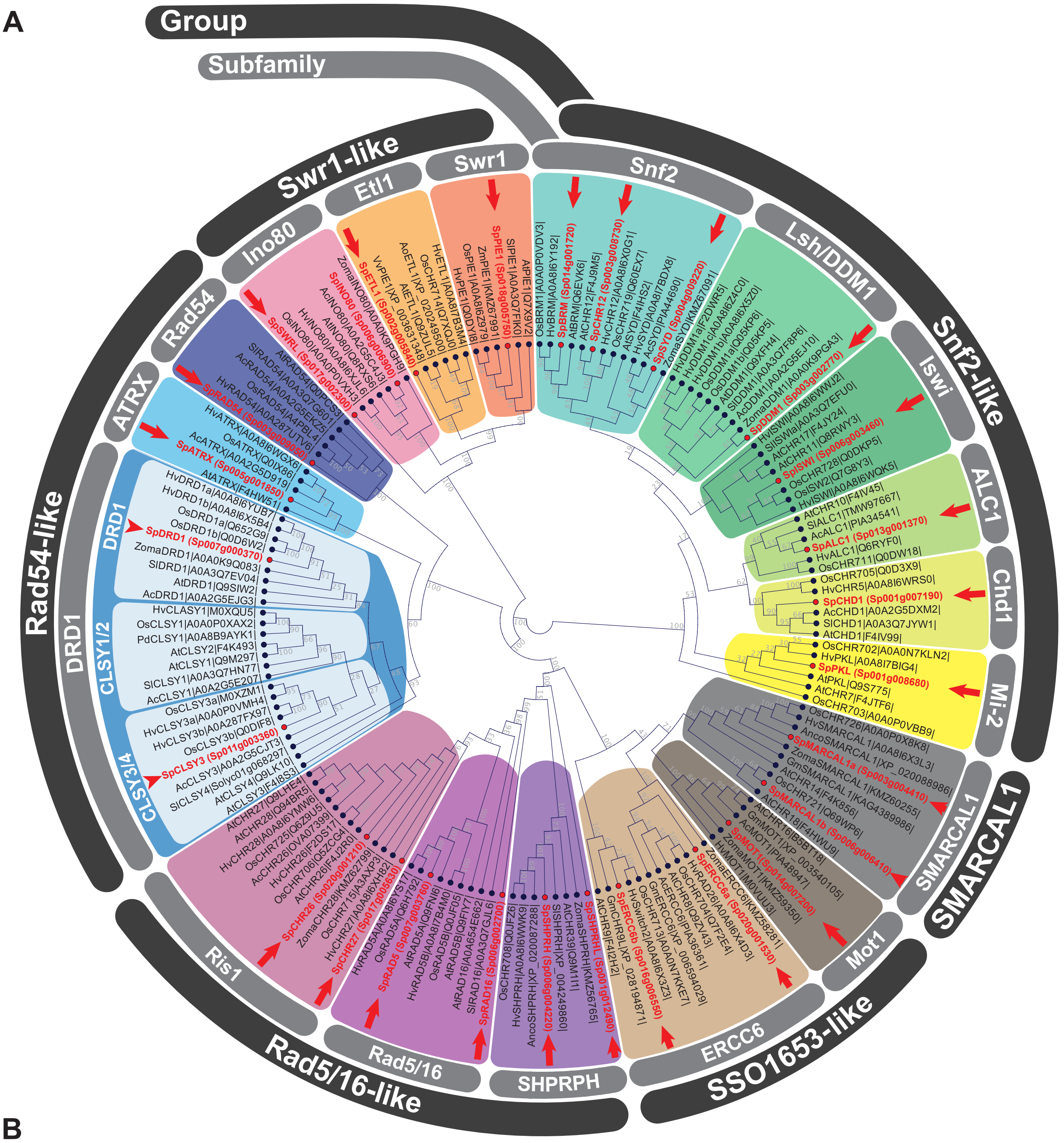

**Legend:**

Ac: *Aquilegia coerulea*
Anco: *Ananas comosus*
Ao: *Asparagus officinalis*
At: *Arabidopsis thaliana*
Ce: *Colocasia esculenta*
Gm: *Glycine max*
Hv: *Hordeum vulgare*
Os: *Oryza sativa*
Pd: *Phoenix dactilifera*
Sl: *Solanum lycopersicum*
Sp: *Spirodela polyrhiza*
Vv: *Vitis vinifera*
Zm: *Zea mays*
Zoma: *Zostera marina*

**B**

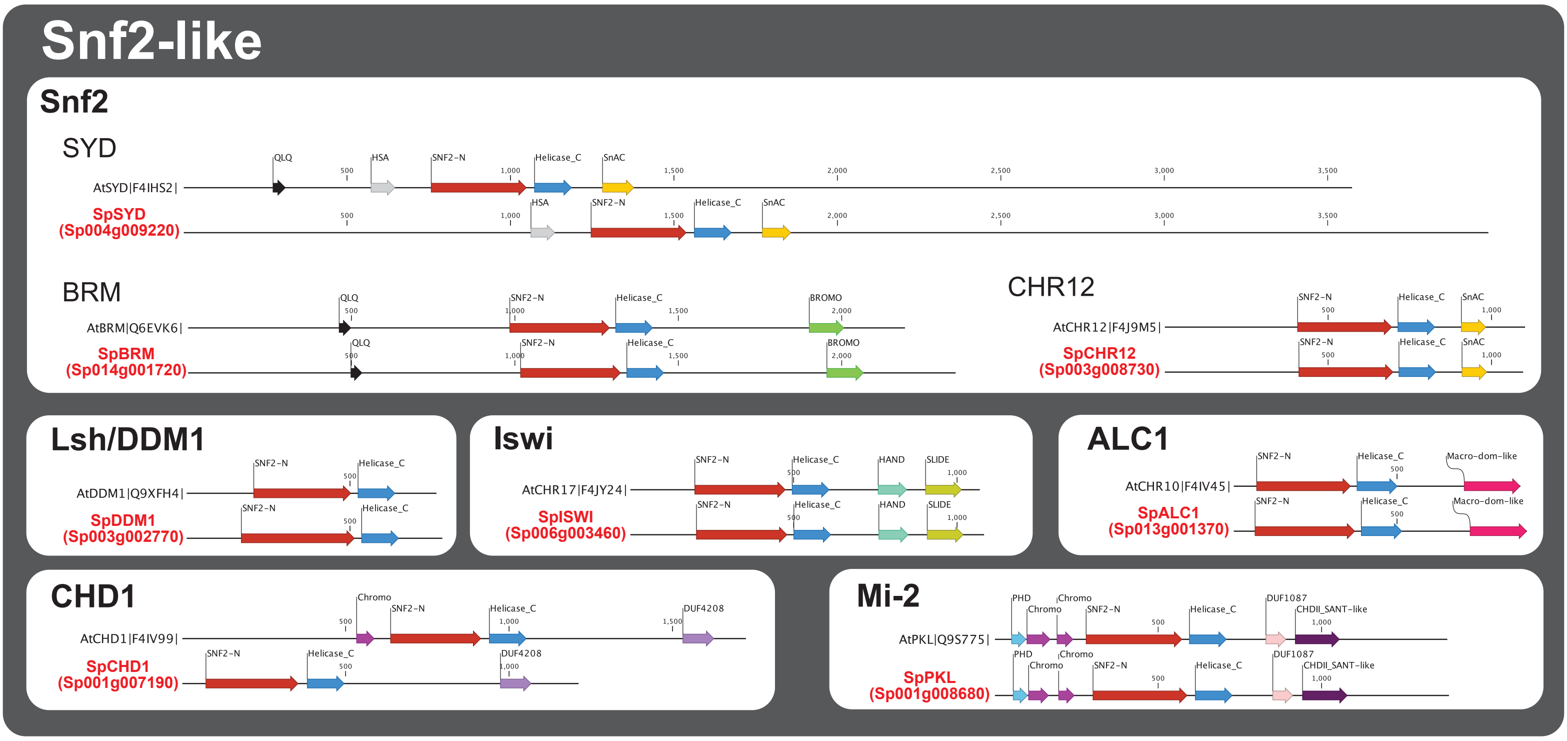

Continues on next page (incl. legend)

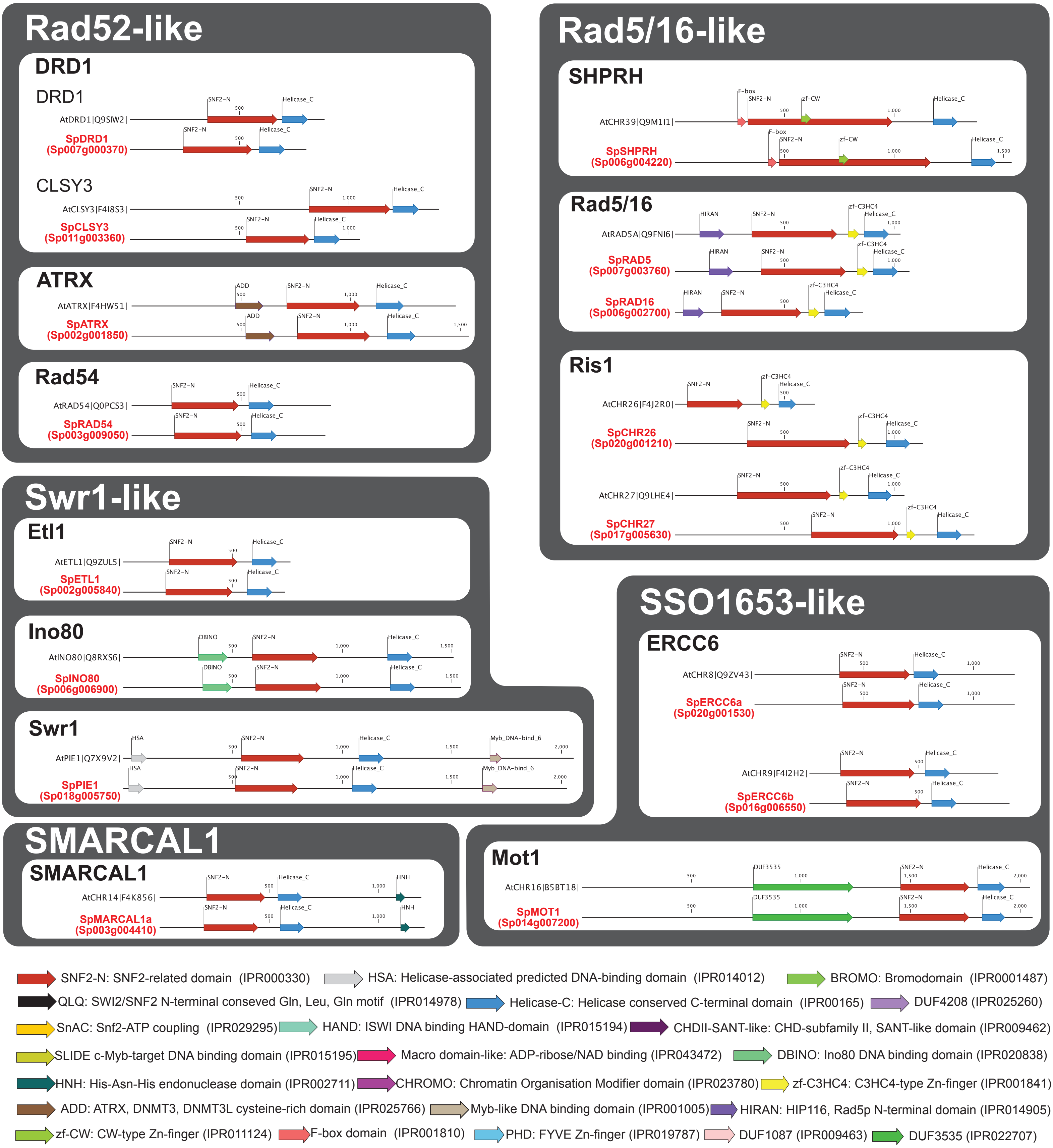

**Supplemental Figure S12: Phylogenetic analysis of *Spirodela polyrhiza* Snf2 chromatin remodelers.** **(A)** Unrooted phylogenetic tree of angiosperm Snf2 ATP-dependent chromatin remodelers from several species including those identified in the *Spirodela polyrhiza* genome. The maximum likelihood tree was constructed with 1000 bootstrap replications. Protein subfamilies are highlighted in bubbles of different colors. Spirodela proteins and annotations were named accordingly to their position in the tree. Uniprot entry codes appear next to the name of proteins in the tree. For *Spirodela*, the gene annotation ID is shown instead. Arrows point to *Spirodela* proteins. The sequence of all proteins used to build the tree, their alignment and tree files can be found in the raw data files corresponding to this figure. **(B)** Domain annotation of *Spirodela* proteins in (A) along those of their orthologues in other plant species. Numbers along linear protein representations indicate positions in amino acids. Domain annotations were obtained using InterPro and Pfam databases, their integrated InterPro ID codes are display in the graphical legend. Filled arrows indicate significant domain predictions, empty arrows indicate non-significant predictions. For simplicity, Sp9509 gene identifiers have been shortened from Sp9509d(xxx)g(yyyyyy) to Sp(xxx)g(yyyyyy) (where (xxx) is chromosome number, and (yyyyyy) gene number).

### SUPPLEMENTAL FIGURE S13

## A

**Legend:**

At: *Arabidopsis thaliana*
Cs: *Citrus siniensis*
Gh: *Gossypium hirsutum*
Gm: *Glycine max*
Ha: *Helianthus annuus*
Hv: *Hordeum vulgare*
Me: *Manihot esculenta*
Os: *Oryza sativa*
Sl: *Solanum lycopersicum*
St: *Solanum tuberosum*
Sb: *Sorghum bicolor*
Sp: *Spirodela polyrhiza*
Zm: *Zea mays*
Zo: *Zingiber officinalis*

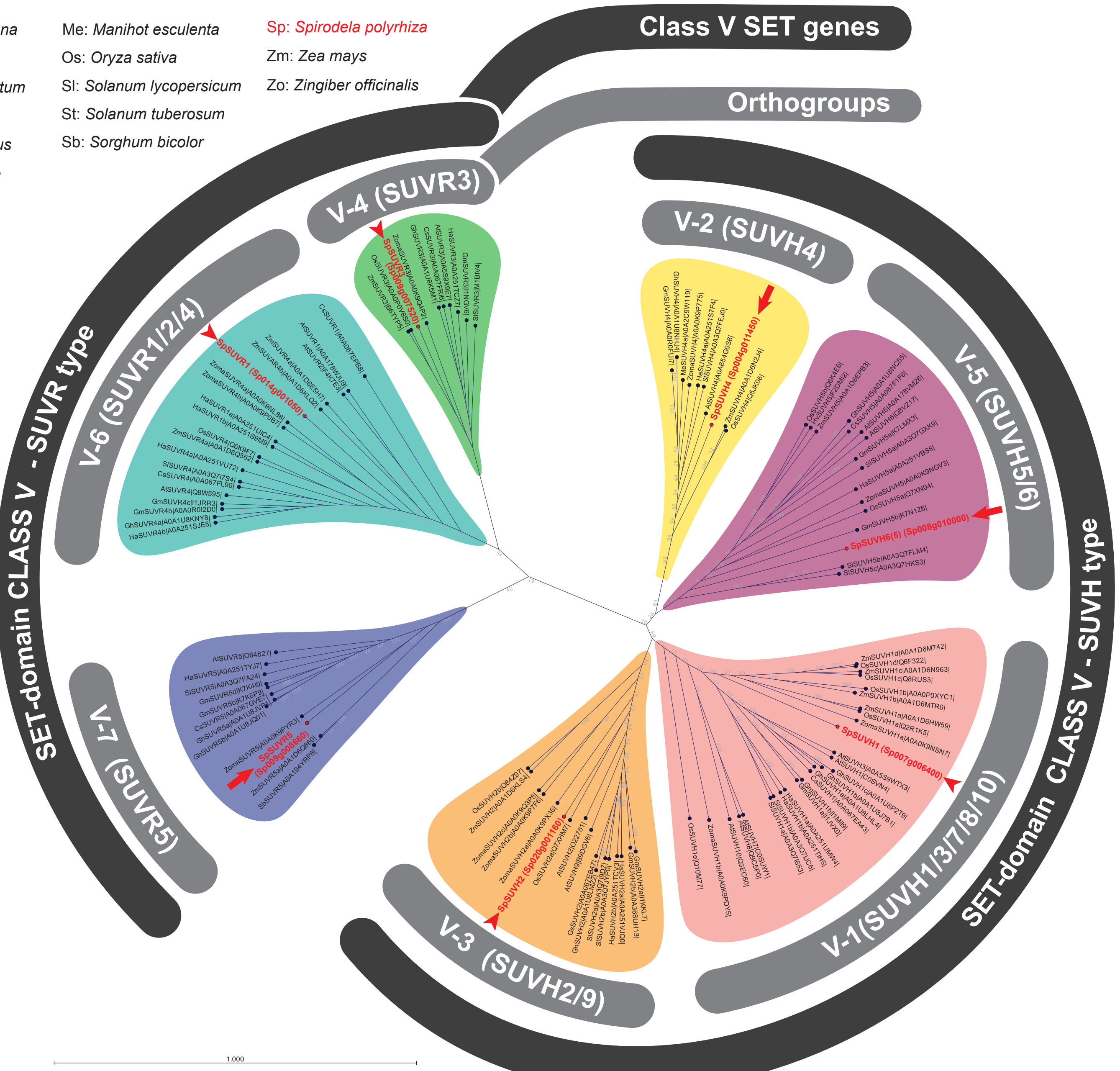

## B

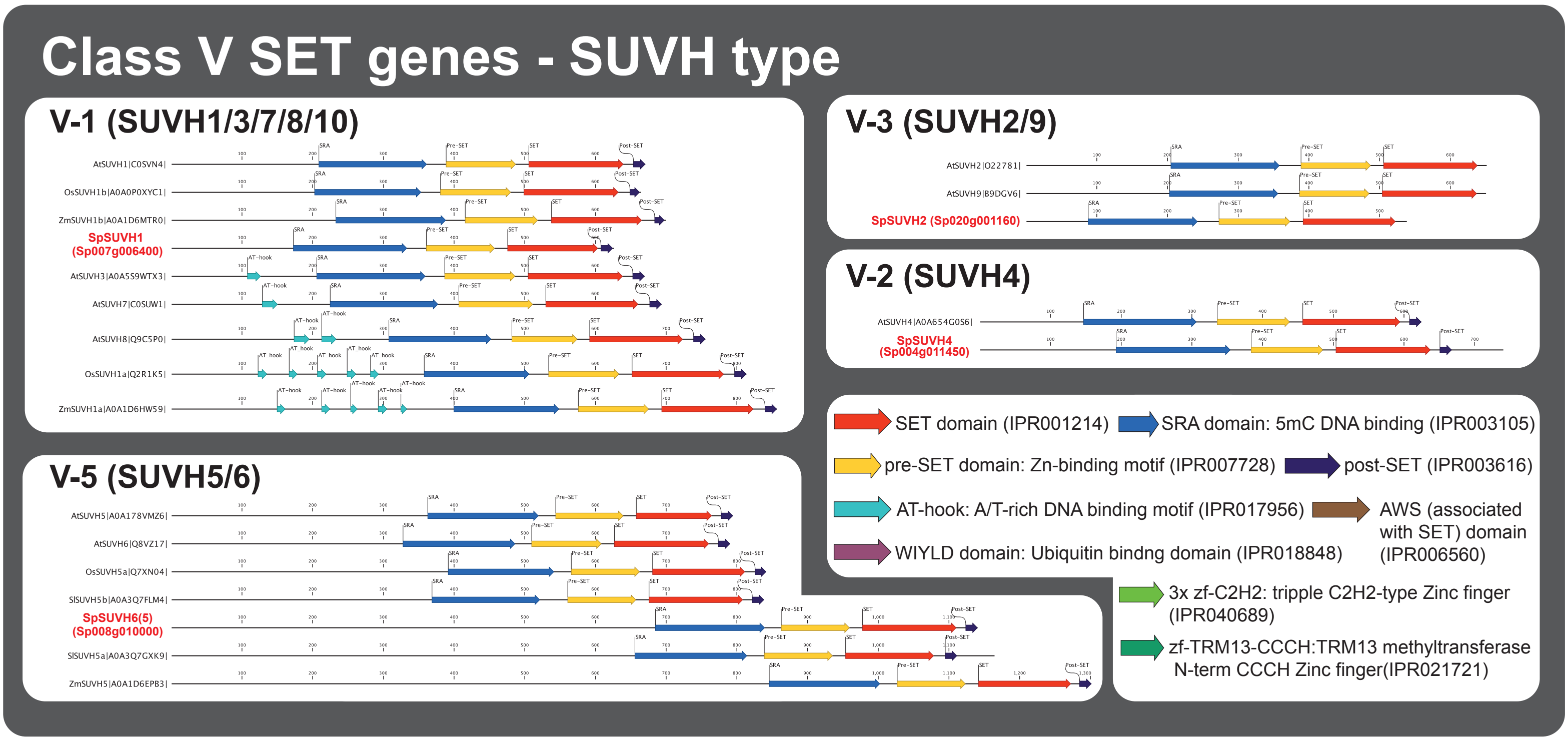

Continues on next page (incl. legend)

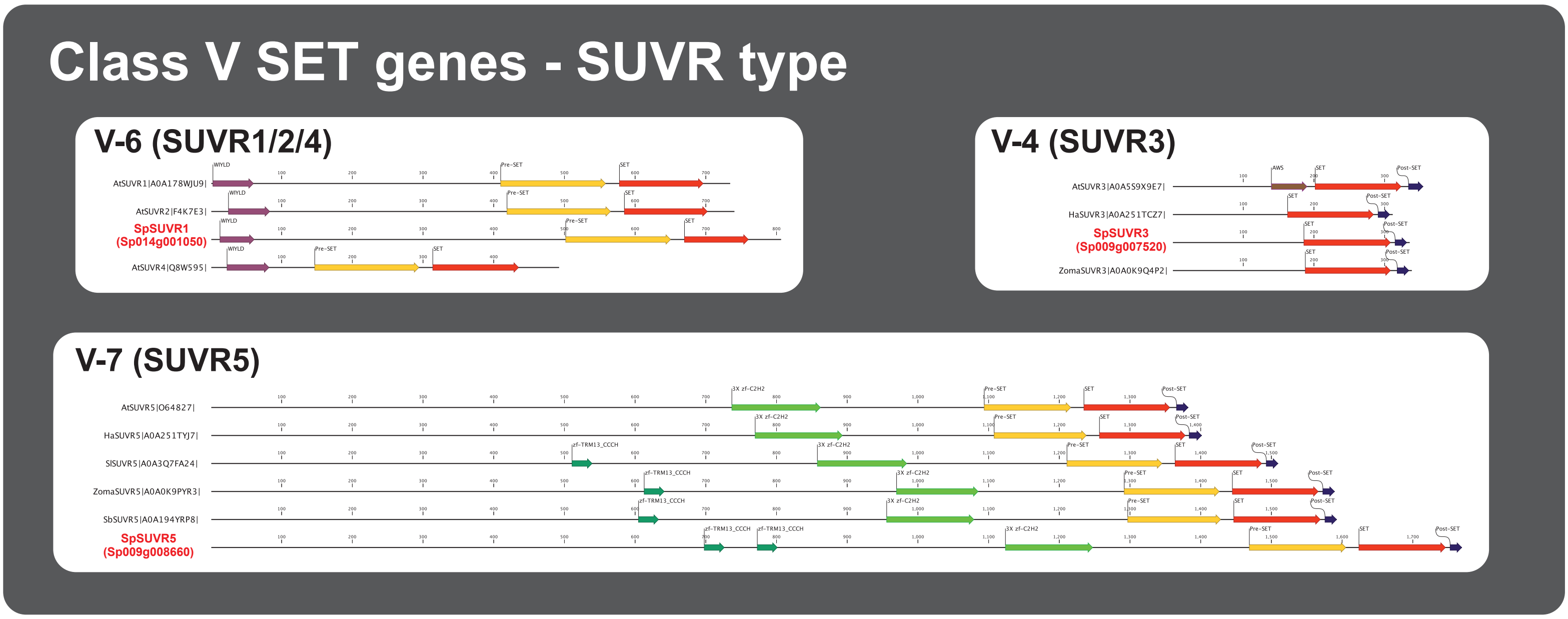

**Supplemental Figure S13: Phylogenetic analysis of *Spirodela polyrhiza* SUVH and SUVR proteins.**
**(A)** Unrooted phylogenetic tree of angiosperm Class V SET domain-containing proteins, Su(var)3-9 homologues and relatives (SUVH and SUVR), from several species including those identified in the *Spirodela polyrhiza* genome. The maximum likelihood tree was constructed with 1000 bootstrap replications. Protein families are highlighted in bubbles of different colors. *Spirodela* proteins and annotations were named accordingly to their position in the tree. Arrows point to *Spirodela* proteins. Uniprot entry codes appear next to the name of proteins in the tree. For *Spirodela*, the gene annotation ID is shown instead. The sequence of all proteins used to build the tree, their alignment and tree files can be found in the raw data files corresponding to this figure.
**(B)** Domain annotation of *Spirodela* proteins in (A) along those of their orthologues in other plant species. Numbers along linear protein representations indicate positions in amino acids. Domain annotations were obtained using Interpro and Pfam databases, their integrated InterPro ID codes are display in the graphical legend. Filled arrows indicate significant domain predictions, empty arrows indicate non-significant predictions. For simplicity, Sp9509 gene identifiers have been shortened from Sp9509d(xxx)g(yyyyyy) to Sp(xxx)g(yyyyyy) (where (xxx) is chromosome number, and (yyyyyy) gene number).

### SUPPLEMENTAL FIGURE S14

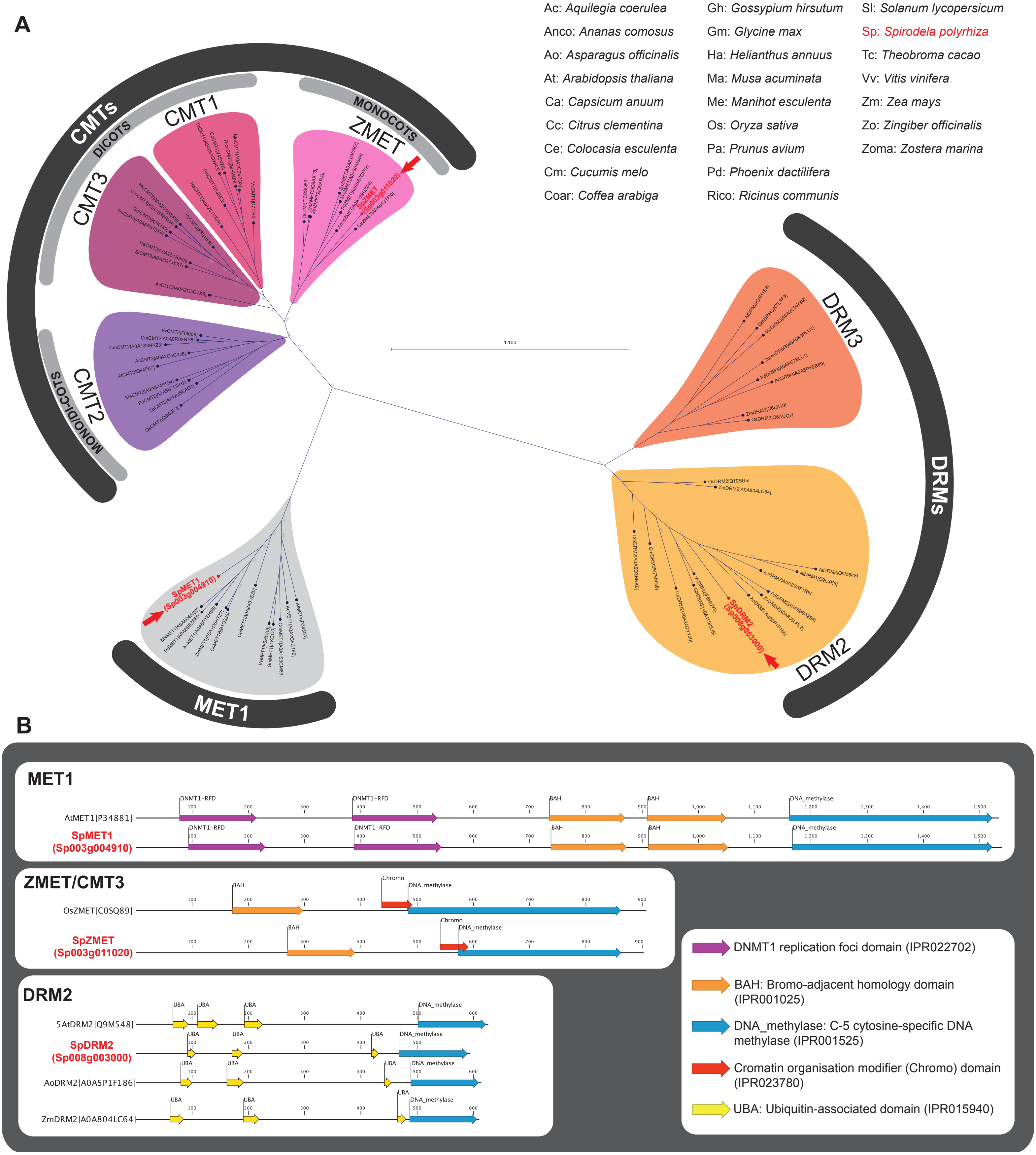

**Supplemental Figure S14: Phylogenetic analysis of *Spirodela polyrhiza* DNA methyltransferases.**
**(A)** Unrooted phylogenetic tree of angiosperm DNA methyltransferase protein families: MET1, Chromomethylases (CMTs) and Domains Rearranged Methyltransferases (DRMs), from several species including those identified in the *Spirodela polyrhiza* genome. The maximum likelihood tree was constructed with 1000 bootstrap replications (see Materials and Methods). Protein families are highlighted in bubbles of different colors. *Spirodela* proteins and annotations were named accordingly to their position in the tree. Uniprot entry codes appear next to the name of proteins in the tree. Arrows point to Spirodela proteins. For *Spirodela*, the gene annotation ID is shown instead. The sequence of all proteins used to build the tree, their alignment and tree files can be found in the raw data files corresponding to this figure. **(B)** Domain annotation of *Spirodela* proteins in (A) along those of their orthologues in other plant species. Numbers along linear protein representations indicate positions in amino acids. Domain annotations were obtained using Interpro and Pfam databases, their integrated InterPro ID codes are display in the graphical legend. Filled arrows indicate significant domain predictions, empty arrows indicate non-significant predictions. For simplicity, Sp9509 gene identifiers have been shortened from Sp9509d(xxx)g(yyyyyy) to Sp(xxx)g(yyyyyy) (where (xxx) is chromosome number, and (yyyyyy) gene number).

### SUPPLEMENTAL FIGURE S15

**A**

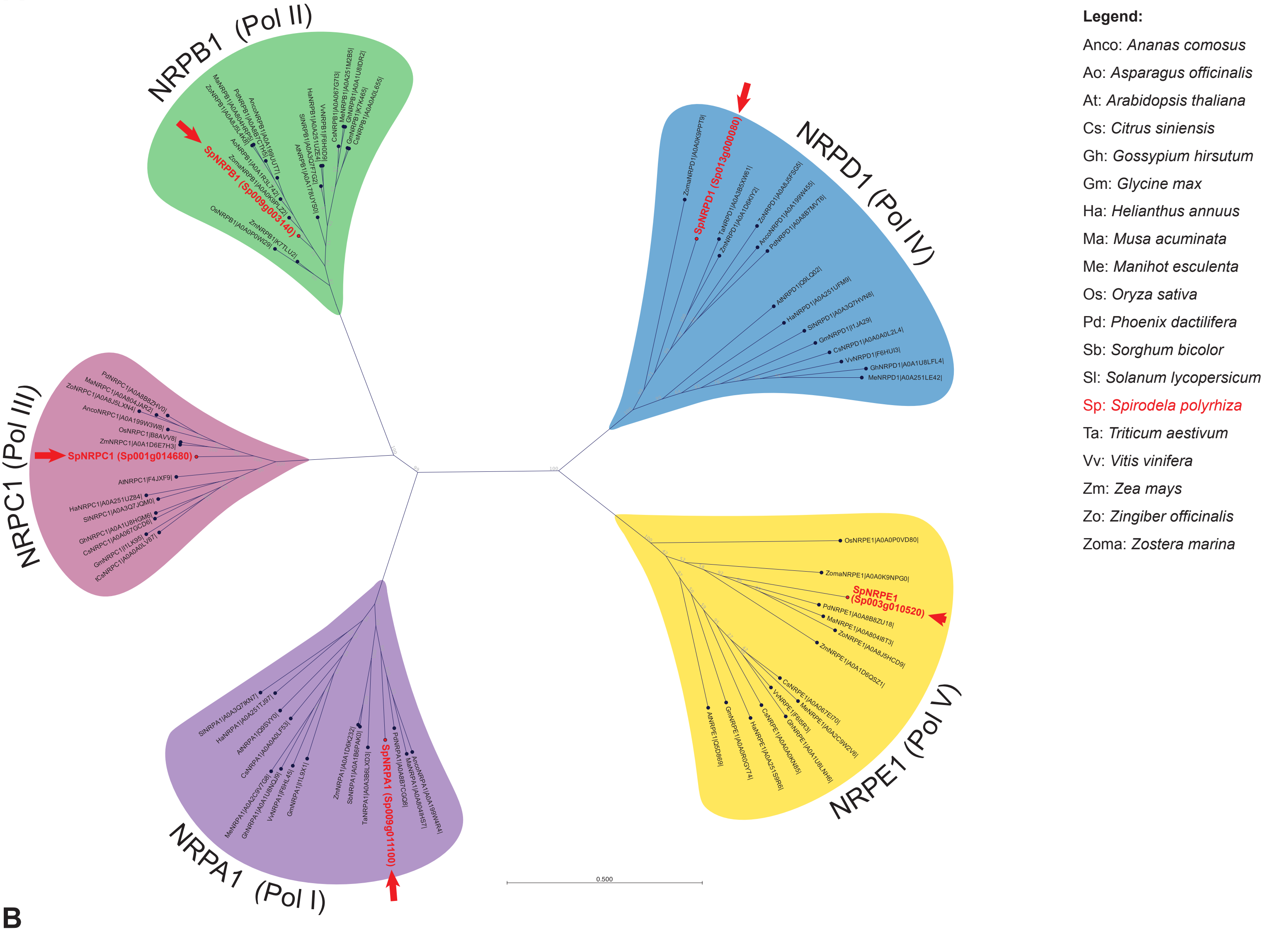

**B**

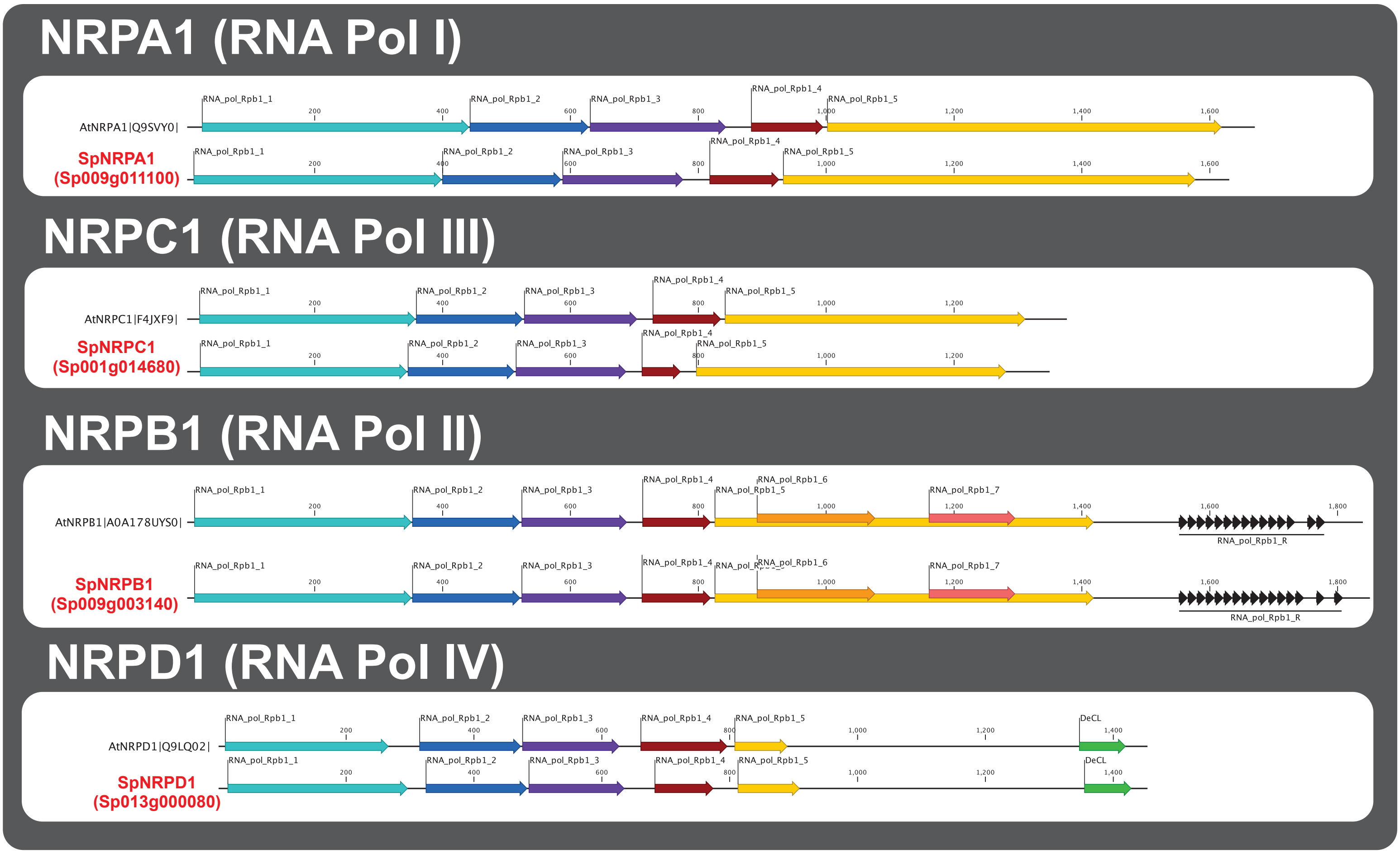

Continues on next page (incl. legend)

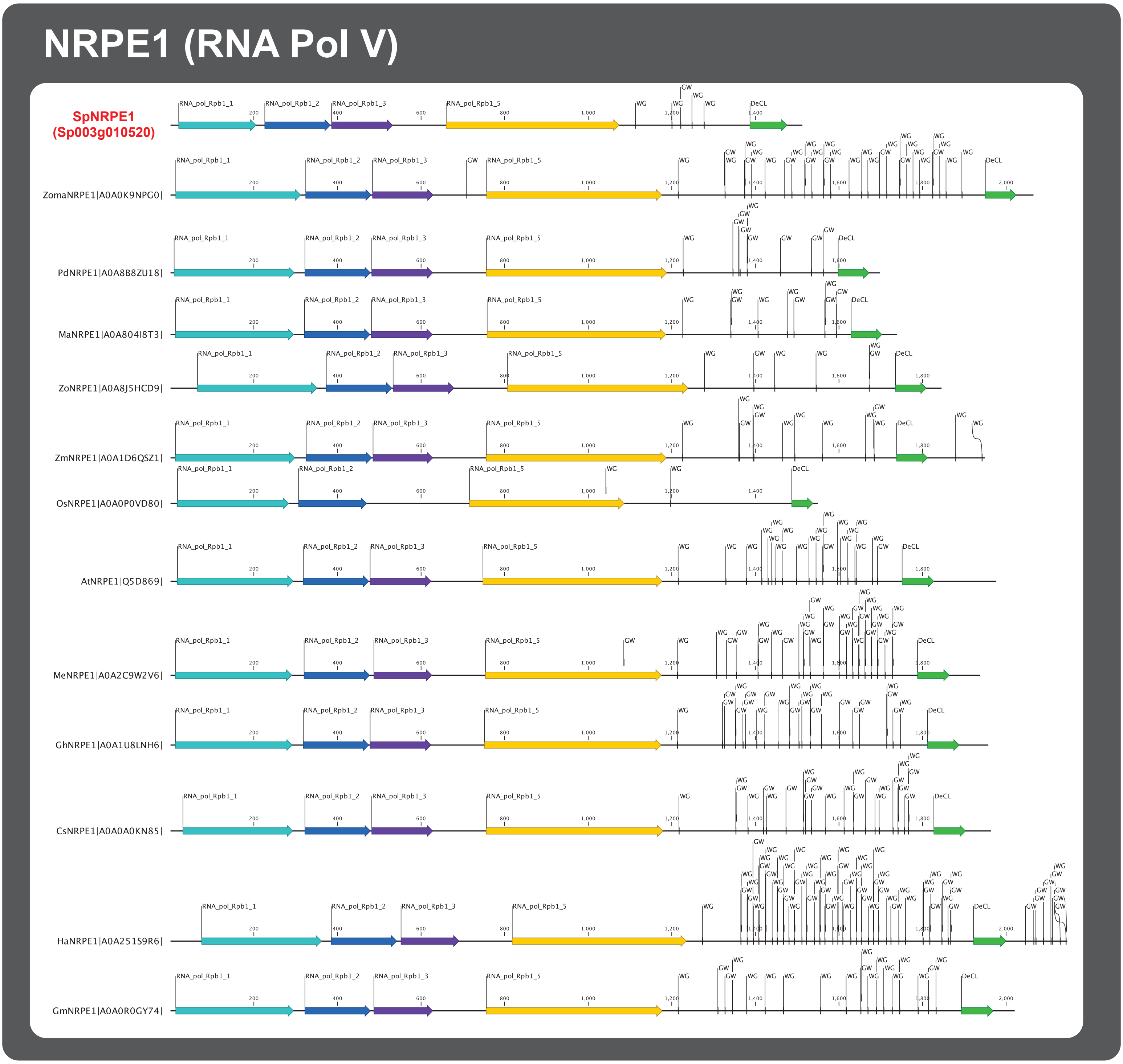

RNA_pol Rpb1_1: DNA-dependent RNA polymerase domain 1. Clamp domain: Maintenance of transcription bubble, DNA and RNA positioning (IPR007080)

RNA_pol Rpb1_2: DNA-dependent RNA polymerase domain 2. Active site (IPR000722)

RNA_pol Rpb1_3: DNA-dependent RNA polymerase domain 3. Pore domain: nucleotides access channel into the active site (IPR007066)

RNA_pol Rpb1_4: DNA-dependent RNA polymerase domain 4. Funnel domain: binding site for elongation factors (IPR007083)

RNA_pol Rpb1_5: DNA-dependent RNA polymerase domain 5. Discontinuous cleft domain: DNA binding (IPR007081)

RNA_pol Rpb1_6: DNA-dependent RNA polymerase domain 6. Shelf module domain (IPR007075)

RNA_pol Rpb1_7: DNA-dependent RNA polymerase domain 7. Part of the lobe domain (IPR007073)

RNA_pol Rpb1_R: Repetitive C-terminal domain (CTD) of RNA Pol II. Regulation of gene expression and Pol II activity dependent of its phosphorylation pattern (IPR000684)

DeCL (DUF3223): RRP6L1 exoribonuclease interacting domain (PF11523 - Not integrated in InterPro)

**GW/WG** repeats: Gly/Trp repeats involved in Argonaute (AGO) binding/recruitment.

**Supplemental Figure S15: Phylogenetic analysis of *Spirodela polyrhiza* RNA Polymerase large subunit (NRP) proteins.** **(A)** Unrooted phylogenetic tree of Nuclear RNA Polymerase (NRP) large subunit proteins from several angiosperms including those identified in the *Spirodela polyrhiza* genome. The maximum likelihood tree was constructed with 1000 bootstrap replications. Protein families are highlighted in bubbles of different colors. Their cognate RNA Polymerase complex (Pol I-V) is indicated in brackets. *Spirodela* proteins and annotations were named accordingly to their position in the tree. Arrows point to Spirodela proteins. Uniprot entry codes appear next to the name of proteins in the tree. For *Spirodela*, the gene annotation ID is shown instead. The sequence of all proteins used to build the tree, their alignment and tree files can be found in the raw data files corresponding to this figure. **(B)** Domain annotation of *Spirodela* proteins in (A) along those of their orthologues in other plant species. Numbers along linear protein representations indicate positions in amino acids. Domain annotations were obtained using InterPro and Pfam databases, their integrated InterPro ID codes are display in the graphical legend. Filled arrows indicate significant domain predictions, empty arrows indicate non-significant predictions. Position of Gly/Trp (GW/WG) repeats (AGO-hooks) for NRPE1 are also indicated. For simplicity, Sp9509 gene identifiers have been shortened from Sp9509d(xxx)g(yyyyyy) to Sp(xxx)g(yyyyyy) (where (xxx) is chromosome number, and (yyyyyy) gene number).

### SUPPLEMENTAL FIGURE S16

**A**

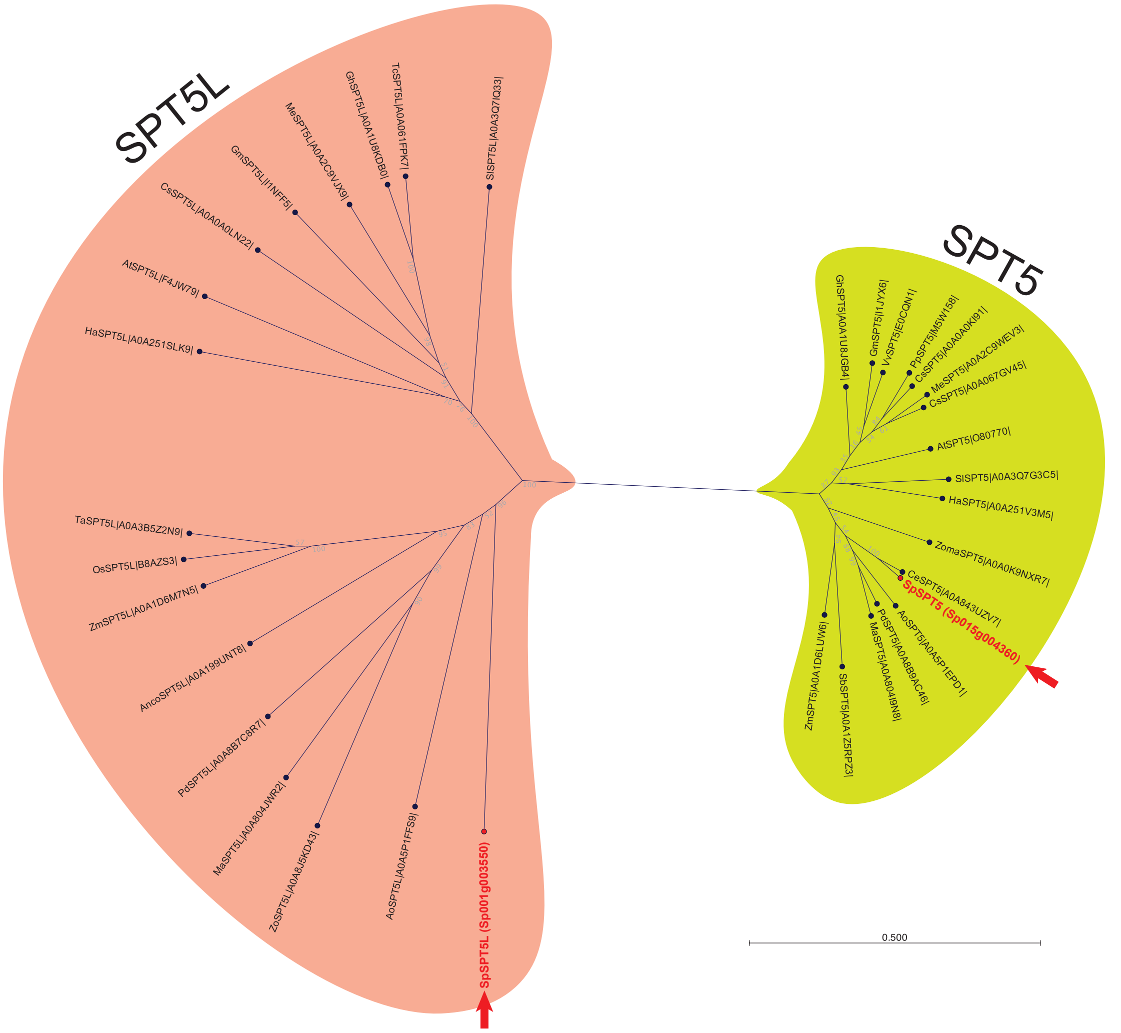

**Legend:**

Anco: *Ananas comosus*
At: *Arabidopsis thaliana*
Ao: *Asparagus officinalis*
Ce: *Colocasia esculenta*
Cs: *Citrus siniensis*
Gh: *Gossypium hirsutum*
Gm: *Glycine max*
Ha: *Helianthus annuus*
Ma: *Musa acuminata*
Me: *Manihot esculenta*
Os: *Oryza sativa*
Pd: *Phoenix dactilifera*
Sb: *Sorghum bicolor*
Sl: *Solanum lycopersicum*
Sp: *Spirodela polyrhiza*
Ta: *Triticum aestivum*
Tc: *Theobroma cacao*
Vv: *Vitis vinifera*
Zm: *Zea mays*
Zo: *Zingiber officinalis*
Zoma: *Zostera marina*

**B**

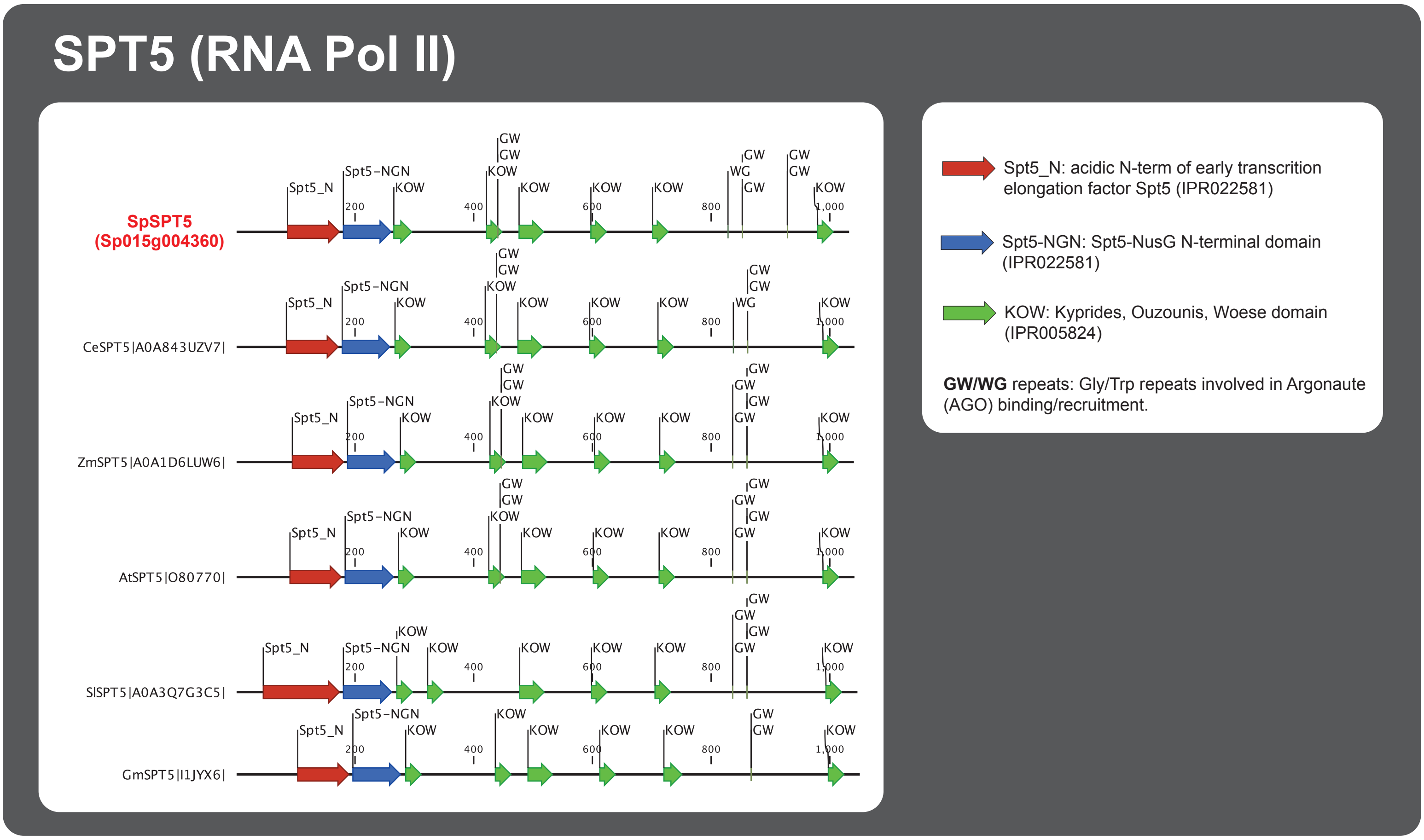

Continues on next page (incl. legend)

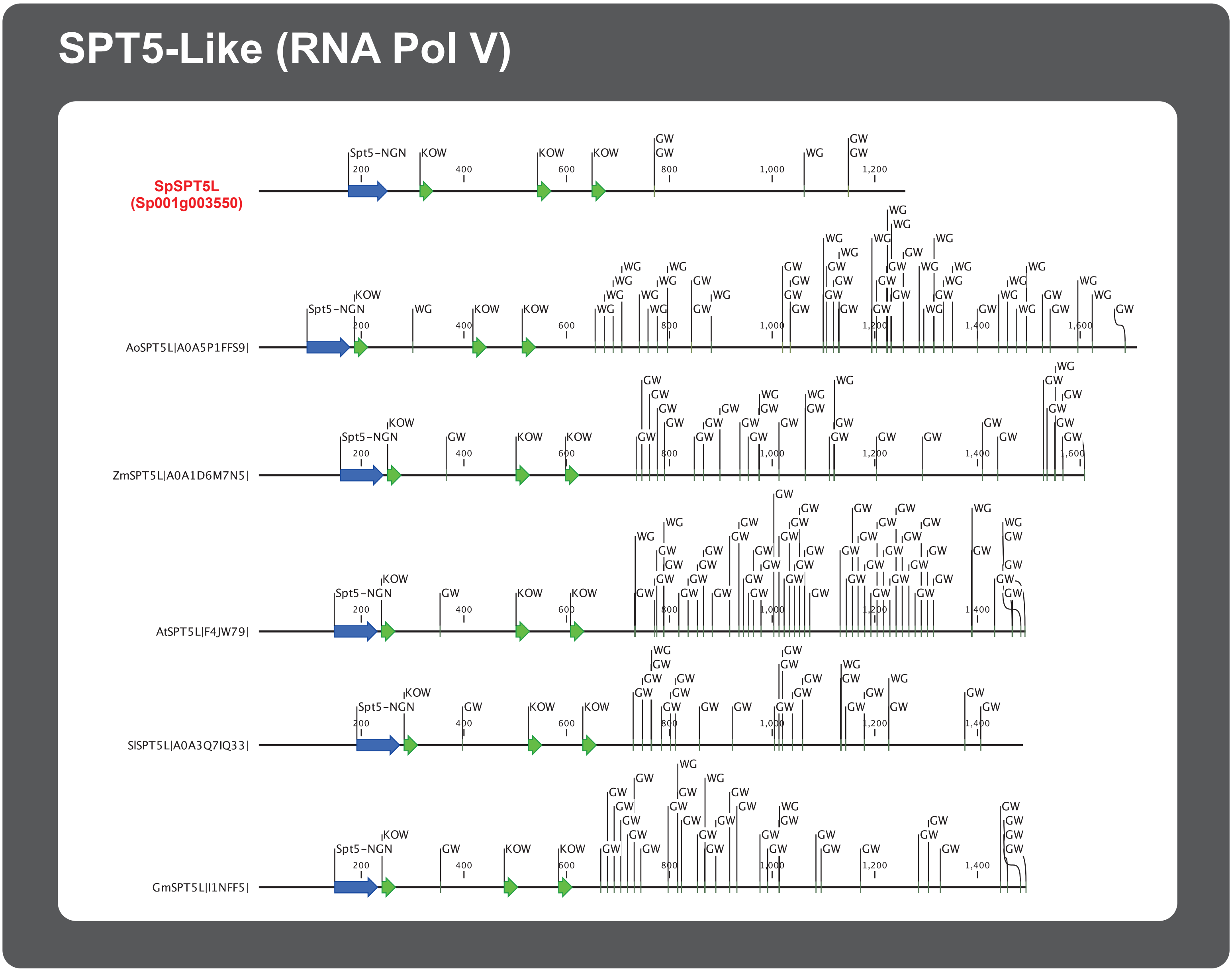

**Supplemental Figure S16: Phylogenetic analysis of *Spirodela polyrhiza* SPT5 and SPT5L proteins.**
**(A)** Unrooted phylogenetic tree of the Pol II transcription elongation factor SPT5 and it's paralogue, the Pol V interactor, SPT5-like (SPT5L) proteins from several angiosperms including those identified in the *Spirodela polyrhiza* genome. The maximum likelihood tree was constructed with 1000 bootstrap replications. *Spirodela* proteins and annotations were named accordingly to their position in the tree. Arrows point to *Spirodela* proteins. Uniprot entry codes appear next to the name of proteins in the tree. For *Spirodela*, the gene annotation ID is shown instead. The sequence of all proteins used to build the tree, their alignment and tree files can be found in the raw data files corresponding to this figure. **(B)** Domain annotation of *Spirodela* proteins in (A) along those of their orthologues in other plant species. Numbers along linear protein representations indicate positions in amino acids. Domain annotations were obtained using InterPro and Pfam databases, their integrated InterPro ID codes are display in the graphical legend. Filled arrows indicate significant domain predictions, empty arrows indicate non-significant predictions. Position of Gly/Trp (GW/WG) repeats (AGO-hooks) are also indicated. For simplicity, Sp9509 gene identifiers have been shortened from Sp9509d (xxx)g(yyyyyy) to Sp(xxx)g(yyyyyy) (where (xxx) is chromosome number, and (yyyyyy) gene number).

### SUPPLEMENTAL FIGURE S17

A

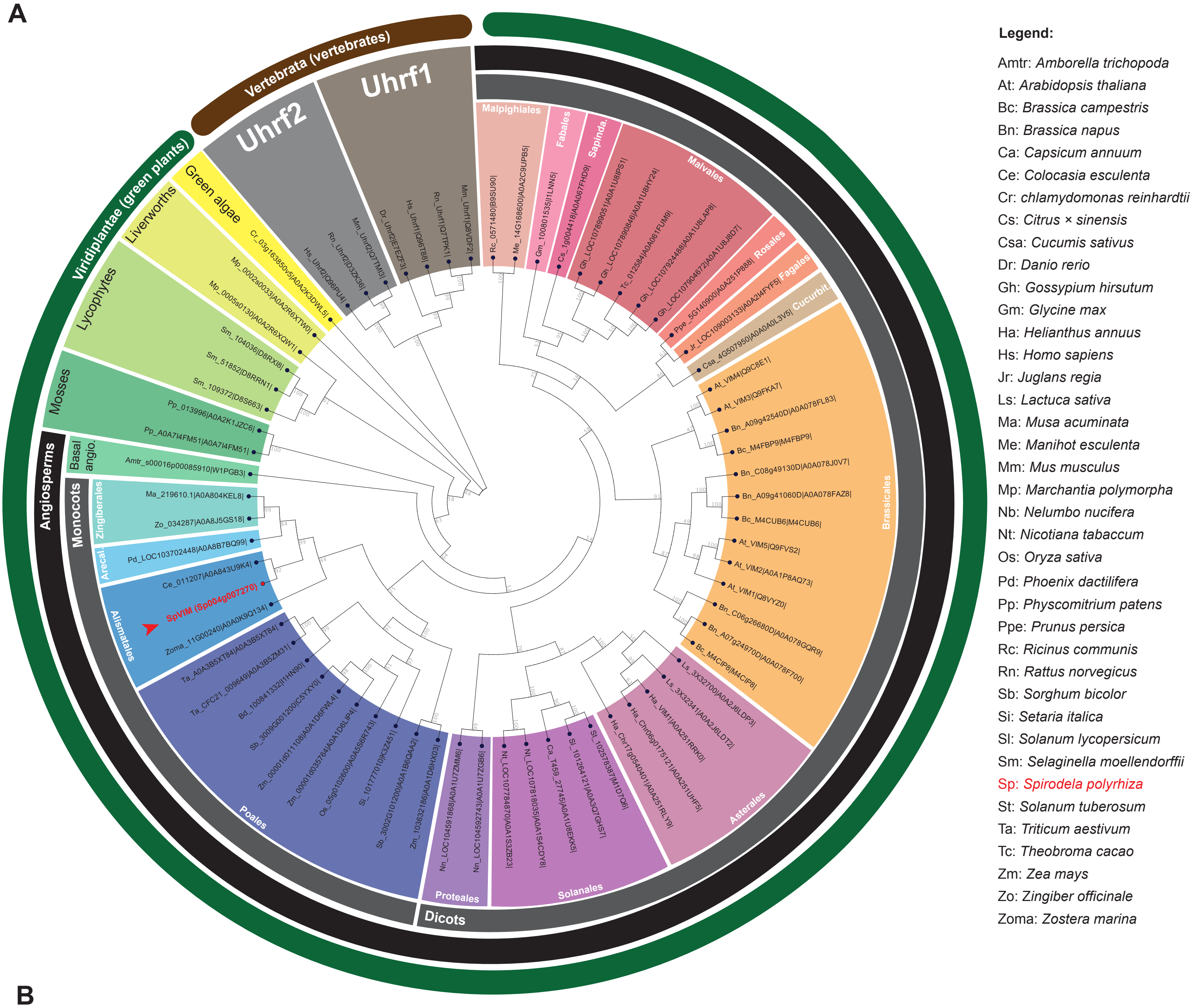

B

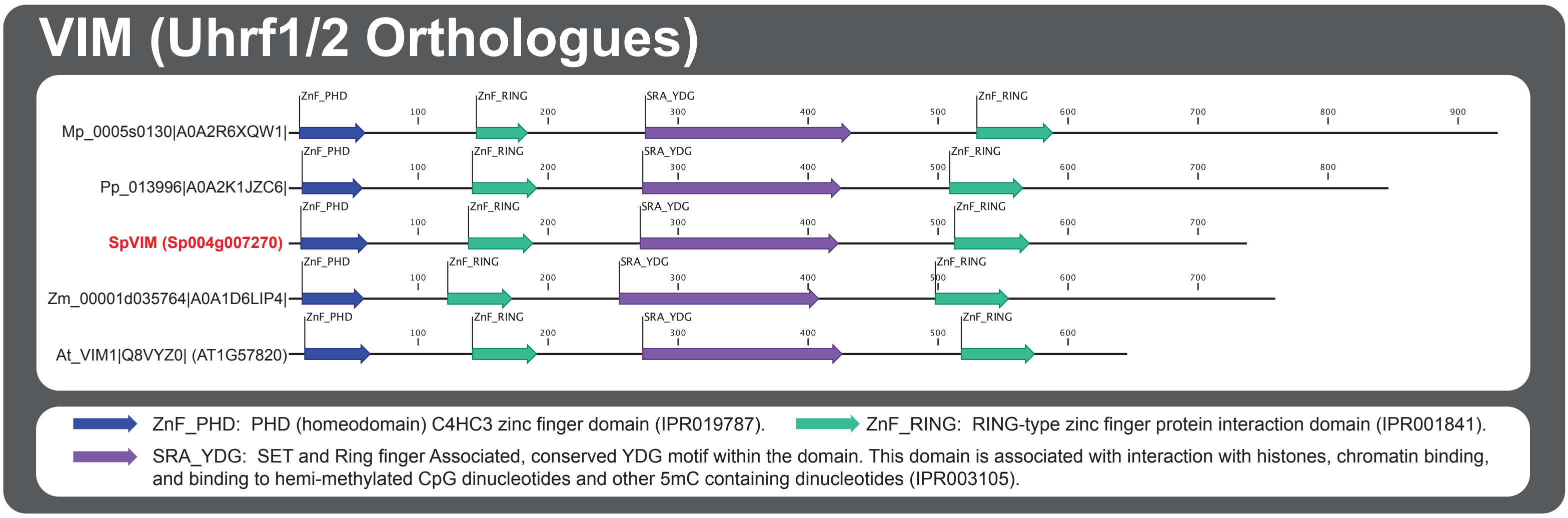

**Supplemental Figure S17: Phylogenetic analysis of *Spirodela polyrhiza* VIM proteins.**
**(A)** Unrooted phylogenetic tree of animal and plant Uhrf1/2 orthologues, VIM, from several species including that identified in the *Spirodela polyrhiza* genome. The maximum likelihood tree was constructed with 1000 bootstrap replications. Gene ID ad Uniprot entry codes appear next to the name of proteins in the tree. For *Spirodela*, the gene annotation ID is shown instead. Arrows point to *Spirodela* proteins. The sequence of all proteins used to build the tree, their alignment and tree files can be found in the raw data files corresponding to this figure. (**B)** Domain annotation of *Spirodela* proteins in (A) along those of their orthologues in other species. Numbers along linear protein representations indicate positions in amino acids. Domain annotations were obtained using the InterPro database, their ID codes are display in the graphical legend. For simplicity, Sp9509 gene identifiers have been shortened from Sp9509d(xxx)g(yyyyyy) to Sp(xxx)g(yyyyyy) (where (xxx) is chromosome number, and (yyyyyy) gene number).

### SUPPLEMENTAL FIGURE S18

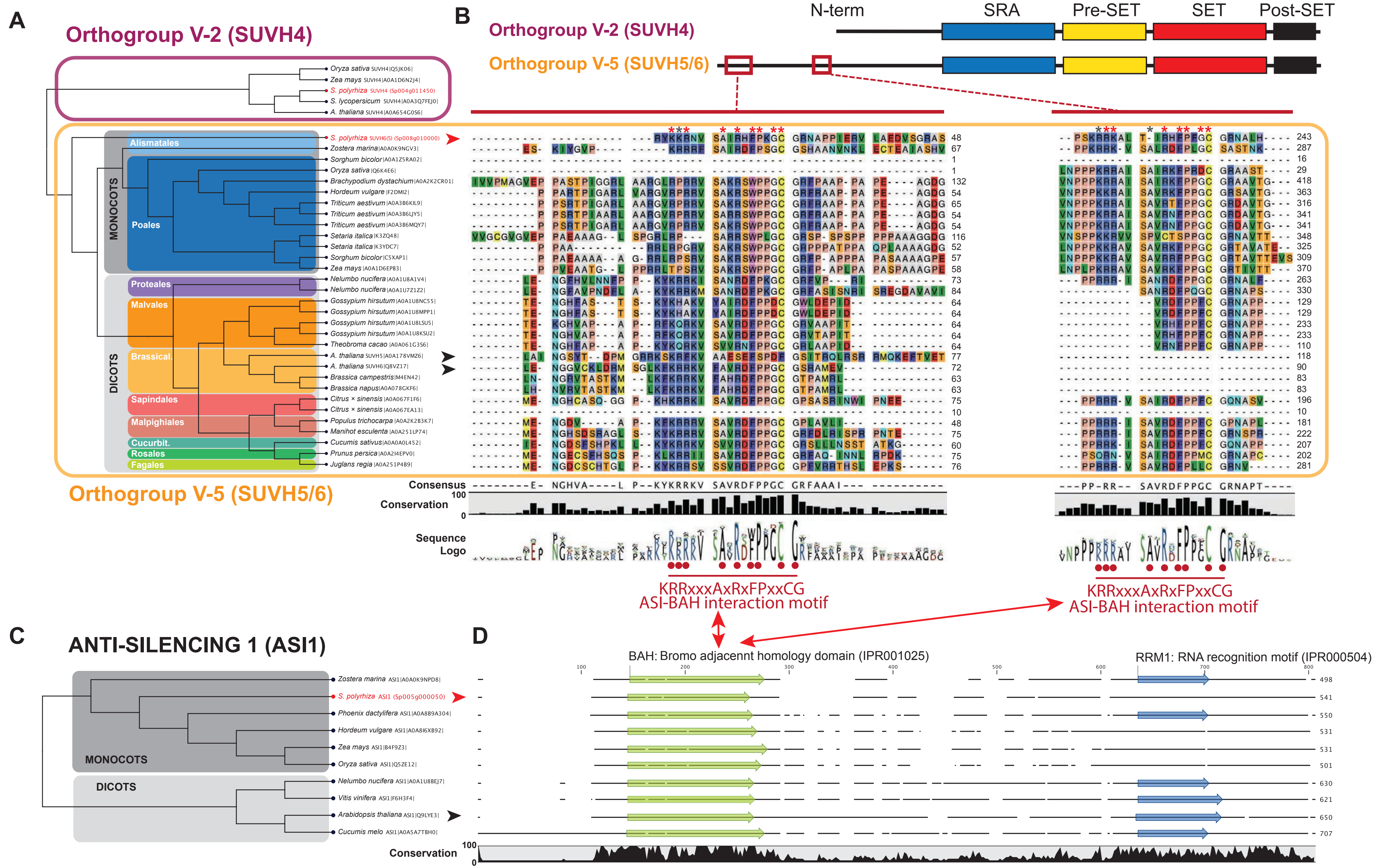

**Supplemental Figure S18: Phylogenetic analysis of *Spirodela polyrhiza* SET-domain Class V orthogroup V-5 (SUVH5/6) and ASI1 proteins.**
**(A)** Unrooted phylogenetic tree of angiosperm Class V SET Orthogroup V-5 proteins (SUVH5/6) from several species including those identified in the *Spirodela polyrhiza* genome. SUVH4 (orthogroup V2) proteins were added as outgroup. The maximum likelihood tree was constructed with 1000 bootstrap replications. Arrows point to *Spirodela* (red) and *Arabidopsis* (black) proteins. Uniprot entry codes appear next to the name of proteins in the tree. For *Spirodela*, the gene annotation ID is shown instead. **(B)** Shchematic representation of SUVH4/5/6 domain structure along their protein sequence and multiple protein sequence alignment of SUVH5/6 proteins from A. Only N-terminal regions from SUVH5/6 containing the conserved ASI-BAH interaction motif are shown along their consensus sequence, sequence logo and conservation. The ASI-BAH interaction motif, as described in Zhang *et al.,* 2023, is indicated below. Red dots mark the conserved aa's identified in Zhang *et al.,* 2023. Asteriks above *Spirodela* SUVH6(5) mark, in red, conserved or, in grey, non-conserved or missing aa from the motif. **(C)** Same as A but for ASI1 orthologues. **(D)** Multiple protein sequence alignment of ASI1 proteins from C along their conservation. Domain annotations were obtained using Interpro and Pfam databases, their integrated InterPro ID codes are displayed. For simplicity, Sp9509 gene identifiers have been shortened from Sp9509d(xxx)g(yyyyyy) to Sp(xxx)g(yyyyyy) (where (xxx) is chromosome number, and (yyyyyy) gene number). The sequence of all proteins used to build the tree, their alignment and tree files can be found in the raw data files corresponding to this figure.

### SUPPLEMENTAL FIGURE S19

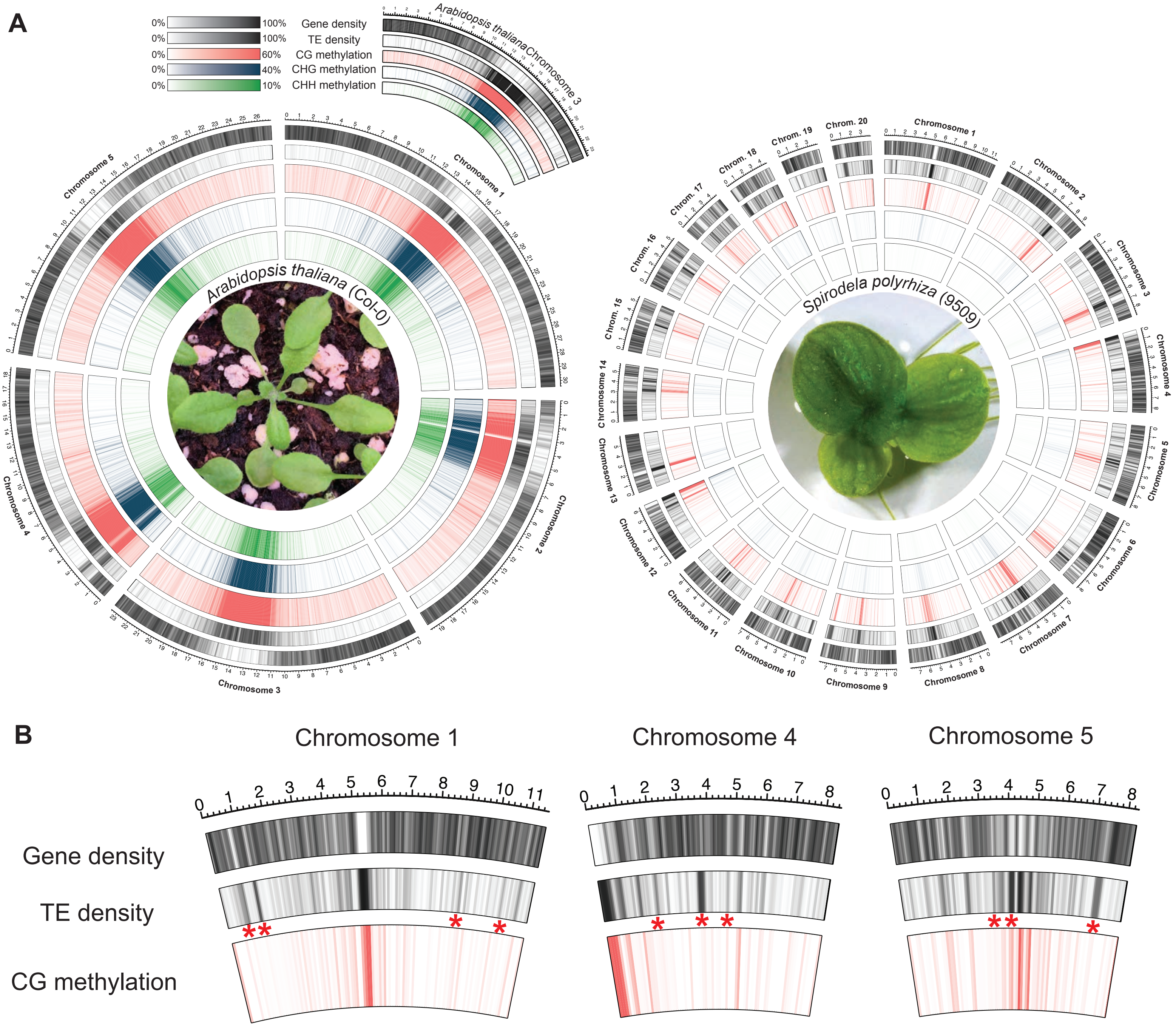

**Supplemental Figure S19: Genome-wide distribution of DNA methylation in *Spirodela polyrhiza*.**
**A** Circos plot of genome-wide distribution of DNA methylation densities by chromosome in *Spirodela* and *Arabidopsis*. From outside to inside: gene density, TE density, CG, CHG and CHH methylation. Mean DNA methylation, gene and TE densities values within the indicated ranges are displayed for 100 kbp intervals in 50 kbp sliding windows. **B** Detail of three *Spirodela* chromosomes from (A) showing gene, TEs and CG methylation densities. Asterisks mark high TE density regions with low CG methylation levels.

### SUPPLEMENTAL FIGURE S20

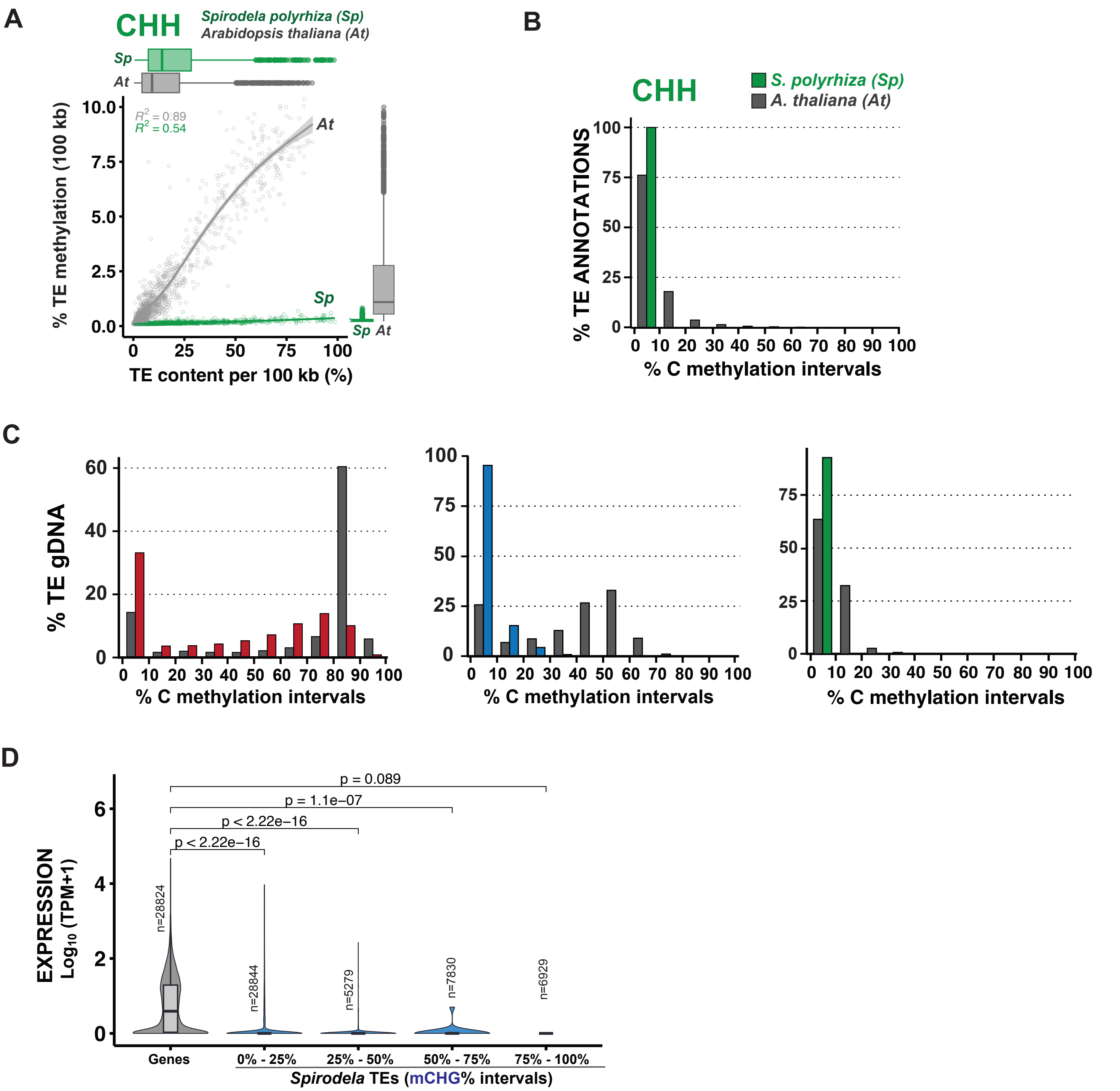

**Supplemental Figure S20: Distribution of CHH DNA methylation in *Spirodela polyrhiza*.**

**(A)** Scatter plot and best fit line of average TE CHH DNA methylation against TE content per 100 kb genomic bins in *Arabidopsis* and *Spirodela*. **(B)** Bar plot of the percentage of TE-occupied genomic space displaying a given DNA methylation level within 10% incremental intervals of CG, CHG and CHH methylation in *Arabidopsis* and *Spirodela*. **(C)** Bar plot of the distribution (%) of TE annotations (>=100 bp) within 10% incremental intervals of CHH methylation in *Arabidopsis* and *Spirodela*. **(D)** Analysis of *Spirodela* TE expression within 25% incremental intervals of CHG methylation. Global gene expression values are shown for comparison. Wilcoxon rank sum test was used to compare expression level between genes and each TE intervals. In all boxplots: median is indicated by solid bar; the boxes extend from the first to the third quartile and whiskers stretch to the furthest value within 1.5 times the interquartile range.

### SUPPLEMENTAL FIGURE S21

| CLASS | ORDER | SUPERFAMILY | CODE |
| --- | --- | --- | --- |
| *Class I: Retrotransposons* (”copy-paste”) **(R)** | Long Terminal Repeat (LTR) **(L)** | *Ty1/Copia* **(C)** | **RLC** |
| | | *Ty3/Gypsy* **(G)** | **RLG** |
| | | *Unknown* **(X)** | **RLX** |
| | non-LTR LINE **(I)** | *LINE* **(L)** | **RIL** |
| | | *Other/Unknown* **(X)** | **RIX** |
| | non-LTR SINE **(S)** | *tRNA* **(T)** | **RST** |
| | | *7SL* **(L)** | **RSL** |
| | | *5S* **(S)** | **RSS** |
| | | *Unknown* **(X)** | **RSX** |
| | *Unknown* **(X)** | *Unknown* **(X)** | **RXX** |
| *Class II: DNA transposons* (cut-paste) **(D)** | Terminal Inverted Repeat (TIR) **(T)** | *Tc1/Mariner* **(T)** | **DTT** |
| | | *Mutator* **(M)** | **DTM** |
| | | *PIF-Harbinger* **(H)** | **DTH** |
| | | *CACTA* **(C)** | **DTC** |
| | | *hAT* **(A)** | **DTA** |
| | | *Unknown* **(X)** | **DTX** |
| | non-TIR **(H)** | *Helitron* **(H)** | **DHH** |
| | | *Unknown* **(X)** | **DHX** |
| | *Unknown* **(X)** | *Unknown* **(X)** | **DXX** |
| *Unknown* **(X)** | *Unknown* **(X)** | *Unknown* **(X)** | **XXX** |

**Supplemental Figure S21: Transposon classification, nomenclature and codes used in this study.**

### SUPPLEMENTAL FIGURE S22

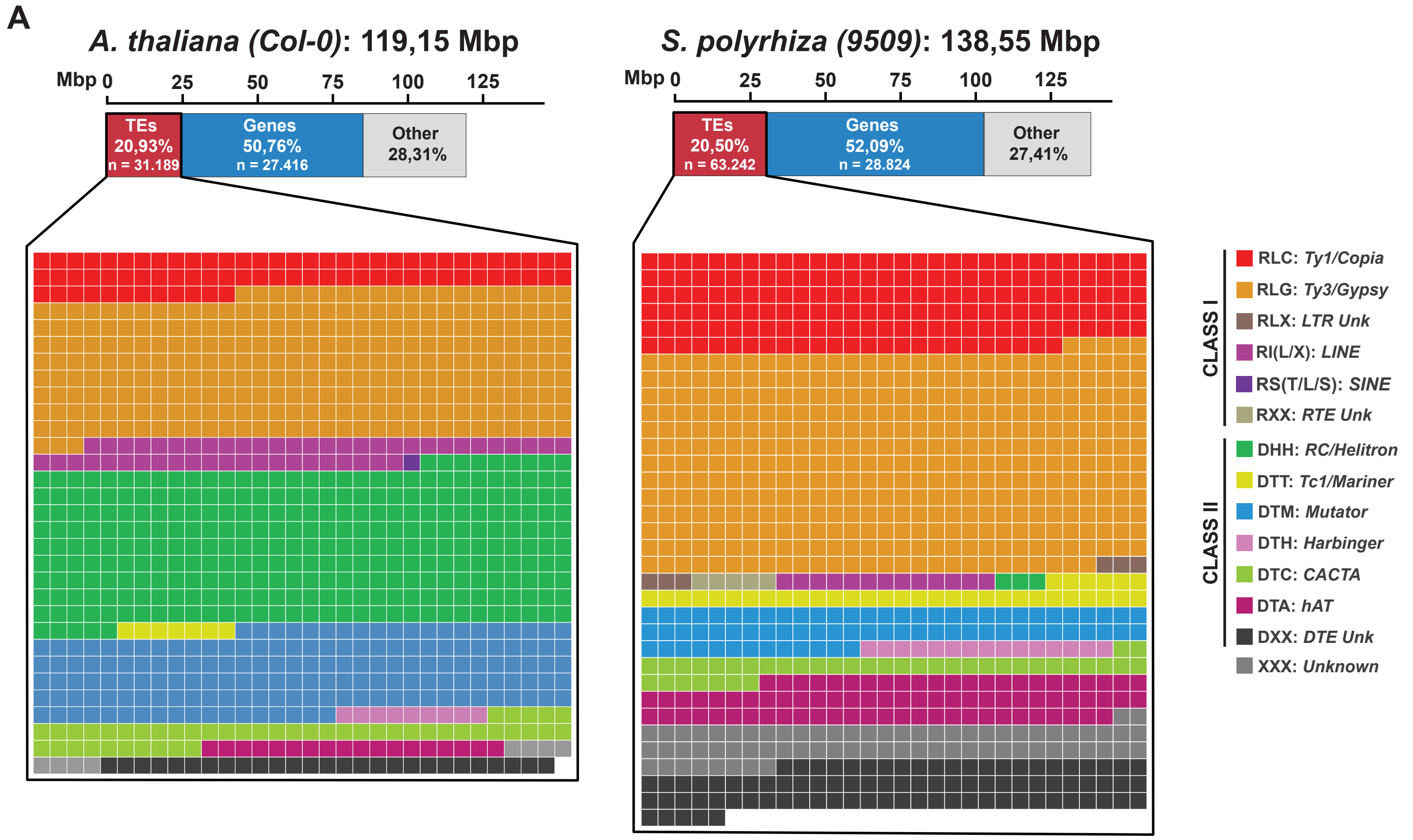

**B**

| | Superfamily | *Arabidopsis thaliana* (Col-0) DNA (kbp) | % gDNA | % TE gDNA | *Spirodela polyrhiza* (9509) DNA (kbp) | % gDNA | % TE gDNA |
|---|---|---|---|---|---|---|---|
| CLASS I | RLC | 1920,719 | 1,612 % | 7,694 % | 5415,233 | 3,908 % | 17,512 % |
| | RLG | 6978,646 | 5,857 % | 27,956 % | 12123,494 | 8,750 % | 39,206 % |
| | RLX | - | - | - | 200,412 | 0,145 % | 0,648 % |
| | RI(L/X) | 1276,831 | 1,072 % | 5,115 % | 411,652 | 0,297 % | 1,331 % |
| | RS(T/L/X) | 32,105 | 0,027 % | 0,129 % | 23,336 | 0,017 % | 0,075 % |
| | RXX | - | - | - | 155,536 | 0,112 % | 0,503 % |
| CLASS II | DHH | 7560,501 | 6,346 % | 30,287 % | 103,235 | 0,075 % | 0,334 % |
| | DTT | 190,03 | 0,159 % | 0,761 % | 1140,593 | 0,823 % | 3,689 % |
| | DTM | 4148,023 | 3,481 % | 1,662 % | 2259,622 | 1,631 % | 7,307 % |
| | DTH | 248,765 | 0,209 % | 0,997 % | 478,383 | 0,345 % | 1,547 % |
| | DTC | 1178,414 | 0,989 % | 4,721 % | 1214,221 | 0,876 % | 3,927 % |
| | DTA | 462,131 | 0,388 % | 1,851 % | 2521,626 | 1,820 % | 8,155 % |
| | DXX | 689,039 | 0,578 % | 2,760 % | 2691,772 | 1,943 % | 8,705 % |
| Unk | XXX | 55,341 | 0,046 % | 0,222 % | 2183,405 | 1,576 % | 7,061 % |

**Supplemental Figure S22: Transposon composition of *Arabidopsis thaliana* (Col-0) and *Spirodela polyrhiza* (9509) genomes**

**(A)** Waffle chart of the relative contribution of different TE superfamilies over the total genomic space occupied by TEs in *Arabidopsis* and *Spirodela.* **(B)** Table with TE superfamily content as total amount of DNA (in kbp), relative to genome size (% gDNA) and the portion of the genome occupied by TEs (%TE gDNA) in both species.

### SUPPLEMENTAL FIGURE S23

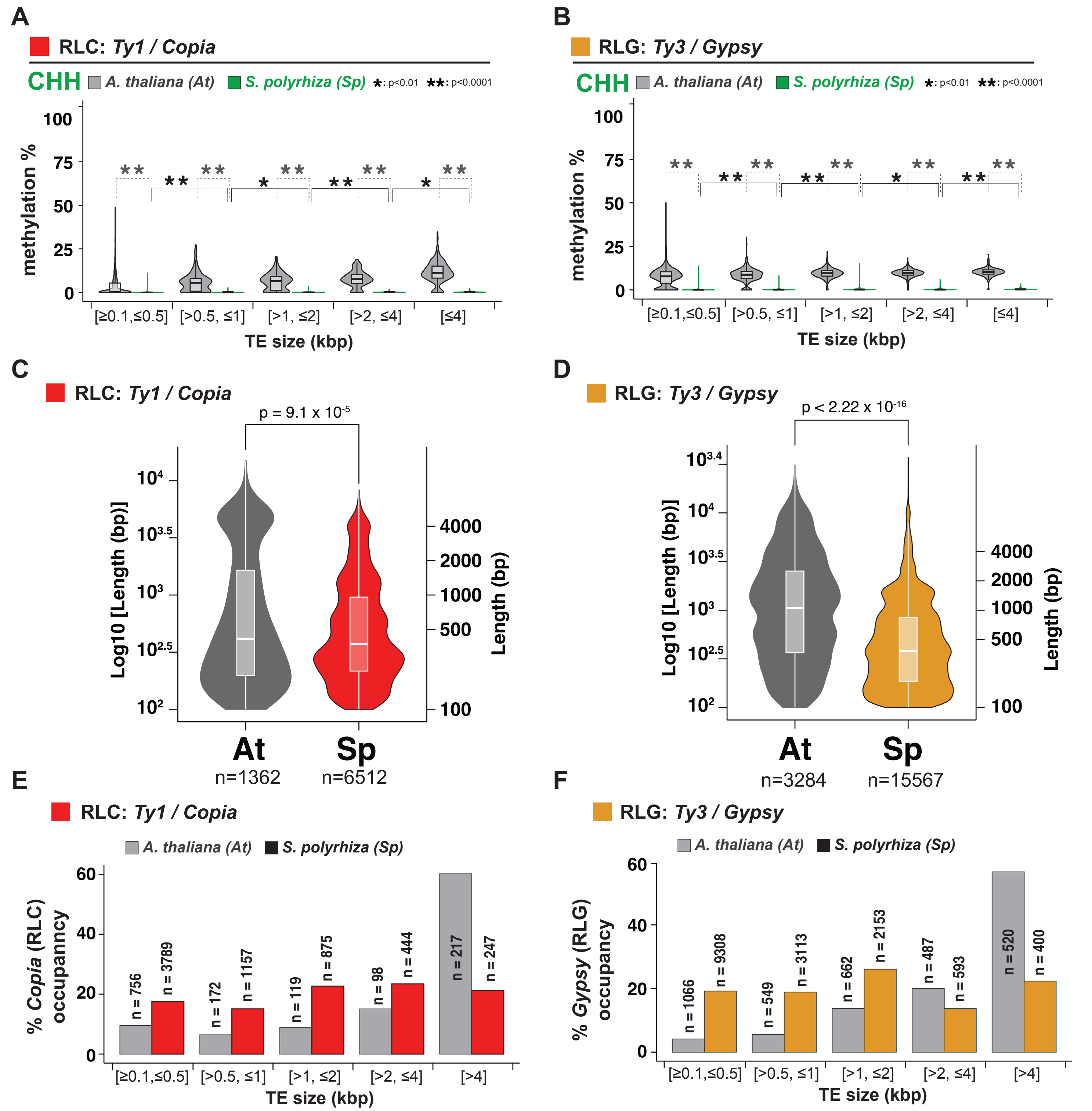

**Supplemental Figure S23: mCHH levels across TE sizes and RLC and RLG length distribution in *Arabidopsis* and *Spirodela* genomes.**

**(A-B)** Analysis of CHH methylation levels (%) between different length range groups for RLC: *Ty1/Copia* (a) and RLG: *Ty3/Gypsy* (b) in *Arabidopsis* and *Spirodela*. n values are indicated in panels E and F **(C-D)** Length distribution of RLC (C) and RLG (D) TE annotations in Arabidopsis and Spirodela genomes. **(E-F)** Number of TE annotations and relative contribution to their superfamily genomic occupancy at each size bin for RLC (E) and RLG (F). (*) indicates p-value <0.01 (**) indicates p-value <0.0001 (Wilcoxon rank sum test) In all boxplots: median is indicated by solid bar; the boxes extend from the first to the third quartile and whiskers stretch to the furthest value within 1.5 times the interquartile range. At: *Arabidopsis*, Sp: *Spirodela*.

### SUPPLEMENTAL FIGURE S24

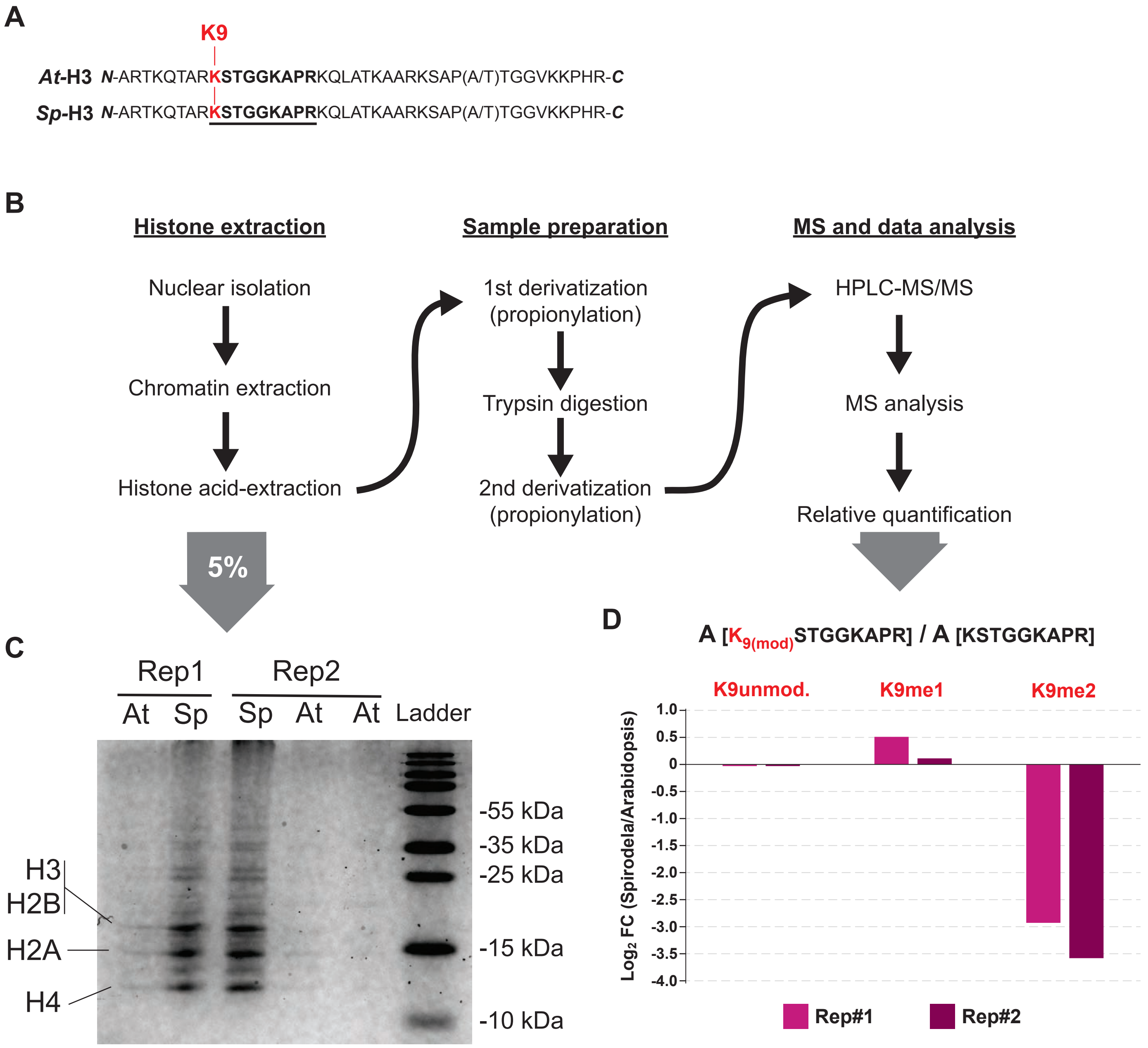

**Supplemental Figure S24: Mass spectrometry (MS) quantification of H3K9me1/2 abundance in *Spirodela*.**
**(A)** Sequence of the 40 first aa of *Spirodela* and *Arabidopsis* Histone 3 N-term. Lysine 9 (K9) and the associated peptide in the MS data is indicated and underlined. **(B)** Simplified workflow scheme of the steps from sample preparation to MS used. **(C)** Coomassie staining of denaturing SDS-PAGE gel of histones after acid extraction for quantity and quality control prior to sample preparation for MS. **(D)** Relative abundance of unmodified H3K9, H3K9me1 and H3K9me2 in *Spirodela* compared to *Arabidopsis*. Abundance of each modification was calculated in each species by the ratio between the area of each modified peptide to the sum area of the peptide containing H3K9 independently of the modifications present.

### SUPPLEMENTAL FIGURE S25

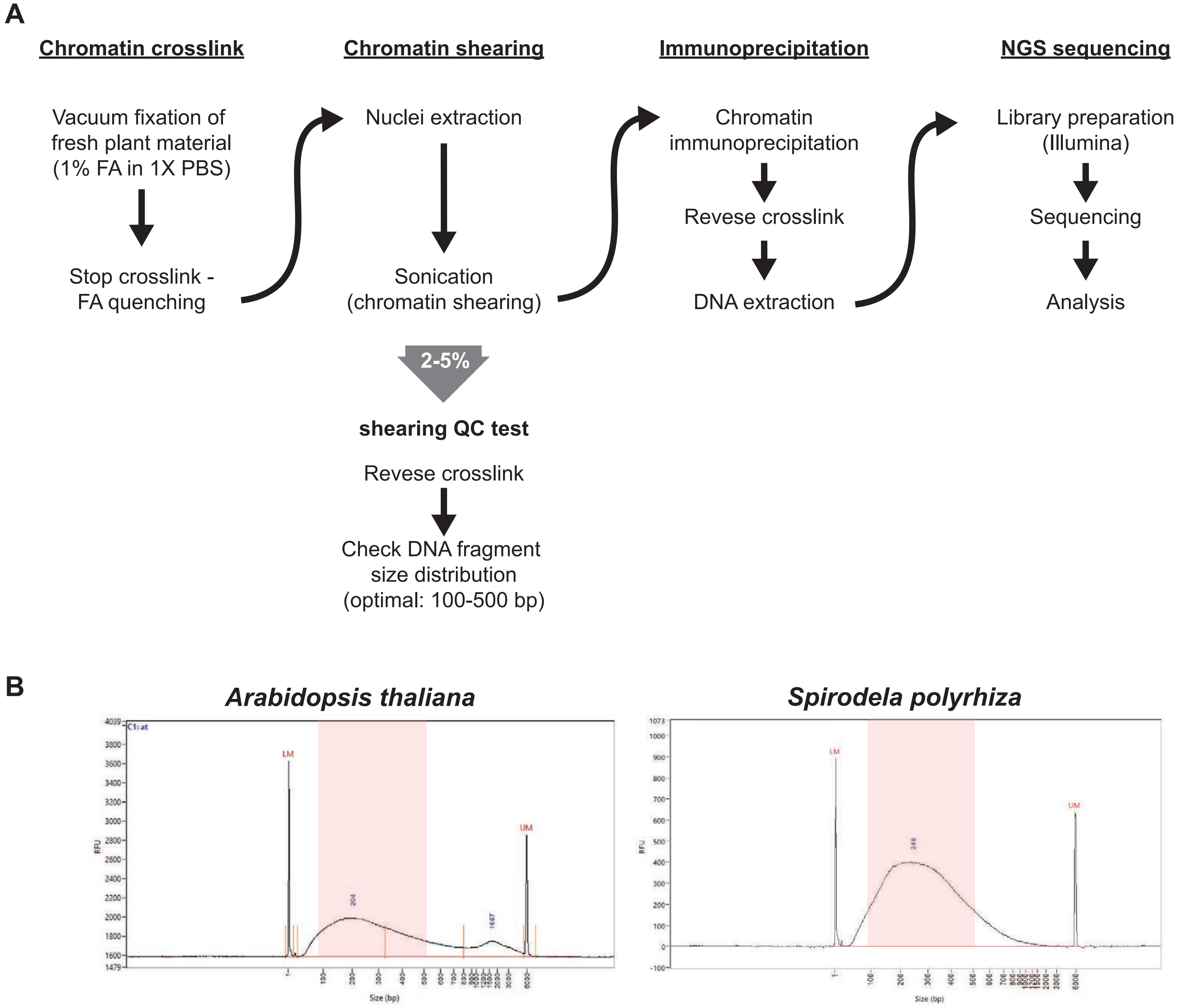

**Supplemental Figure S25: Chromatin immunoprecipitation (ChIP) workflow and quality control in *Spirodela*.**
**(A)** Simplified workflow scheme of the ChIP and control steps. **(B)** Size distribution of DNA fragments after chromatin shearing and sample preparation quality control prior to IP (by innvestigating DNA fragment legth distribution in a fragment analyser). The 100 to 500 bp size range is shadowed. LM and UM, lower and upper marker from Agilent Fragment Analyser respectively.

### SUPPLEMENTAL FIGURE S26

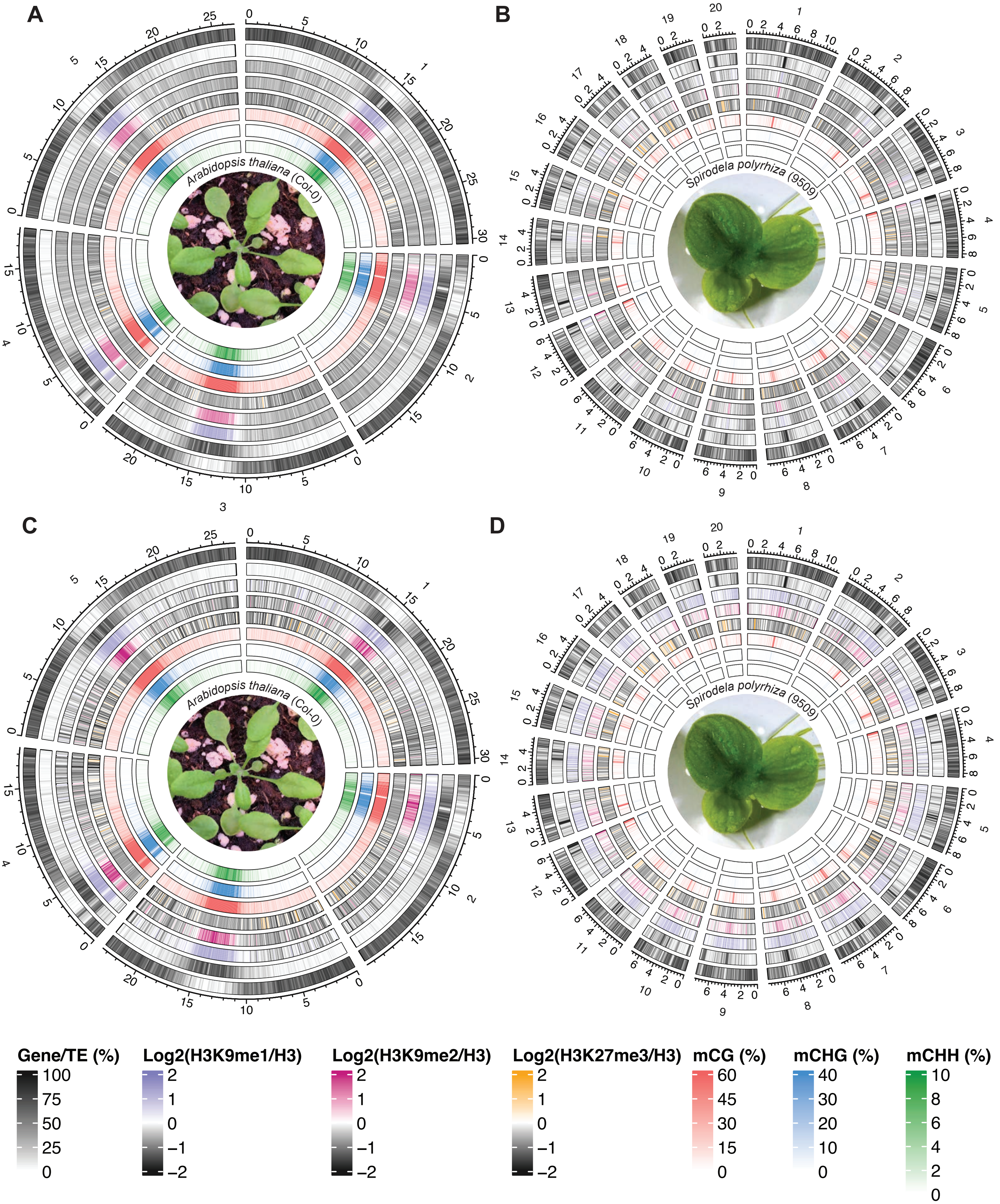

**Supplemental Figure S26: Genome-wide distribution of H3K9me1, H3K9me2 and H3K27me3 in *Spirodela polyrhiza*.**
**(A-B)** Circos plot of genome-wide distribution of H3K9me1, H3K9me2 and H3K27me3 enrichment densities by chromosome in *Spirodela* and *Arabidopsis*. From outside to inside: gene density, TE density, H3K9me1, H3K9me2, H3K27me3, CG, CHG and CHH methylation. Mean values within the indicated ranges are displayed for 100 kbp intervals in 50 kbp sliding windows. **(C-D)** Same as (A-B) but mean values were calculated only considering TE annotations (>=100 bp) for 100 kbp intervals in 50 kbp sliding windows.

### SUPPLEMENTAL FIGURE S27

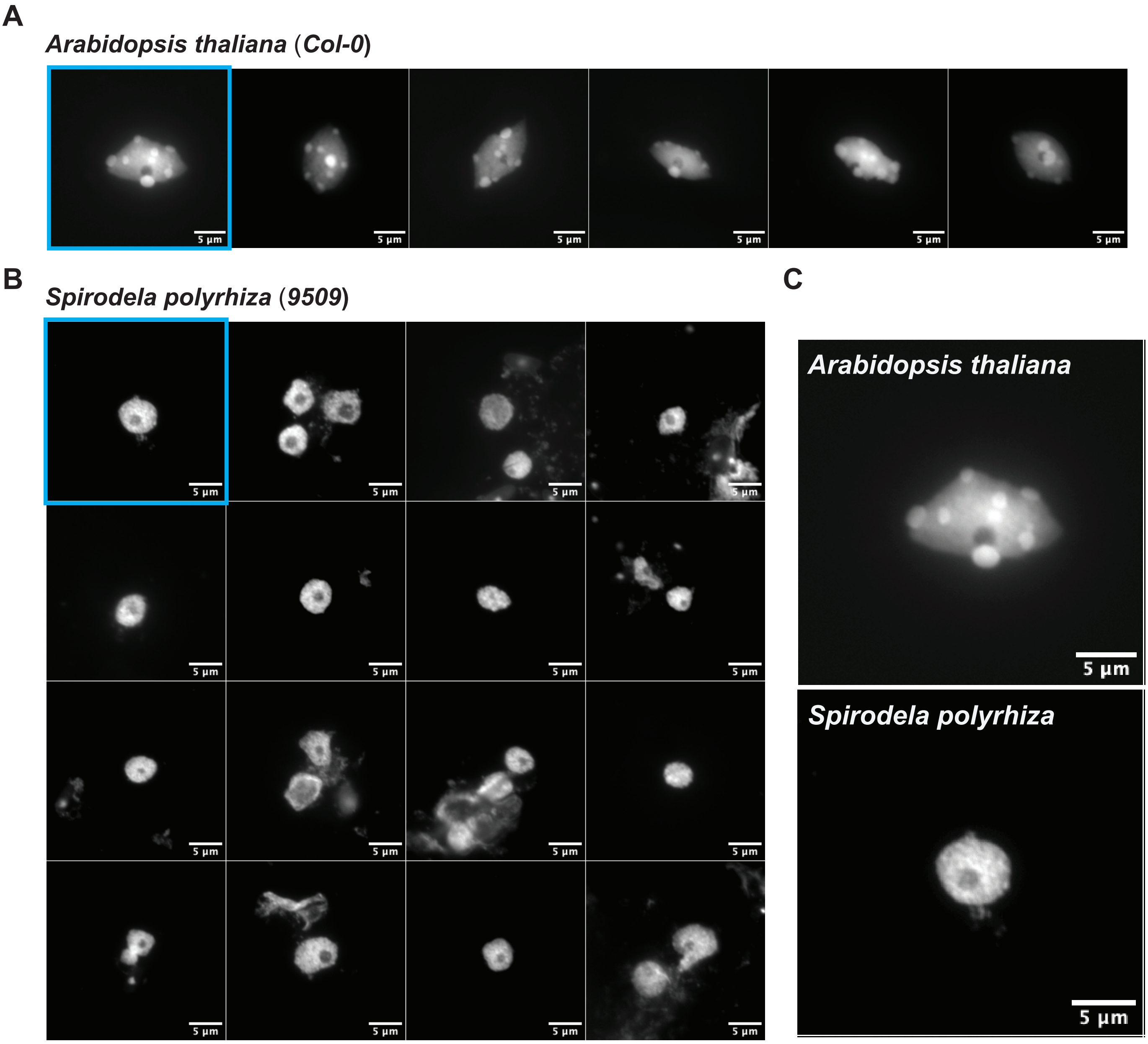

**Supplemental Figure S27: *Spirodela* chromatin organization in interphase nuclei.**
**(A-B)** Gray-scale fluorescent images of DAPI-stained interphase nuclei of *Arabidopsis* (A) and *Spirodela* (B). **C** Enlargement of representative nuclei from both species (framed in blue in A and B). Scale bars = 5 µm.

### SUPPLEMENTAL FIGURE S28

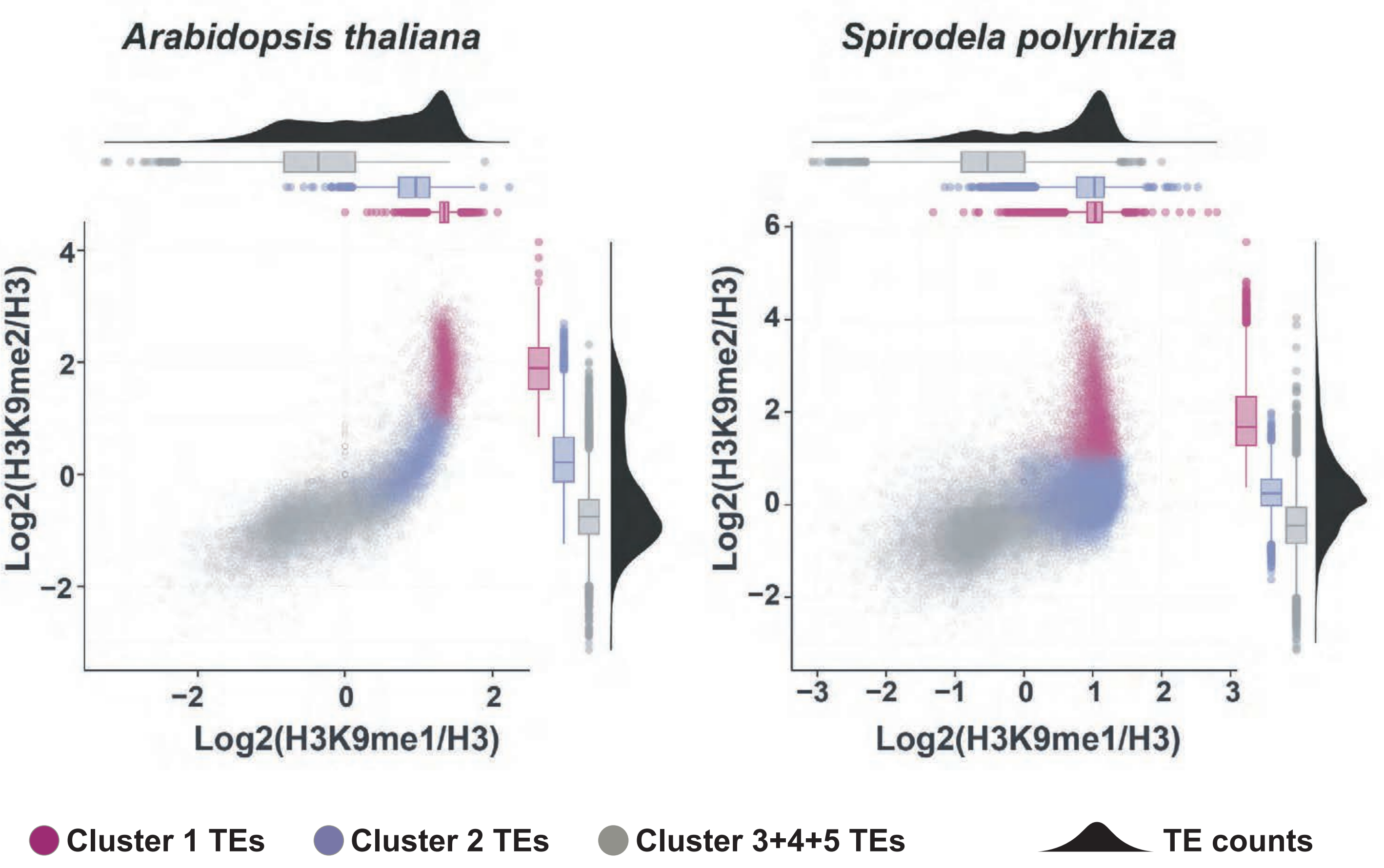

**Supplemental Figure S28: Distribution of H3K9me1/2 in Spirodela TEs.**
Scatterplots of H3K9me1 and H3K9me2 enrichment at *Spirodela* and *Arabidopsis* TEs colored by cluster (as defined in Figure 5).

### SUPPLEMENTAL FIGURE S29

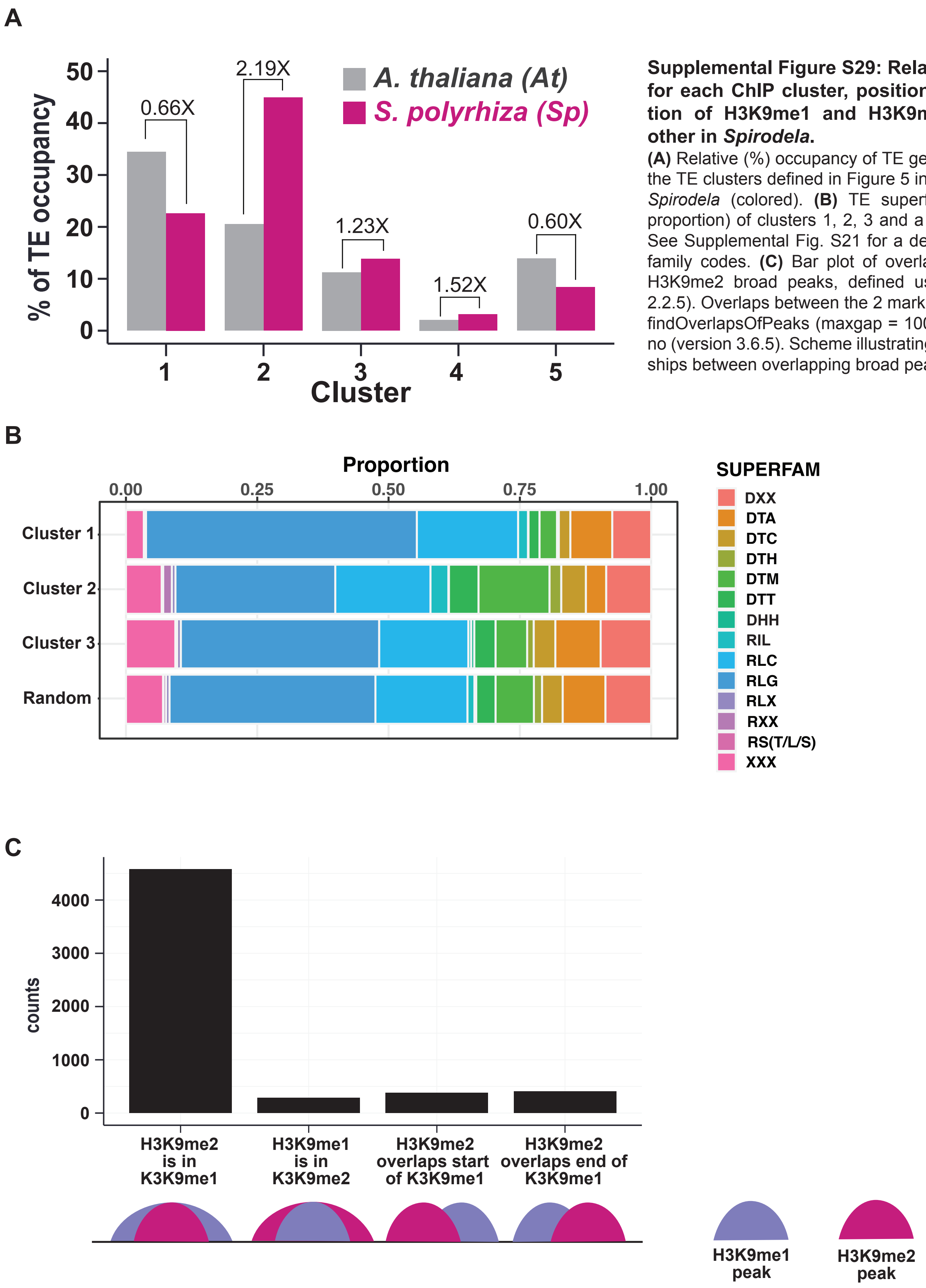

**Supplemental Figure S29: Relative TE occupancy for each ChIP cluster, position and TE composition of H3K9me1 and H3K9me2 peaks to each other in *Spirodela*.**
**(A)** Relative (%) occupancy of TE genomic space of each of the TE clusters defined in Figure 5 in *Arabidopsis* (grey) and *Spirodela* (colored). **(B)** TE superfamily composition (as proportion) of clusters 1, 2, 3 and a random subsampling. See Supplemental Fig. S21 for a description of TE superfamily codes. **(C)** Bar plot of overlapping HeK9me1 and H3K9me2 broad peaks, defined using MACS2 (version 2.2.5). Overlaps between the 2 marks were identified using findOverlapsOfPeaks (maxgap = 1000) from ChIPpeakAnno (version 3.6.5). Scheme illustrating the possible relationships between overlapping broad peaks.

### SUPPLEMENTAL FIGURE S30

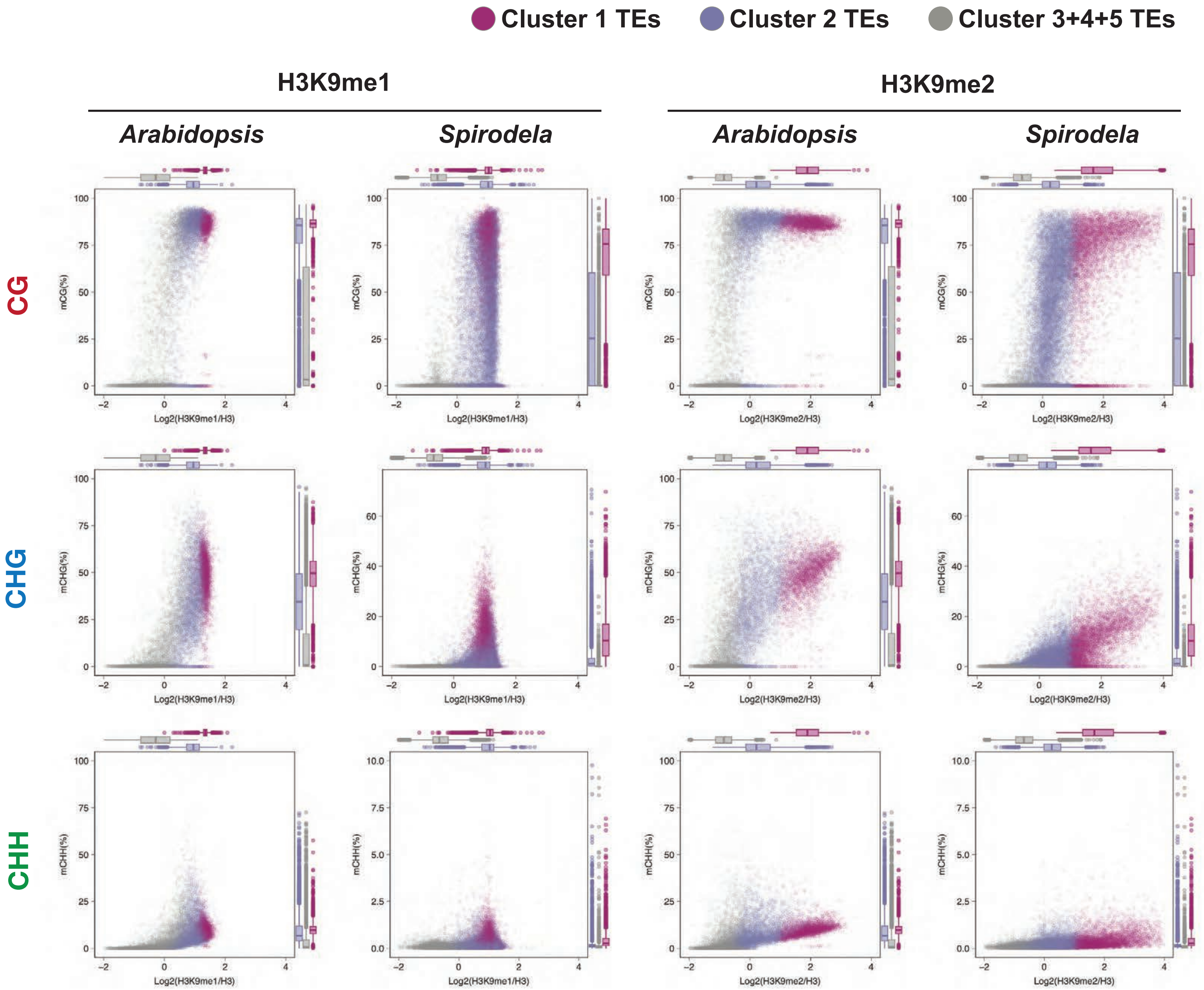

**Supplemental Figure S30: Analysis of H3K9me1/2 association to DNA methylation in *Arabidopsis* and *Spirodela*.** Scatter plots of DNA methylation in all three contexts against H3K9me1 and H3K9me2 enrichments of TEs in Cluster 1, 2 and 3/4/5 combined (as defined in Figure 5) in *Spirodela* and *Arabidopsis*.

### SUPPLEMENTAL FIGURE S31

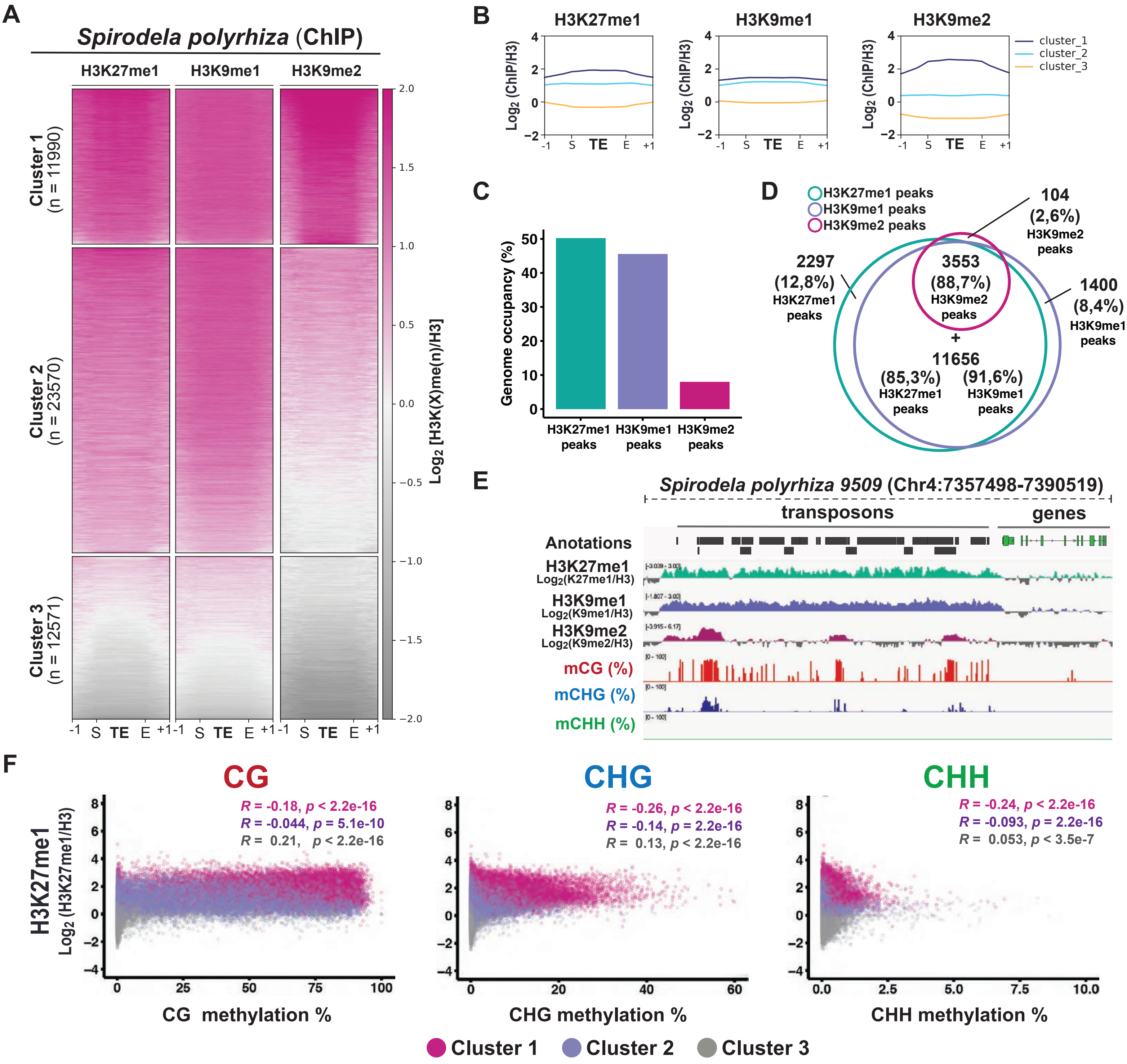

**Supplemental Figure S31: Patterns of H3K27me1 and H3K9me1/2 distribution in *Spirodela polyrhiza* transposons.**
**(A)** Heatmap of *k*-means clustering of transposons in *Spirodela* using H3K27me1, H3K9me1 and H3K9me2 enrichment over H3, presented as $log_2$[ChIP(X)/ChIP(H3)]. Enrichment is represented by color, while depletion is shown in gray. Each row corresponds to a length-normalized TE annotation ± 1 kb. Transposons are grouped into clusters 1 to 3 based on the abundance of examined marks. **(B)** Metaplots of average weighted H3K27me1, H3K9me1 and H3K9me2 enrichments over H3 at Cluster 1, 2 and 3 transposons ± 1 kb in *Spirodela*. **(C)** Genome occupancy (%) of H3K27me1, H3K9me1 and H3K9me2 in the *Spirodela* genome. **(D)** Venn diagram of the overlap between H3K27me1, H3K9me1 and H3K9me2 peaks along *Spirodela* genome. **(E)** Genome browser capture of the distribution of H3K27me1, H3K9me1 and H3K9me2 enrichments, together with DNA methylation, along genes and transposons in *Spirodela*. **(F)** Scatter plots of DNA methylation in all three contexts against H3K27me1 enrichment of TEs in *Spirodela*. *R* and *p* indicate Pearson correlation coefficients and p-values.

### SUPPLEMENTAL FIGURE S32

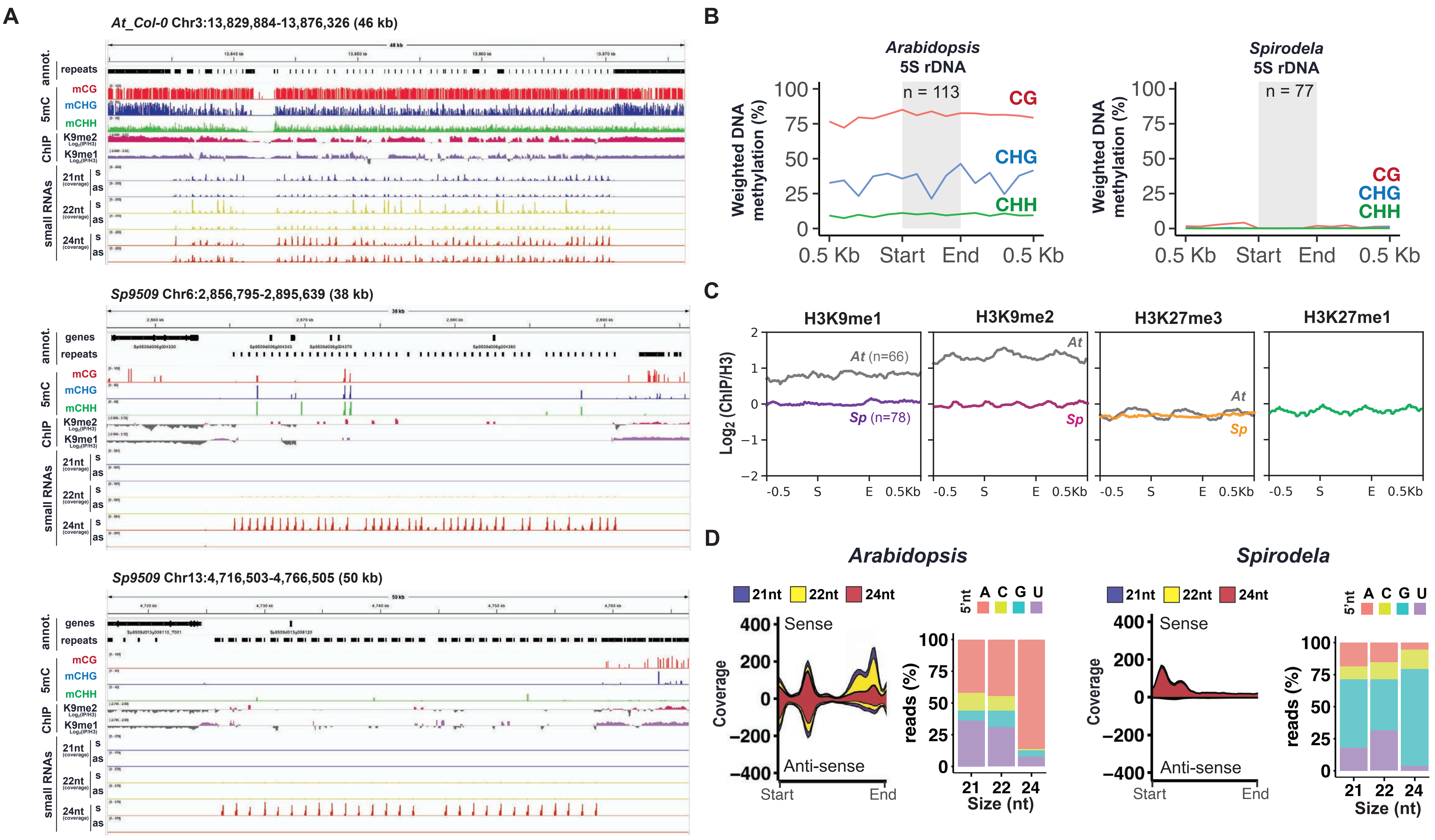

**Supplemental Figure S32: Analysis of the epigenetic and small RNA landscape of *5S rRNA* repeats in *Spirodela*.**
**(A)** Genome browser captures of *Arabidopsis Col-0 5S rDNA* array in Chromosome 3 and the two *Spirodela 9509 5S rDNA* arrays in Chromosomes 6 and 13. From top to bottom tracks display: gene and TE annotations, mCG, mCHG and mCHH methylation; H3K9me2 and H3K9me1 enrichment profiles; sense and anti-sense (relative to the *5S rDNA* annotations) 21-, 22- and 24-nt siRNAs. **(B)** Metaplots of averaged weighted DNA methylation in all three contexts over *5S rDNA* annotations ± 0.5 kb flanking regions in *Arabidopsis* and *Spirodela*. n indicates the number of 5S rDNA loci for which enough information (coverage) was available for analysis **(C)** Metaplots of average weighted H3K9me1, H3K9me2, H3K27me3 and H3K27me1 (only for *Spirodela*) enrichments over H3 at *5S rDNA* annotations ± 0.5 kb in *Arabidopsis* (grey line) and *Spirodela* (colored line). n indicates the number of *5S rDNA* loci for which enough H3 ChIP coverage (> 5 reads) was available for analysis. **(D)** Stacked distribution of 21-22-24-nt siRNAs and their 5'-nucleotide composition mapping to *5S rDNA* in *Arabidopsis* and *Spirodela*.

### SUPPLEMENTAL FIGURE S33

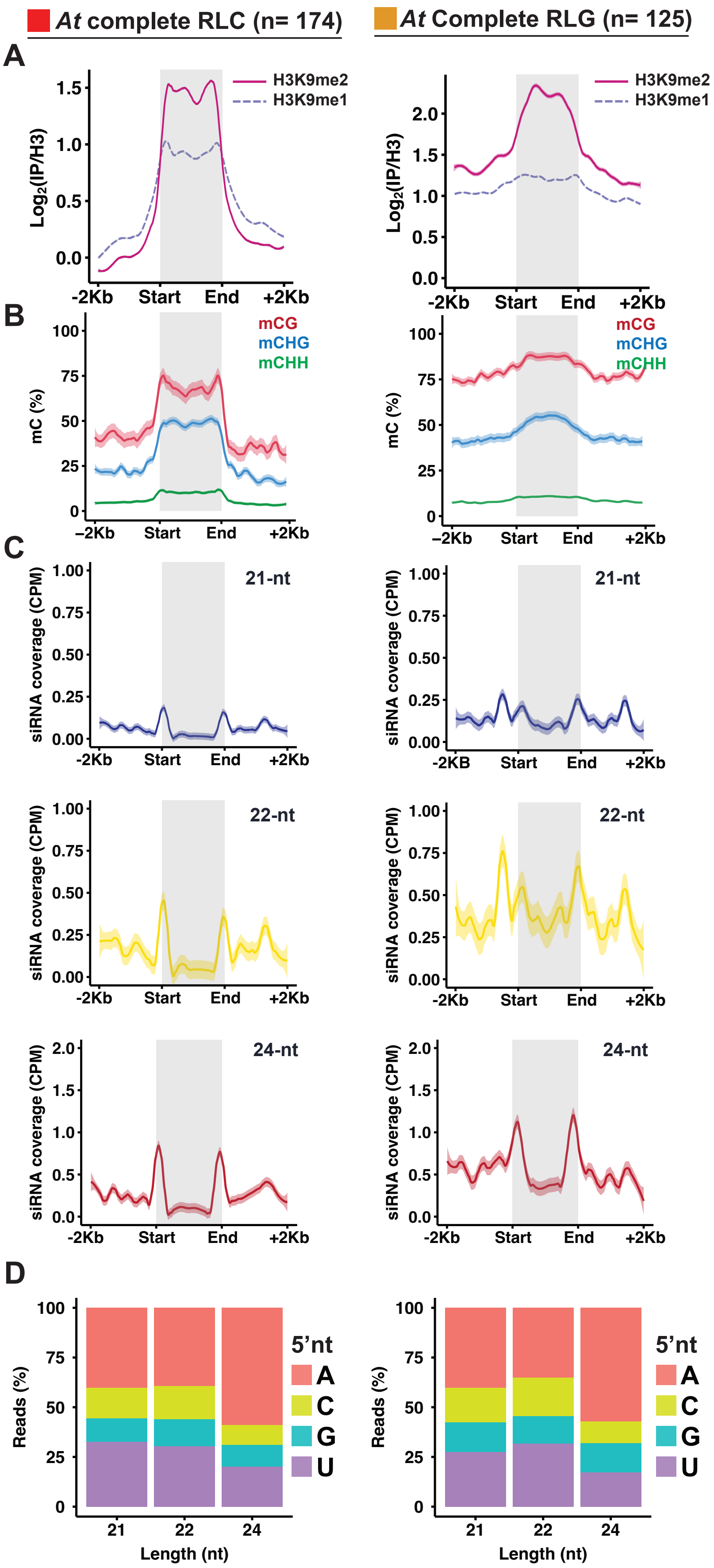

**Supplemental Figure S33: Distribution of silencing epigenetic marks and small RNAs in *Arabidopsis* RLC and RLG complete TEs.**

**(A)** Metaplots of average weighted H3K9me1 (solid line) and H3K9me2 (dashed line) enrichments over H3 at complete RLC (left) and RLG (right) transposon annotations ± 2 kb in Arabidopsis. Complete TEs were defined as those for which all RLC and RLG protein domains where identified. n indicates number of complete elements in each superfamily. **(B)** Metaplots of averaged weighted DNA methylation in all three contexts over RLC and RLG transposons ± 2 kb flanking regions in *Arabidopsis*. **(C)** Metaplots of averaged weighted 21-, 22- and 24-nt small RNA abundance over complete RLC and RLG ± 2 kb flanking regions in *Arabidopsis*. In all metaplots (A-D), solid lines represent the mean and colored shadows the standard deviation of the mean. **(D)** 5'-nucleotide (5'nt) composition distribution (as % of reads) of 21-, 22- and 24-nt siRNAs mapping to intact RLC and RLG ± 2 kb flanking regions in *Arabidopsis*. d 5'-nucleotide (5'nt) composition distribution (as % of reads) of 21-, 22- and 24-nt siRNAs mapping to intact RLC and RLG ± 2 kb flanking regions in *Arabidopsis*.

### SUPPLEMENTARY FIGURE S34

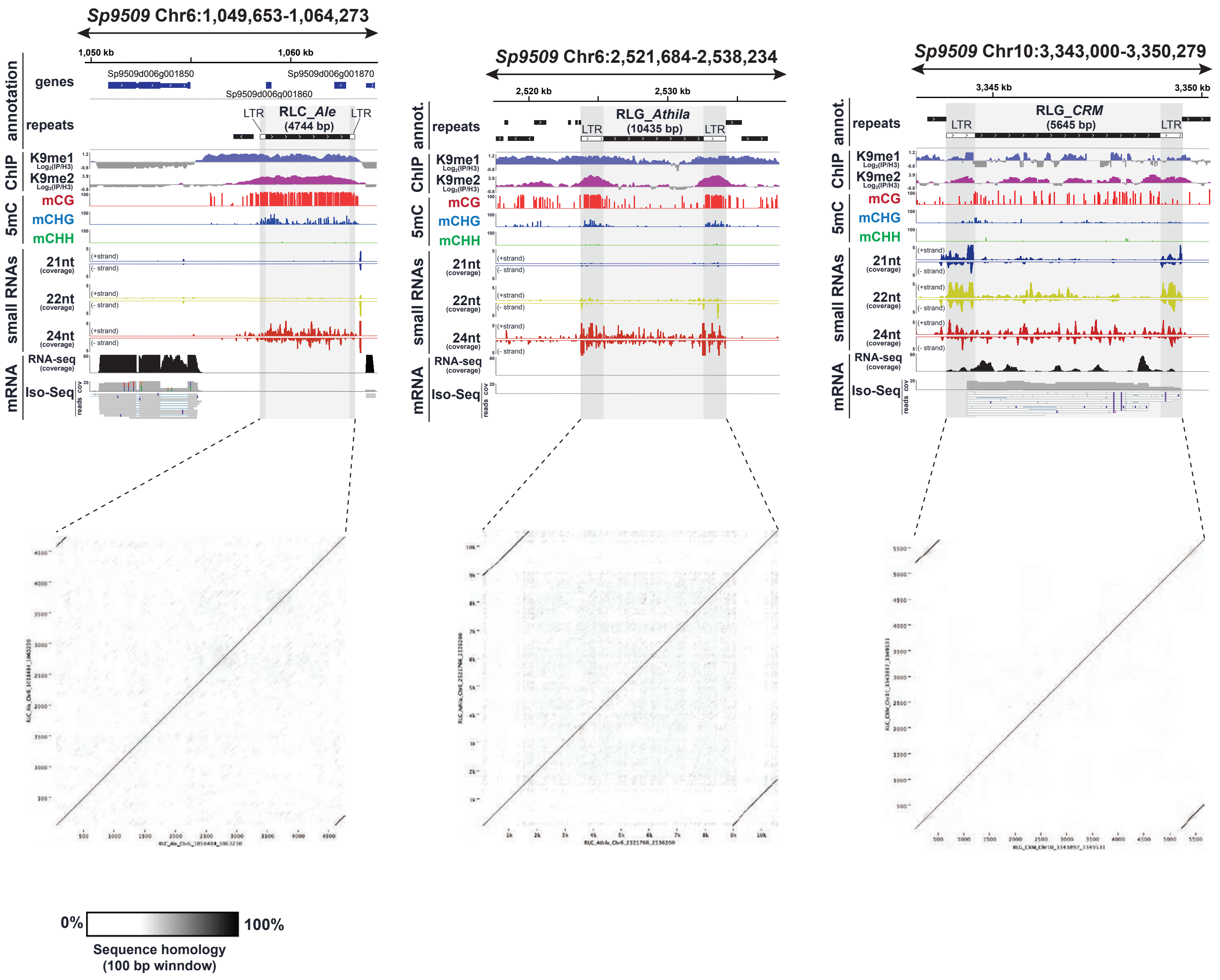

**Supplementary Figure S34: LTR identification for complete TEs in Figure 6.**
To identify LTRs of complete TEs, dot plots of their sequences against themselves were generated to identify the direct repeats. Dot plots were generated using a 100 bp window sliding at single bp intervals.

### SUPPLEMENTAL FIGURE S35

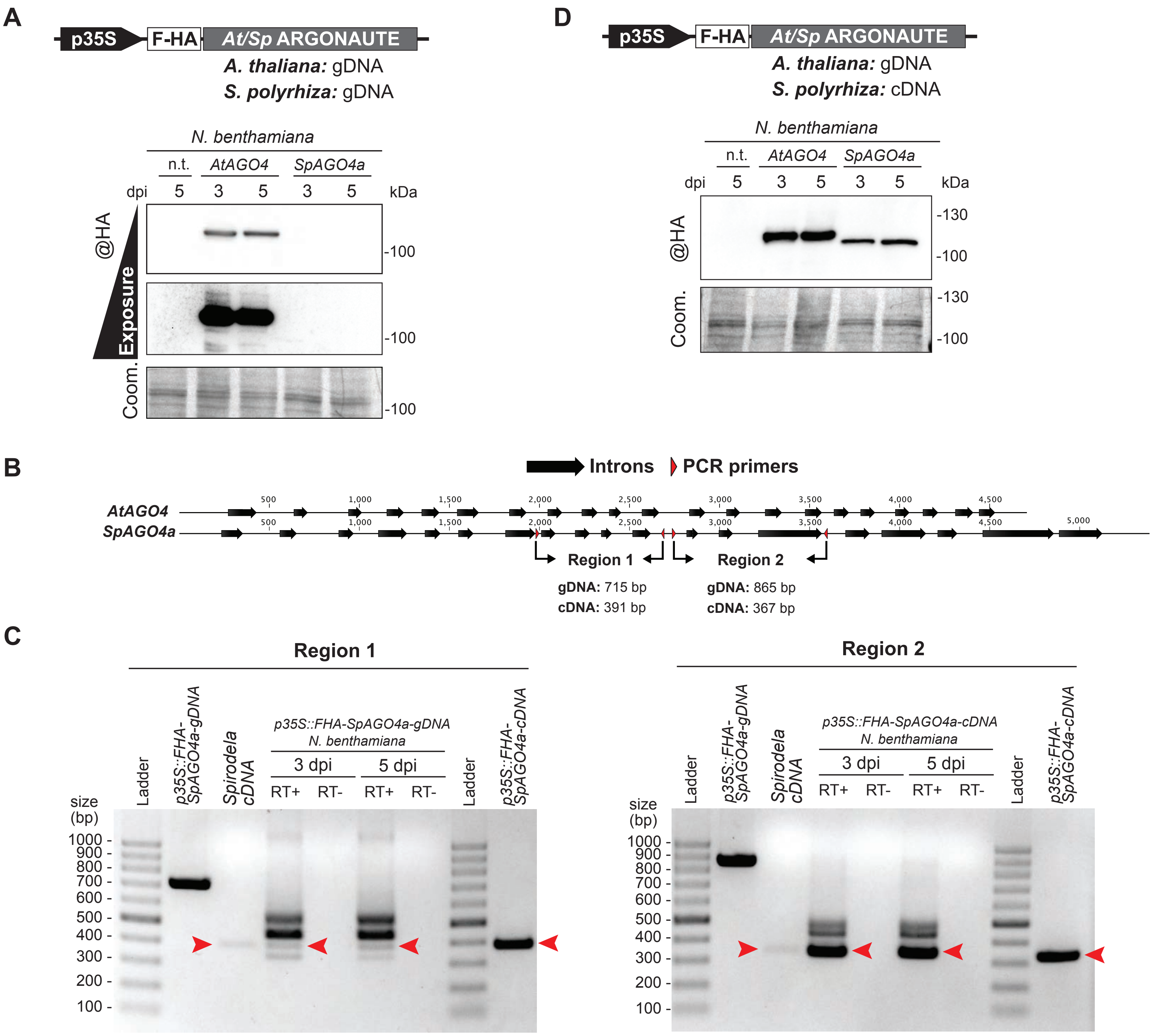

**Supplemental Figure S35: Defects in *SpAGO4a* splicing in *N. benthamiana* prevent its transient expression from gDNA.**
**(A)** Protein blot analysis of *Arabidopsis* and *Spirodela* AGO4 (*AtAGO4* and *SpAGO4a*) gDNA transient expression in *N. benthamiana* at 3 and 5 days post infiltration (dpi). n.i.: non-infiltrated. Coomassie (Coom.) staining of blot is shown as loading. Scheme of constructs infiltrated in *N. benthamiana* leaves for transient expression is shown above. p*35S*: CaMV 35S promoter; F-HA: in-frame N-term Flag and HA peptide tags. Genomic (gDNA) or cDNA origin of the *AGO4* sequences is indicated. **(B)** Schematic representation of *AtAGO4* and *SpAGO4a* loci. Introns are represented by arrows. Position of PCR primers used to determine efficiency and completion of splicing across indicated regions are indicated by red triangles. Expected amplicon sizes in each case is indicated below **(C)** Agarose gel electrophoresis of RT-PCRs of regions in (B) of Sp*AGO4a* transient expression in *N. benthamiana*. Arrowheads indicated the expected size of properly spliced mRNA as expected from in silico predictions and validated by RT-PCR in *Spirodela* cDNA. Plasmid DNA containing *SpAGO4a* gDNA is used as control for unspliced PCR, Spirodela total cDNA and plasmid containing Sp*AGO4a* cDNA are used as positive controls for properly spliced mRNA size. **(D)** Protein blot analysis of *AtAGO4* gDNA and *SpAGO4a* cDNA transient expression in *N. benthamiana* at 3 and 5 days post infiltration (dpi). n.i.: non-infiltrated. Coomassie (Coom.) staining of blot is shown as loading. Scheme of constructs infiltrated in *N. benthamiana* leaves for transient expression is shown above. p*35S*: CaMV 35S promoter; F-HA: in-frame N-term Flag and HA peptide tags. Genomic (gDNA) or cDNA origin of the *AGO4* sequences is indicated.

### SUPPLEMENTAL FIGURE S36

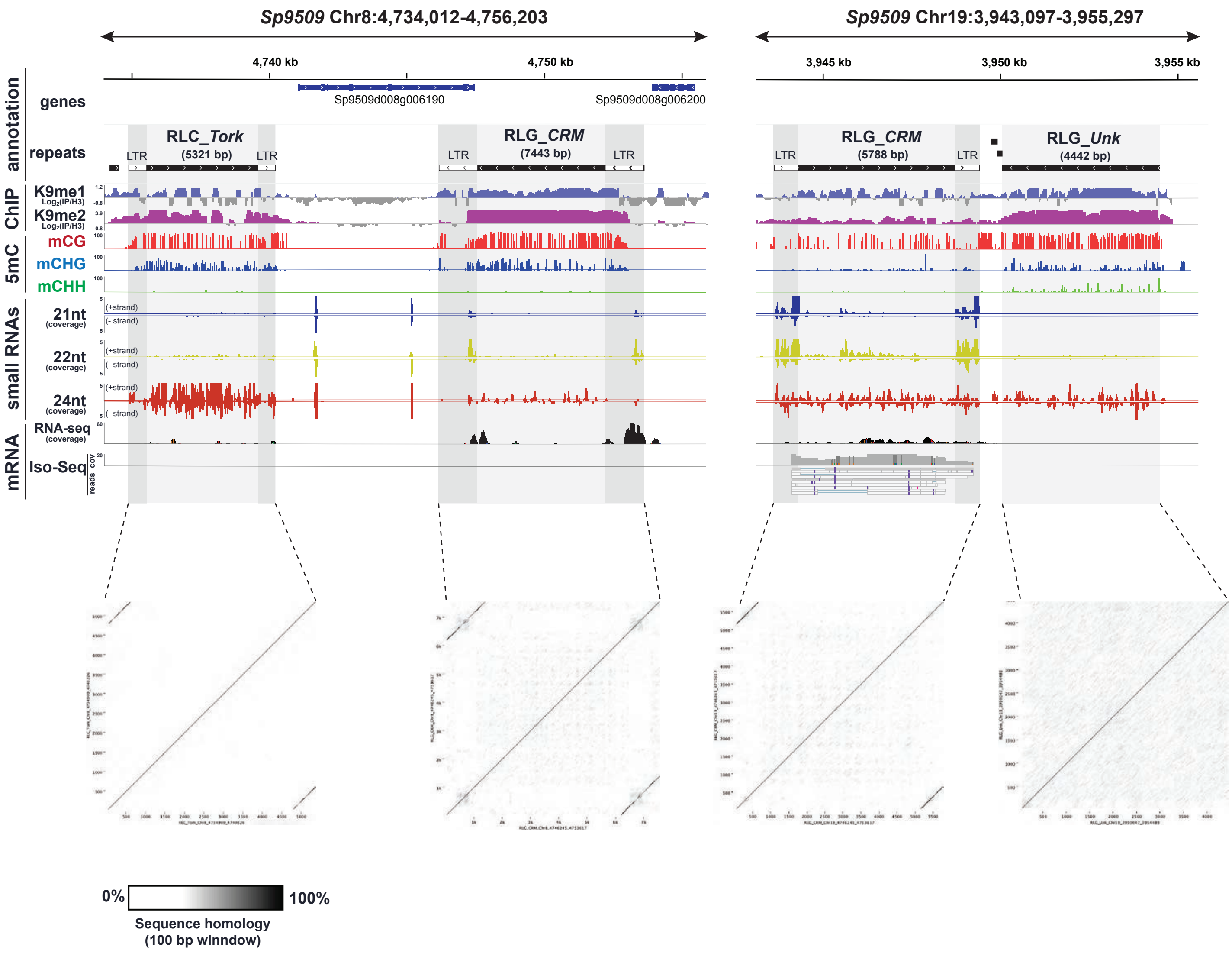

**Supplemental Figure S36: Additional examples of siRNA producing TEs in *Spirodela*.**
Genome browser captures of selected complete TEs in *Spirodela*. From top to bottom tracks display: gene and TE annotations, H3K9me1 and H3K9me2 enrichment profiles; mCG, mCHG and mCHH methylation; 21-, 22- and 24-nt siRNA mapping to Watson (+) and Crick (-) DNA strands; RNA expression by short (illumina) and long reads (Iso-Seq). Superfamily, clade and length (in bp) of TEs is indicated above their annotation. Light and dark gray shadow delimit TEs and LTRs respectively across all tracks. Dot plots of their sequences against themselves were generated to identify the direct repeats using a 100 bp window sliding at single bp intervals.

### SUPPLEMENTAL FIGURE S37

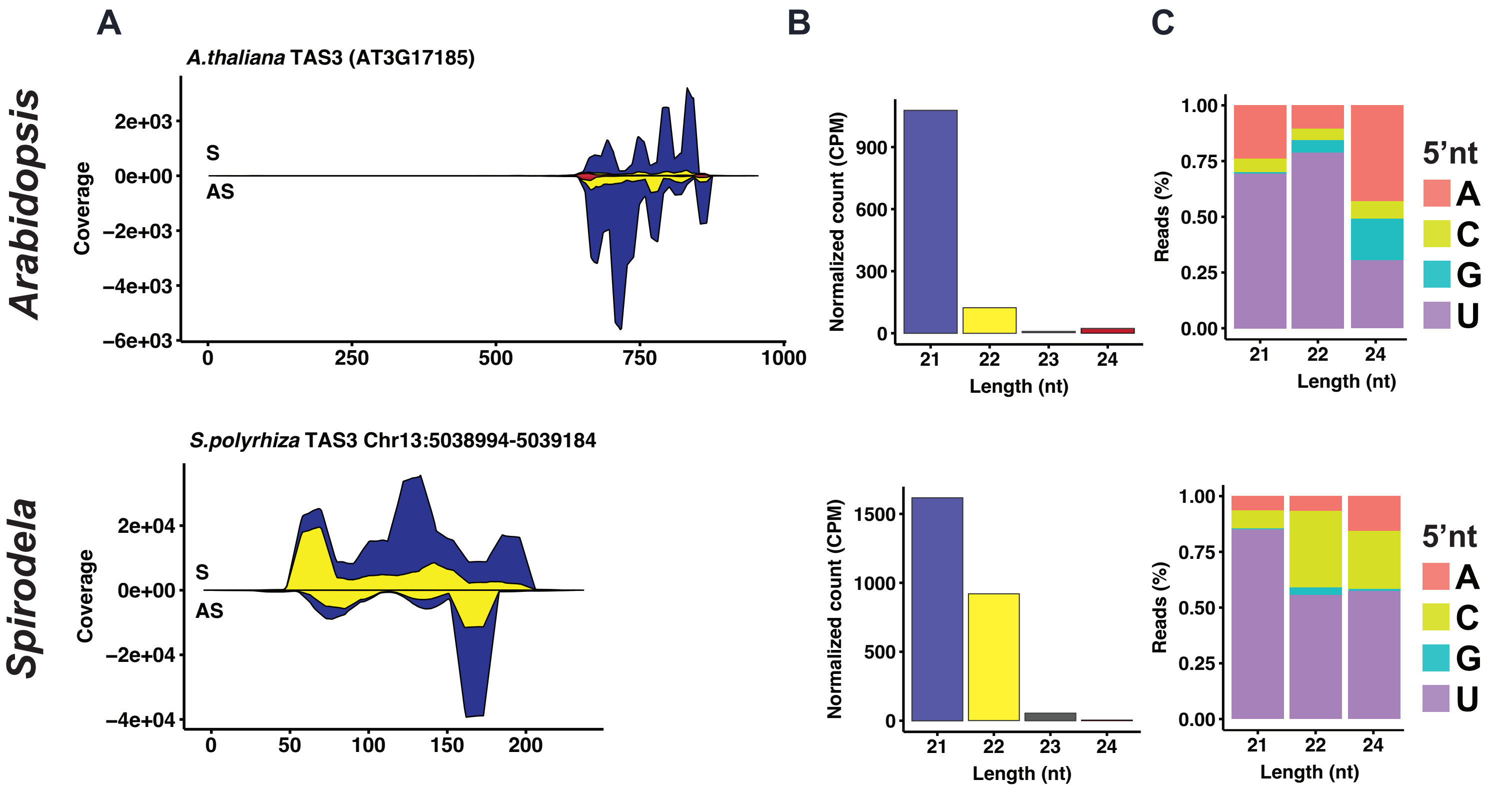

**Supplemental Figure S37:** ***TAS3*** **tasiRNAs in** ***Spirodela*****.**
**(A)** Stacked distribution of sense (S) and antisense (AS) *TAS3* tasiRNA in *Arabidopsis* and *Spirodela*. **(B)** Size distribution and abundance of *TAS3* tasiRNA in *Spirodela* and *Arabidopsis*. **(C)** 5’-nucleotide (5’nt) composition distribution of 21-, 22- and 24-nt siRNAs mapping *TAS3* in *Spirodela* and *Arabidopsis*.

### SUPPLEMENTAL FIGURE S38

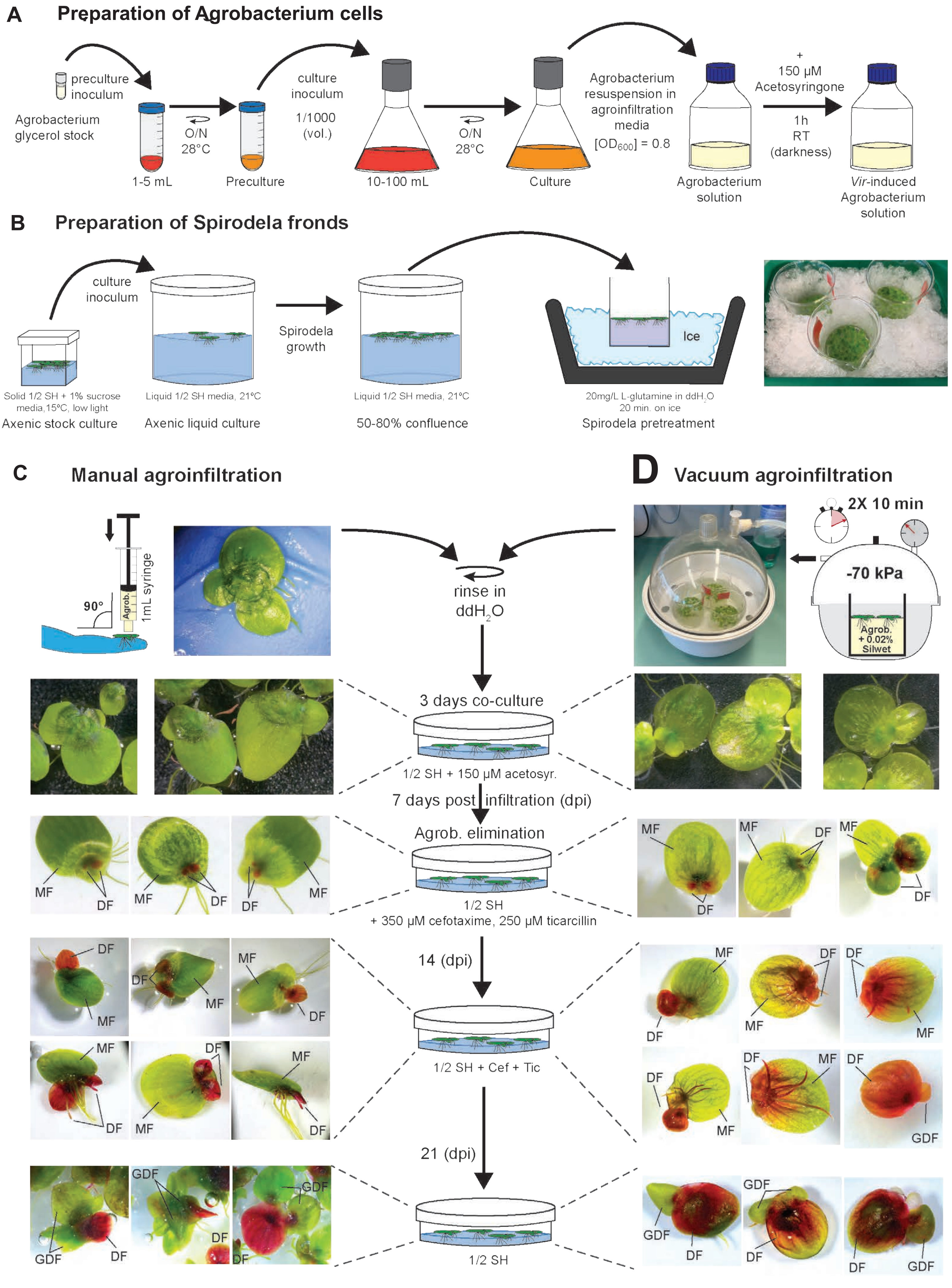

Legend in next page

**Supplemental Figure S38: Transient expression procedure in *Spirodela*.**
**(A)** For *Agrobacterium*-mediated transient expression in *Spirodela* standard procedures for the preparation and induction of *Agrobacterium* cells carrying plant binary vectors with the desired construct were followed. **(B)** *Spirodela* cultures are started from axenic stock cultures and grown to 50-80% confluence as saturated cultures are more recalcitrant to the procedure. Prior agroinfiltration, fronds are pretreated in a cold solution of 20mg/L of L-glutamine on ice for 20 min. Induced *Agrobacterium* cultures can then be infiltrated into *Spirodela* fronds manually (C) or applying vacuum (D). **(C)** Procedure for manual agroinfiltration. Bacterial solutions are infiltrated locally above the budding pockets. **(D)** Procedure for vacuum infiltration. *Spirodela* cultures are placed on bacteria solutions in a vacuum bell and vacuum is applied twice. In both procedures, excess of Agrobacterium is eliminated before plants are co-cultured with the bacteria. Then *Agrobacterium* is eliminated from media with antibiotics and plants are cultured normally. While manual infiltration yields a more uniform transient expression in developing fronds at the infiltrated area (C), it is more tedious and less plants can be processed. Vacuum infiltration on the other hand allows for the infiltration of a greater number of plants, however the resulting expression is less homogeneous. In both cases daughter fronds appear to express uniformly transgenic reporters, their progeny does not express it, hence we considered the system to be transient .MF: mother frond; DF: daughter frond; GDF: grand-daughter frond.

### SUPPLEMENTAL FIGURE S39

| Name | Sequence 5'->3' | Use | Note |
|---|---|---|---|
| **FLAG F** | GAAACGACAATCTGATCCAAGCTCAAGCTAAGCTTGAGCTCCAACTTTGTATAG | FLAG Gibson cloning | |
| **FLAG R** | GTAGGATCCGCCACCTCCCTTGTCATCGTCATCCTTGTAATCCATGAAGCCTGCTTTTTTGTACAAACTTGCTG | FLAG Gibson cloning | |
| **HA F** | GGCGGATCCTACCCATACGATGTTCCAGATTACGCTGGAGGcGGTCTAGATACCCAGCTTTCTTGTACAAAGTGG | HA cloning | |
| **HA R** | CAGACCCGGGAGGTCACTGGATTTTGGTTTTAGG | HA cloning | |
| **AtAGO4_gDNA F** | *CATACGATGTTCCAGATTACGCTGGAGGCGGT***GATTCAACAAATGGTAACGGAGC** | At*AGO4*_gDNA Gibson cloning | Homologous sequence to backbone for Gibson assembly in italics. In bold At*AGO4* sequence. |
| **AtAGO4_gDNA R** | *CACCACTTTGTACAAGAAAGCTGGGTATCTAG***TTAACAGAAGAACATGGAGTTGGC** | At*AGO4*_gDNA Gibson cloning | Homologous sequence to backbone for Gibson assembly in italics. In bold At*AGO4* sequence. |
| **SpAGO4_gDNA F** | *GCAACTAGTG***GAAAAACCAGAGGGGGAAAGAACAG** | Sp*AGO4*_gDNA BcuI cloning | BcuI RE site italics. In bold Sp*AGO4* sequence. |
| **SpAGO4_gDNA R** | *GCAACTAGT***TCAGCAGAAGAACATGGAACTGCTC** | Sp*AGO4*_gDNA BcuI cloning | BcuI RE site italics. In bold Sp*AGO4* sequence. |
| **SpAGO4_cDNA F** | *GCAACTAGTG***CCCATGAAAAGACCAGGAACGGGAA** | Sp*AGO4*_cDNA BcuI cloning | BcuI RE site italics. In bold Sp*AGO4* sequence. Primer match both SpAGO4 in Spirodela |
| **SpAGO4_cDNA R** | *GCAACTAGT***TCAGCAGAAGAACATGGAGCTGCTC** | Sp*AGO4*_cDNA BcuI cloning | BcuI RE site italics. In bold Sp*AGO4* sequence. Primer match both SpAGO4 in Spirodela |
| **SpAGO4a r1 F** | **TCCCCAAGAGAGGATGAGAG** | RT-PCR | See Supplemental_Fig_S35 |
| **SpAGO4a r1 R** | **GTCCACCCTAACATTTGG** | RT-PCR | See Supplemental_Fig_S35 |
| **SpAGO4a r2 F** | **CGCTTTCTTCTGTGTATTCTTC** | RT-PCR | See Supplemental_Fig_S35 |
| **SpAGO4a r2 R** | **GAGATATTGAAGGCCACTGCC** | RT-PCR | See Supplemental_Fig_S35 |
| **Scarlet probe F** | **ATGGTGAGCAAGGGTGAAGCCG** | Northern probe | |
| **Scarlet probe R** | **TTACTTGTAGAGTTCGTCCATAC** | Northern probe | |
| **RUBY probe F** | **CAACACCATCGAAGTAGTGC** | Northern probe | |
| **RUBY probe R** | **TACCACATCCTCCACATTCG** | Northern probe | |
| **miR159a probe** | **TAGAGCTCCCTTCAATCCAAA** | Northern probe | |

**Supplemental Figure S39: Oligos used in this study.**

### Supplemental Methods

**Plant material description and growth conditions:**

***Spirodela polyrhiza* sterilization and stocks.** Upon arrival, the initial culture was amplified and further sterilized to obtain axenic cultures. In brief, 50 fronds of *Spirodela* were incubated in 10 ml of a 0.5% DanKlorix® solution (Colgate-Palmolive; commercial detergent containing 2.8% Na-Hypochlorite) for 1 minute under agitation. Subsequently, the sterilizing solution was removed, and the fronds were washed three times with sterile water before being transferred to individual wells in 6-well plates containing 6 ml of sterile ½ SH + 2% sucrose media. After one week, fronds in wells with no visible growth of microorganisms were transferred to plates with solid (2% Gelrite™, Duchefa Biochemie, #G1101) ½ SH medium + 10% sucrose for further confirmation of absence of contamination. For long term storage, fronds were grown on solid ½ SH medium + 10% sucrose in Magenta™ GA-7 containers, in a plant growth cabinet (SEQI-P4, Percival) in long day conditions (16h light /8h dark) at 15°C, with low light (intensity of 19.5 µM $m^{-2}$ $s^{-1}$). Axenic stocks were sub-cultured every 6 months.

***Arabidopsis***: *Arabidopsis thaliana* ecotype *Columbia*-0 (Col-0, salk_CS60000) was used as WT. The PolIV mutant, *nrpd1a-4* (salk_083051) was previously described (Kanno et al. 2005). These lines were provided by The Nottingham *Arabidopsis* Stock Centre (NASC) (http://arabidopsis.info/). For all experiment with *Arabidopsis*, seeds were surface sterilized and sown in half-strength Murashige and Skoog (1/2 MS) media, (Duchefa Biochemie, #M0231) solid medium plates, stratified in the dark at 4°C for 3 days, and transferred to a growth chamber with long-day conditions (16h light /8h dark) at 21°C with a LED light intensity of 85 µM $m^{-2}$ $s^{-1}$. Seedlings were then harvested and flash-frozen. When needed, seedlings were transferred to soil and grown under long-day conditions (16h light /8h dark) at 21°C, 60% humidity with a LED light intensity of 89 µM $m^{-2}$ $s^{-1}$.

***Nicotiana bethamiana***: After 7 days at 4°C in the dark, Nicotiana seeds were sown in soil, germinated, and grown under long-day conditions (16h light /8h dark) at 21°C, 60% humidity with a LED light intensity of 89 µM $m^{-2}$ $s^{-1}$. They were used for agroinfiltration experiments once they have developed 4-6 true leaves. WT and *rdr6i* mutant lines seeds were previously described (Schwach et al. 2005).

**Plant imaging:**

Pictures of plants were taken with a Canon EOS 80D camara or with an Andonstar AD409 digital microscope. Scans of *N. benthamiana* leaves and *Spirodela* cultures in dishes were performed with an Epson Perfection V600 photo scanner.

**Phylogenetic analysis and protein domain annotation:**

To identify *Spirodela* orthologues and paralogues, DNA and protein sequences from *Arabidopsis* genes involved in silencing were retrieved from TAIR10 and blasted against *Spirodela* protein database (BLAST+(Camacho et al. 2009) version 2.8.1, blastp -qcov_hsp_perc 60 or 30) or against the *Spirodela* genome (BLAST+ version 2.8.1, tblastn with deafult parameters) to retreive as may putative orthologues as possible. Candidate genes and proteins were sorted based on the presence on known domains and orthogroups formed based on known orthologues from other angiosperms. Protein sequences from several angiosperms were obtained from the EBI-EMBL Uniprot databse (Consortium et al. 2022) (https://www.uniprot.org) based on blast searches against known proteins off each family and orthogroup. Protein alignments were performed using CLC Main Workbench 20 (QIAGEN) with Gap open cost = 10; Gap extension cost = 1 parameters. Using the same software, alignments were then used to generate unrooted trees with the Neighbor Joining method and Jukes-Cantor substitution model for protein distance. Maximum likelihood trees were constructed with 1000 bootstrap replicates. Domain annotations were carried as well in CLC with the Pfam-A v35 database (Mistry et al. 2020) and further validated and expanded by independent annotation using predictions with the Interpro database (Paysan-Lafosse et al. 2022).

**Protein extraction and western blot:**

**Protein extraction:** For protein extraction from *Arabidopsis* and *Spirodela*, plant material was flash-frozen upon harvesting and ground with a Silamat S6 as described in the RNA extraction section. For N. benthamiana, leaf tissue was grounded in liquid nitrogen using mortar and pestle. 100mg of ground tissue was mixed and homogeinized for 5 min. with 500µl of extraction buffer (0.7 M sucrose, 0.5 M Tris-HCl, pH 8, 5 mM EDTA, pH 8, 0.1 M NaCl, 2% b-mercaptoethanol) and cOmpleteTM mini EDTA-free protease inhibitor cocktail (Roche, #04693159001) following manufacturer's instructions. 1 volume of TRIS-buffered phenol (pH 8) was added and incubated with agitation for 5 minutes. The upper phenolic phase was recovered after centrifugation (12000g for 10 min at 4ºC) and proteins precipitated with 5 volumes of 0.1 M ammonium acetate dissolved in methanol at -20oC overnight. Following centrifugation (5000g for 15 min at 4ºC), the resulting protein pellet obtained was washed twice with 0.1 M ammonium acetate dissolved in methanol and resuspended in 50-100 µL of resuspension buffer (3% SDS, 62.3 mM Tris-HCl, pH 8, 10% glycerol).

**Protein blots:** Western blot was carried following standard denaturing SDS-PAGE and blotting procedures (Sambrook and Russell 2001). Unless otherwise specified, 150µg of protein were resolved on SDS-polyacrilamide gels, transferred onto Inmobilon-P PVDF membrane (Millipore, # IPVH00010) and incubated with the appropiate antibodies (see below) in PBS with 0.1% Tween-20 and 5% non-fat dried milk. The primary antibody was incubated overnight at 4oC, and the HRP-conjugated secondary antibody for 1 hour at room temperature. Detection was performed using Clarity Max™ Western ECL Substrate (Biorad, #1705062), and imaged with the ChemiDoc Imaging system (Bio-rad). Following detection, loading was verified by coomassie staining of membranes.

**Primary antibodies used for western blot:** monoclonal (mouse) anti-HA tag [12CA5] (Abcam, #ab1424), dilution 1:10000. Polyclonal (rabbit) anti-H3 (Abcam #ab1791), dilution 1:10000. Polyclonal (rabbit) anti-H3K9me1 (Abcam #ab8896), dilution 1:2000. Monoclonal (mouse) anti-H3K9me2 (Abcam #ab1220), dilution 1:2000.

**Secondary antibodies used for western blot:** Polyclonal (goat) anti-mouse HRP-conjugated (ThermoFisher, #62-6520), dilution 1:10000. Polyclonal (goat) anti-rabbit HRP-conjugated (ThermoFisher, #65-6120), dilution 1:10000.

**Histone PTM analysis by mass spectrometry:**

**Sample preparation:** Nuclear isolation and acid-extraction of histones was performed from 2gr of *Arabidopsis* 12 days-old seedlinngs and *Spirodela* fronds following the protocol described in (Scheid et al. 2022). To check for quality of the samples, an aliquot corresponding to 5% of the extracted protein samples were run on a denaturing 15% SDS-PAGE stained with coomassie for visualization of bands. Protein samples were brought to 40µL using a vacuum concentrator and 20 µl of each histone sample was supplemented with 2 µl of 1 M triethylammonium bicarbonate (TEAB). Propionic anhydride (Sigma-Aldrich, Nr 240311) was freshly diluted in a ratio of 1:100 with molecular-grade $H_2O$, and 2 µl of the dilution were added to each sample and incubated at room temperature for 2 min. The reaction was quenched by adding 2 µl of 80 mM hydroxylamine (Thermo Scientific, #90115) and incubation at 37°C for 20 min. Histones were digested by adding 250 ng trypsin (Trypsin Gold, Promega #V5280) and incubating at 37°C overnight. For the second propionylation on peptide level, 6µl of a fresh Propionic anhydride 1:100 dilution was added to the sample which was again incubated for 2min at room temperature before being quenched with 3µl of 80 mM hydroxylamine as described above. Samples were acidified by adding 50 µL of 5% formic acid and desalted on C18 (Empore, #2215-C18) StageTips as in (Rappsilber et al. 2007).

**nanoLC-MS/MS Analysis:** The nano HPLC system (UltiMate 3000 RSLC nano system, ThermoFisher Scientific) was coupled to a Q Exactive HF-X or an Orbitrap Exploris 480 mass spectrometer equipped with a Nanospray Flex ion source (all parts ThermoFisher Scientific). Peptides were loaded onto a trap column (PepMap Acclaim C18, 5 mm × 300 µm ID, 5 µm particles, 100 Å pore size, ThermoFisher Scientific) at a flow rate of 25 µl/min using 0.1% TFA as mobile phase. After loading, the trap column was switched in line with the analytical column (PepMap Acclaim C18, 500 mm × 75 µm ID, 2 µm, 100 Å, ThermoFisher Scientific). Peptides were eluted using a flow rate of 230 nl/min, starting with the mobile phases 98% A (0.1% formic acid in water) and 2% B (80% acetonitrile, 0.1% formic acid) and linearly increasing to 45% B over the next 120 min. This was followed by a steep gradient to 95% B in 5 min, stayed there for 5 min and ramped down in 2 min to the starting conditions of 98% A and 2% B for equilibration at 30°C. The Q Exactive HF-X mass spectrometer was operated in data-dependent mode, using a full scan (m/z range 380-1500, nominal resolution of 60,000, target value 1e6) followed by MS/MS scans of the 10 most abundant ions. MS/MS spectra were acquired using normalized collision energy of 28, isolation width of 1.0 m/z, resolution of 30.000, a target value of 1e5 and maximum fill time of 105 ms. Precursor ions selected for fragmentation (exclude charge state 1, 7, 8, >8) were put on a dynamic exclusion list for 20s. Additionally, the minimum AGC target was set to 5e3 and intensity threshold was calculated to be 4.8e4. The peptide match feature was set to preferred and the exclude isotopes feature was enabled. The Orbitrap Exploris 480 mass spectrometer was operated in data-dependent mode 'Cycle Time', performing a full scan (m/z range 350-1200, resolution 60,000, normalized AGC target 100%, compensation voltage CV of -45V), followed by MS/MS scans of the most abundant ions for a cycle time of 2.0 seconds. MS/MS spectra were acquired using an isolation width of 1.2 m/z, normalized AGC target 200%, HCD collision energy of 30, orbitrap resolution of 30,000, maximum injection time of 100 ms and minimum intensity set to 50,000. Precursor ions selected for fragmentation (include charge state 2-6) were excluded for 20 s. The monoisotopic precursor selection (MIPS) mode was set to Peptide and the exclude isotopes feature was enabled.

**Peptide identification and data analysis:** For peptide identification, the RAW-files were loaded into Proteome Discoverer (version 2.5.0.400, Thermo Scientific). All hereby created MS/MS spectra were searched using MSAmanda v2.0.0.19924 (Dorfer et al. 2014). For the protein-identification-set, the peptide and fragment mass tolerance was set to ±10 ppm, the maximum number of missed cleavages was set to 2, using tryptic enzymatic specificity without proline restriction. Peptide and protein identification was performed in two steps. For an initial search the RAW-files were searched against the *Arabidopsis* database called TAIR10 (32,785 sequences; 14,482,855 residues) or the database called Spirodela_polyrhiza_3_9509 (28,688 sequences; 8,725,633 residues), supplemented with common contaminants and sequences of proteins of interest. The result was filtered to 1 % FDR on protein level using the Percolator algorithm (Käll et al. 2007) integrated in Thermo Proteome Discoverer. A sub-database was generated for further processing. For the second search, the RAW-files were searched against the created sub-database using the same settings as above, but considering 5 missed cleavages and the following additional variable modifications: oxidation on methionine, deamidation on asparagine and glutamine, acetylation on lysine, phosphorylation on serine, threonine and tyrosine, methylation on lysine and arginine, di-methylation on lysine and arginine, tri-methylation on lysine, propionyl on lysine, propionyl on peptide N-terminal, glutamine to pyro-glutamate conversion at peptide N-terminal glutamine and acetylation on protein N-terminus. The localization of the post-translational modification sites within the peptides was performed with the tool ptmRS, based on the tool phosphoRS (Taus et al. 2011). Identifications were filtered again to 1 % FDR on protein and PSM level, additionally an Amanda score cut-off of at least 150 was applied.Proteins were filtered to be identified by a minimum of 2 PSMs in at least 1 sample. Peptides were subjected to label-free quantification using IMP-apQuant (Doblmann et al. 2019). Protein areas have been computed by summing up all peptides. Resulting protein areas were normalised using iBAQ (Schwanhäusser et al. 2011). For relative quantification of H3K9 modifications, the area of the peptides carrying each modification was normalized by the sum of the areas of the same peptide.

The mass spectrometry proteomics data have been deposited to the ProteomeXchange Consortium via the PRIDE partner repository with the dataset identifier PXD050443(Perez-Riverol et al. 2021).

**Chromatin immunoprecipitation (ChIP):**

Chromatin immunoprecipitation (ChIP) was conducted on *Arabidopsis* 10-day-old seedlings (WT Col-0) and *Spirodela polyrhiza* #9509 fronds. Tissues (2g each) were fixed with 1% formaldehyde in PBS for 10 minutes at -70 kPa, and the cross-linking reaction was halted with 125 mM glycine. For nuclei isolation, the crosslinked tissues were frozen, ground into a fine powder in liquid nitrogen, and subsequently resuspended in M1 buffer (10 mM sodium phosphate pH 7.0, 100 mM NaCl, 1 M hexylene glycol, 10 mM 2-mercaptoethanol, and cOmplete$^{TM}$ mini EDTA-free protease inhibitor cocktail (Roche, #04693159001)). The suspension was filtered through a double layered miracloth (Merck, #475855-1r), prior nuclei precipitation by centrifugation for 5 min, 2,000*g* at 4ºC**,** followed by 5 rounds of washing using M2 buffer (10 mM sodium phosphate pH 7.0, 100 mM NaCl, 1 M hexylene glycol, 10 mM MgCl2, 0.5% Triton X-100, 10 mM 2-mercaptoethanol, and cOmplete$^{TM}$) and a final washing round with M3 buffer (10 mM sodium phosphate pH 7.0, 100 mM NaCl, 10 mM 2-mercaptoethanol, and cOmplete$^{TM}$). Washed nuclei were resuspended in sonication buffer (10 mM Tris-HCl pH 8.0, 1 mM EDTA, 0.1% SDS, and protease inhibitor). Chromatin was sheared using a Covaris E220 High Performance Focused Ultrasonicator for 12 min at 4 °C (Duty factor 5.0; PIP 140.0; Cycles per Burst: 200) in 1 ml Covaris milliTUBE (Covaris, #520128), and debris was removed by centrifugation for 5 min, 12,000*g* at 4ºC. The quality of the shearing step was assessed by controlling the length distribution of the DNA after sonication by reverse cross-linking DNA (as described below) on an aliqot of the sample (5%). DNA size distributions were analyzed on a 5200 Fragment Analyser System (Agilent, #M5310AA). A good shearing should yield DNA fragment of 150 to 300bp (Supplementary Figure 23). With the remaining sample, chromatin fragments were diluted 3X by adding 2 vol. of ChIP dilution buffer (16.7 mM Tris-HCl pH 8.0, 167 mM NaCl, 1.2 mM EDTA, 1.1% Triton X-100, 0.01% SDS, and cOmplete$^{TM}$). Protein A/G magnetic beads (ThermoFisher, #10002D and 10004D) (50mL of slurry per gr of tissue) were added to the sheared chromatin depending on the antibody used (protein A beads for rabbit polyclonal antibodies and protein G for mouse monoclonal ones, see below) , followed by incubation (at 4 °C for 1 hr with rotation). Pre-cleared samples were then incubated overnight at 4°C overnight under rotation with 5 µg of specific antibodies: anti-H3 (rabbit polyclonal, Abcam, #ab1791), anti-H3K27me3 (rabbit polyclonal, Millipore, #07–449), anti-H3K4me3 (rabbit polyclonal, Abcam, #ab8580), anti-H3K9me1 (rabbit polyclonal, Abcam, 3ab8896), anti-H3K9me2 (mouse monoclonal, Abcam, #ab1220) or or anti-H3K27me1 (rabbit polyclonal, Merck-Millipore, #17–643). The following day, the samples were mixed and incubated with 30 µl of protein A/G magnetic beads (at 4 °C for 3 hr. with rotation), followed by several rounds of washing as follows: 2 rounds of low salt buffer (20 mM Tris-HCl pH 8.0, 150 mM NaCl, 2 mM EDTA, 1% Triton X-100, and 0.1% SDS), one of high salt buffer (20 mM Tris-HCl pH 8.0, 500 mM NaCl, 2 mM EDTA, 1% Triton X-100, and 0.1% SDS), one of LiCl buffer (10 mM Tris-HCl pH 8.0, 1 mM EDTA, 0.25 M LiCl, 1% IGEPAL CA-630, and 0.1% deoxycholic acid), and one final of TE buffer (10 mM Tris-HCl pH 8.0 and 1 mM EDTA). Immunoprecipitated DNA was eluted by adding 500 µl of elution buffer (1% SDS and 0.1 M $NaHCO_3$), followed by an incubation at 65°C for 15 minutes. Reverse crosslink was carried using 51 µl of reverse crosslink buffer (40 mM Tris-HCl pH 8.0, 0.2 M NaCl, 10 mM EDTA, 0.04 mg/ml proteinase K (ThermoFisher, #EO0491), incubated for 3 hr at 45°C followed by 16 hr at 65°C. Reverse crosslinked DNA was treated with 10 µg of RNaseA (ThermoFisher #EN0531) for 30min at RT , and purified with a MinElute PCR purification kit (Qiagen, #28004).

**Staining of nuclei:**

*Arabidopsis* tissue was obtained from the second youngest leaf of a 3-week-old rosette grown in soil. The leaf was cut and placed on a previously chilled (4ºC) sterile wet gauze and immediately processed. *Spirodela* fronds were obtained from cultures as described above. Tissue was processed according to (Galbraith et al. 1983) using a sharp razor blade (Astra™ superior platinum double edge, P&G) in 500µl of modified Otto's buffer I: 0.1M citric acid (PanReac AppliChem, #131808.121), 0.5% Triton-x-100 (PanReac AppliChem, #A1388.1000), pH 1.5 (Otto et al. 1981). After filtering through a 35µm cell strainer (Falcon™, #352235), 10µl of the nuclei solution were spread onto a glass slide, let dry and mounted with 10µl of DAPI-containing

Vectashield Antifade Mounting Medium (Vector Laboratories, #H-1200). Images of the DAPI-stained nuclei were taken using a Zeiss Axio Imager.Z2 with a sCMOS camera.

**AGO4 cloning:**

Genomic DNA from *Spirodela* and *Arabidopsis* was purified using Dneasy® Plant mini kit (Qiagen,#69104) following the manufacturer's instructions and RNA isolated using TRIzol™ (ThermoFisher, #15596018) as described elsewhere. For cDNA cloning, 1µg of total RNA was treated with DNase I (Themofisher, #EN0521) and cDNA was synthesized using Maxima® First Strand cDNA Synthesis Kit (Thermo Scientific, #K1641) according to manufacturer's instructions. Amplification of all fragments was performed using 100ng of genomic DNA or cDNA as template with Phusion™ High-Fidelity DNA Polymerase (Thermo Scientific,#531S) and the primers described in Supplemental material.

A plasmid for the expression in plants of proteins tagged at the N-terminus with Flag and HA under the CaMV35s promoter was produced (35spromoter-Flag-HA-35sTerm). First by introducing a DNA fragment with the Flag sequence into a binary vector plasmid containing the 35S promoter used as backbone (Addgene #167122) through BamHI and HindIII (Thermo Scientific, #FD0054 and #FD0505) digestion and assembled using Gibson Assembly® Master Mix (NewEngland Biolabs, #E2611). The resulting plasmid (CaMV35Sp-Flag) and a DNA fragment containing the HA tag and CaMV35s terminator separated by XbaI sites, were further combined using BamHI and SmaI (Thermo Scientific, #FD0054 and #FD0663) and ligated with T4 DNA ligase (Thermo Scientific, #EL0021). The XbaI sites were introduced by PCR in At*AGO4* gDNA and Sp*AGO4a* gDNA, cloned into the plasmid described above (35spromoter-Flag-HA-35sTerm), through XbaI (Thermo Scientific, #FD0685) digestion sites using Gibson Assembly. The PCR-amplified Sp*Ago4a* cDNA was cloned into the same plasmid by digesting the PCR fragment with introduced BcuI sites in the extremities (Thermo Scientific, #FD1254), compatible with XbaI sites present in the plasmid, through T4 DNA ligase. All primers used for cloning can be found within **Supplemental Fig. S39.** All plasmids were transformed into chemically competent *E.coli* (DH5α), confirmed by SANGER sequencing, and deposit in Addgene with the following numbers: p*35S*:*FHA*-At*AGO4*_gDNA (#216838), p*35S*:*FHA*-Sp*AGO4a*_gDNA (216841), p*35S*::*FHA*-Sp*AGO4a*_cDNA (#216842).

**Splicing analysis by RT-PCR:**

RNA and cDNA were produced as described above. Oligonucleotides were designed to amplify two different regions containing several introns in Sp*AGO4a*. All fragments were amplified by PCR using 100mg of cDNA or 10ng of plasmid, as a template, and the primers described in Supplemental material (**Supplemental Fig. S39**). The resulting fragments were resolved in a 1X TBE 1% agarose gel, stained with peqGreen (Peqlab, #732-3196) and imaged using the Molecular Imager® Gel Doc™ XR System (Bio-Rad).

***Agrobacterium*-mediated transient expression:**

***N. benthamiana:*** Agrobacterium-mediated transient expression in *N. benthamiana* was performed as in (Felippes and Weigel 2010). In brief, *A. tumefaciens* strain GV3101 harbouring the vectors described below were cultured at 28°C in LB medium with appropriate antibiotics. After 36 hours, cultures were pelleted at 2000g, 10 minutes at 20°C and resuspended to an $OD_{600}$ of 0.5 in filtered sterilized agroinfiltration buffer: 2mM MES, 50mM $NaH_2PO_4$, 0.5% (w/v) glucose, pH 5.6. Acetosyringone was added prior to use at 100 µM. After 1 hour of incubation at room temperature in darkness, the bacterial solutions were infiltrated into the abaxial side of young *N. benthamiana* true leaves, using a 1ml syringe without needle. At 3 or 5 days after infiltration (dpi), leaf discs were collected, flash-frozen in liquid nitrogen and ground in liquid nitrogen using morter and pestle. The ground samples were stored at -80C.

***Spirodela polyrhiza***: For Agrobacterium-mediated transient expression in *Spirodela*, *Agrobacterium* strain EHA105 was used. Bacteria cultures were prepared as for *N. benthamiana* infiltrations with the following modifications: Bacteria was resuspended to $OD_{600}$ 0.8 in *Spirodela* agroinfiltration media (10 mM MgCl2, 5% sucrose (w/v), pH 5.6). *Spirodela* fronds were prepared as it follows: Cultures were initiated from axenic stocks by inoculating fronds into 2L glass beakers containing 0.5-1L of ½ SH media, covered with the lid of a 145 x 20 mm polystyrene culture dish (Greiner Bio-One #639102) sealed with 2.5 cm wide Leucopore tape (Duchefa Biochemie #L3301). Cultures were grown to 50-80% confluence (as surface occupancy). Prior to agroinfiltration, fronds were pretreated as in (Yang et al. 2018) by transferring them to an ice-cold 20 mg/L L-Glutamine (PanReac AppliChem #A1420) solution for 20 min. Following pretreatment, fronds were infiltrated with the *Vir*-induced Agrobacterium solution either (i) manually, at the budding pouch area with a 1 mL syringe without needle by gently applying pressure with the plunger on the upper side of fronds or, (ii) trough vacuum infiltration by transferring the fronds to 100 mL agrobacterium solution complemented with 0.02% (v/v) Silwet L-77 (Kurt-Obermieier GmbH #7060-10) in a 200 mL beaker in a vacuum desiccator. -70 kPa vacuum was applied twice for 10 min. Following either manual or vacuum agroinfiltration, excess of bacteria was removed by rinsing the fronds in $ddH_2O$ and plants transferred to 100 x 20 mm culture dishes (Greiner Bio-One #664160) containing 30-50 mL ½ SH media + 100 µM acetosyringone for a 3 day co-culture. 4 days post-infiltration (dpi) transfer fronds to plates containing ½ SH media. At 7 dpi agrobacterium elimination was initiated by or replacing the media with ½ SH + 250 µM ticarcillin (Duchefa #T0190.0010, lab stock at 250 mg/mL in H2O) + 300 µM cefotaxime (Duchefa #C0111.0005, lab stock at 300 mg/mL in H2O) for a week. Plants were cultured with regular (1/week) ½ SH media replacements. See Supplementary Fig. S36 for a graphic depiction of the protocol and workflow.

**Immunoprecipitation of Flag-tagged AGO4:**

100-150mg of *N. benthamiana* ground tissue was homogenized in 1 ml of IP buffer (50 mM Tris-HCl, pH 7.5, 150 mM NaCl, 10% glycerol, 0.1% NP-40) containing 2 µM MG-132 and cOmplete$^{TM}$ mini EDTA-free protease inhibitor cocktail (Roche, #04693159001) for 30 minutes at 4 °C. After lysis, debris were removed by centrifugation (12000*g* for 10 min at 4ºC). An aliquot of lysate was kept as input, while the remainder was incubated with ANTI-FLAG$^{R}$ M2 Magnetic beads (20µl of slurry/sample) (Sigma-Aldrich, #M8823) for 3 hours at 4°C, on a orbital shaker. After incubation, the resin was separated from supernatant in a magnetic rack (DynaMag-5, Thermofisher, #12321D) and washed with 1 ml of IP buffer for 20 minutes at 4°C, for 3 times. After the final wash, 100µl of PBS were added and 10µl put aside, used to check immunoprecipitation quality by western blot by adding Laemli buffer and boiling the sample for 10 min. at 95ºC. The remaining 90µl was used to analyze the small RNA composition upon extraction using TRIzol$^{TM}$ (ThermoFisher, #15596018). The isolated small RNAs were used to generate libraries and sequenced as described above. TraPR purified small RNAs (described above) from not-infiltrated plants were used to asses *N. benthamiana* small RNA profile.

**Small RNA blot analysis:**

AGO-loaded small RNAs were purified from 100mg of starting material using TraPR columns, as described above. The resulting small RNAs were mixed with an equal volume of 2X Novex™ TBE-Urea sample buffer (ThermoFisher, #LLC6876), separated on a 17.5% polyacrylamide-urea gel, transferred onto Hybond-NX membrane (Cytiva, #RPN303T) and chemically crosslinked (12.5M 1-methylimidazol (Merk, #M50834), 31.25mg/mL N-(3-dimethylaminopropyl)-N'-ethylcarbodiimide hydrochloride (Merk, #E7750) as in (Pall and Hamilton 2008). Probes produced from PCR templates were radiolabelled with [α-$^{32}$P]-dCTP (Hartmann Analytic, #SRP-305) using Prime-it II Random Primer Labelling kit (Agilent, #300385) and oligonucleotides used as a probe were radiolabelled with [$\gamma$-$^{32}$P]-ATP (Hartmann analytic #SRP-501), using T4 Polynucleotide kinase (Thermofisher, #EK0031) (see Supplemntary figures for a list of oligonucleotides used). The probes were incubated in PerfectHyb™ Plus Hybridization Buffer (Sigma Aldrich, #H7033) and washed with 2XSSC, 0.1%SDS. The signal was detected with the Amersham™Typhoon™ (Cytivia). Probes and oligonucleotides used are listed in **Supplemental Fig. S39**.

**Small RNA extraction:**

**Total RNA:** For small RNA extraction from total RNA and TraPR input fraction, 10µg of total RNA (as described above) or RNA extracted as described above from TraPR inputs was separated on a 15% denaturing polyacrylamide-urea gel. For size reference, 20µl of microRNA marker (New Englad Biolabs, #N2102S) were loaded in the same gel. Following gel staining with peqGreen (Peqlab, #732-3196), and using the marker as reference, small RNAs ranging from 17-to-25 nt were excised and recovered using the ZR small-RNA PAGE Recovery Kit (Zymo Research, #R1070).

**TraPR:** Purifucation of Argonaute-loaded small RNAs was performed according to the protocol described in (Grentzinger et al. 2020). In brief, 100mg of flash-frozen plant tissue was ground to fine powder using a Silamat S6 (Ivoclar Vivadent) twice for 3s in 1.5 mL microcentrifuge tubes with ~20 1 mm ø glass beads (Carl Roth, #A554.1). Frozen powder was incubated with TraPR lysis buffer and the homogenate loaded into the TraPR columns. TraPR eluates were collected and small RNAs extracted by adding and mixing 1 volume of acidic Phenol:Chloroform:Isoamyl alcohol (50:49:1, pH 4.5-5) (Carl Roth, #X985.1). After centrifugation (12000*g* for 10 min at 4ºC) RNA was precipitated from the aqueous phase with 1 volume of isopropanol and 10% (v/v) of sodium actetate 3M pH 5.2, overnight at -20°C. RNA was collected by centrifugation (12000*g* for 30 min at 4ºC) and the resulting pellet was washed with 75% ethanol and resuspeneded in 10-20 µL RNase free water.

#### Supplementary References to Supplementary Methods

Camacho C, Coulouris G, Avagyan V, Ma N, Papadopoulos J, Bealer K, Madden TL. 2009. BLAST+: architecture and applications. *BMC Bioinform* **10**: 421.

Consortium TU, Bateman A, Martin M-J, Orchard S, Magrane M, Ahmad S, Alpi E, Bowler-Barnett EH, Britto R, Bye-A-Jee H, et al. 2022. UniProt: the Universal Protein Knowledgebase in 2023. *Nucleic Acids Res* **51**: D523–D531.

Doblmann J, Dusberger F, Imre R, Hudecz O, Stanek F, Mechtler K, Dürnberger G. 2019. apQuant: Accurate Label-Free Quantification by Quality Filtering. *J Proteome Res* **18**: 535–541.

Dorfer V, Pichler P, Stranzl T, Stadlmann J, Taus T, Winkler S, Mechtler K. 2014. MS Amanda, a Universal Identification Algorithm Optimized for High Accuracy Tandem Mass Spectra. *J Proteome Res* **13**: 3679–3684.

Felippes FF de, Weigel D. 2010. Transient assays for the analysis of miRNA processing and function. *Methods in molecular biology (Clifton, NJ)* **592**: 255–264.

Galbraith DW, Harkins KR, Maddox JM, Ayres NM, Sharma DP, Firoozabady E. 1983. Rapid Flow Cytometric Analysis of the Cell Cycle in Intact Plant Tissues. *Science* **220**: 1049–1051.

Grentzinger T, Oberlin S, Schott G, Handler D, Svozil J, Barragan-Borrero V, Humbert A, Duharcourt S, Brennecke J, Voinnet O. 2020. A universal method for the rapid isolation of all known classes of functional silencing small RNAs. *Nucleic Acids Res* **48**: gkaa472-.

Käll L, Canterbury JD, Weston J, Noble WS, MacCoss MJ. 2007. Semi-supervised learning for peptide identification from shotgun proteomics datasets. *Nat Methods* **4**: 923–925.

Kanno T, Huettel B, Mette MF, Aufsatz W, Jaligot E, Daxinger L, Kreil DP, Matzke M, Matzke AJM. 2005. Atypical RNA polymerase subunits required for RNA-directed DNA methylation. *Nature Genetics* **37**: 761–765.

Mistry J, Chuguransky S, Williams L, Qureshi M, Salazar GA, Sonnhammer ELL, Tosatto SCE, Paladin L, Raj S, Richardson LJ, et al. 2020. Pfam: The protein families database in 2021. *Nucleic Acids Res* **49**: gkaa913-.

Otto FJ, Oldiges H, Göhde W, Jain VK. 1981. Flow cytometric measurement of nuclear DNA content variations as a potential in vivo mutagenicity test. *Cytometry* **2**: 189–191.

Pall GS, Hamilton AJ. 2008. Improved northern blot method for enhanced detection of small RNA. *Nature Protocols* **3**: 1077–1084.

Paysan-Lafosse T, Blum M, Chuguransky S, Grego T, Pinto BL, Salazar GA, Bileschi ML, Bork P, Bridge A, Colwell L, et al. 2022. InterPro in 2022. *Nucleic Acids Res* **51**: D418–D427.

Perez-Riverol Y, Bai J, Bandla C, García-Seisdedos D, Hewapathirana S, Kamatchinathan S, Kundu DJ, Prakash A, Frericks-Zipper A, Eisenacher M, et al. 2021. The PRIDE database resources in 2022: a hub for mass spectrometry-based proteomics evidences. *Nucleic Acids Res* **50**: D543–D552.

Rappsilber J, Mann M, Ishihama Y. 2007. Protocol for micro-purification, enrichment, pre-fractionation and storage of peptides for proteomics using StageTips. *Nat Protoc* **2**: 1896–1906.

Sambrook J, Russell DW. 2001. *Molecular Cloning*. CSHL Press.

Scheid R, Dowell JA, Sanders D, Jiang J, Denu JM, Zhong X. 2022. Histone Acid Extraction and High Throughput Mass Spectrometry to Profile Histone Modifications in Arabidopsis thaliana. *Curr Protoc* **2**: e527.

Schwach F, Vaistij FE, Jones L, Baulcombe DC. 2005. An RNA-Dependent RNA Polymerase Prevents Meristem Invasion by Potato Virus X and Is Required for the Activity But Not the Production of a Systemic Silencing Signal. *Plant Physiol*.

Schwanhäusser B, Busse D, Li N, Dittmar G, Schuchhardt J, Wolf J, Chen W, Selbach M. 2011. Global quantification of mammalian gene expression control. *Nature* **473**: 337–342.

Taus T, Köcher T, Pichler P, Paschke C, Schmidt A, Henrich C, Mechtler K. 2011. Universal and Confident Phosphorylation Site Localization Using phosphoRS. *J Proteome Res* **10**: 5354–5362.

Yang G-L, Fang Y, Xu Y-L, Tan L, Li Q, Liu Y, Lai F, Jin Y-L, Du A-P, He K-Z, et al. 2018. Frond transformation system mediated by Agrobacterium tumefaciens for Lemna minor. *Plant Mol Biol* **98**: 319–331.
